# freqpcr: estimation of population allele frequency using qPCR ΔΔCq measures from bulk samples

**DOI:** 10.1101/2021.01.19.427228

**Authors:** Masaaki Sudo, Masahiro Osakabe

## Abstract

PCR techniques, both quantitative (qPCR) and non-quantitative, have been used to estimate allele frequency in a population. However, the labor required to sample numerous individuals, and subsequently handle each sample, makes quantification of rare mutations, including pesticide resistance genes at the early stages of resistance development, challenging. Meanwhile, pooling DNA from multiple individuals as a “bulk sample” may reduce handling costs. The qPCR output for a bulk sample, however, contains uncertainty owing to variations in DNA yields from each individual, in addition to measurement errors. In this study, we developed a statistical model for the interval estimation of allele frequency using ΔΔCq-based qPCR analyses of multiple bulk samples collected from a population. We assumed a gamma distribution as the individual DNA yield and developed an R package for parameter estimation, which was verified with real DNA samples from acaricide-resistant spider mites, as well as a numerical simulation. Our model resulted in unbiased point estimates of the allele frequency compared with simple averaging of the ΔΔCq values, while their confidence intervals suggest that collecting and pooling additional samples from individuals may produce higher precision than individual PCR tests with moderate sample sizes.

## Introduction

Estimating the frequency of specific alleles in populations is a key technique not only in population genetics and molecular ecology, but also in agricultural and regulatory sciences (Falconer, 1960; Kim et al., 2011; Yamamura & Hino, 2007). In applied entomology, field monitoring has been performed to detect resistance genes of arthropod pests to pesticides and genetically modified insecticidal plants, such as *Bt* crops (Andow & Alstad, 1998; Sonoda et al., 2017).

Entomologists have traditionally estimated resistance allele frequencies via bioassays (Gould et al., 1997; Li et al., 2016; Tabashnik et al., 2000), in which insects directly collected from fields, or their offspring reared in laboratories, are exposed to chemical compounds of interest to obtain measurements, such as mortality rate. However, bioassays associated with the treatment of living organisms have certain inherent drawbacks. Specifically, they are often labor-intensive and time-consuming. Although the resistance level can be directly measured using bioassays that detect the mortality of tested individuals, additional information, including the dominance of the resistance gene, is required to estimate allele frequency.

In accordance with the development of genome-wide association studies on resistance genes (ffrench-Constant, 2013; Snoeck et al., 2019; Sugimoto et al., 2020), rapid advancements have recently been made in molecular diagnostics (Donnelly et al., 2016; Samayoa, et al., 2015; Toda et al., 2017). To quantify resistance-associated point mutations at the population scale, the most fundamental molecular technique is an individual-based polymerase chain reaction (PCR) analysis (Toda et al., 2017). Moreover, quantitative PCR (qPCR), based on real-time PCR, is also used for the point mutation of allele frequencies (Germer et al., 2000). If the alleles are distributed randomly in the target population, a simple binomial assumption enables us to estimate the population allele frequency and its confidence interval. However, collecting, processing, and analyzing multiple DNA samples may not feasible, particularly when dealing with numerous samples from multiple sites, or when estimation of a rare (<1%) mutation frequency is required for a given population, as is often the case in the early phase of resistance development.

Although rearing living insects is no longer necessary, the field of molecular diagnostics is still lacking a metaphorical silver bullet capable of reducing the required time and cost associated with handling multiple samples, while guaranteeing estimation precision and accuracy. The use of a “bulk sample” (i.e., pooling multiple individual samples and processing a single DNA extract), in coordination with statistical methods, such as group testing, may address some of these challenges. In fact, Osakabe et *al*. (2017) and Maeoka et *al*. (2020) developed diagnostic methods for detecting resistance to the acaricide, etoxazole, in the two-spotted spider mite, *Tetranychus urticae* Koch (Acari: Tetranychidae), which is conferred by an amino acid substitution in chitin synthase 1 (*CHS1*; I1017F) (Van Leeuwen et al., 2010). They used a bulk sample to measure the frequency of the resistant point mutation in field mite populations. To calculate the point estimate, these studies compared the relative quantity of the resistance allele with an internal reference (housekeeping gene) in the sample, known as the ΔΔCq method (Livak and Schmittgen, 2001). In the etoxazole-R diagnosis by Osakabe et al. (2017), glyceraldehyde-3-phosphate dehydrogenase (*GAPDH*) was used as the housekeeping gene.

In this study, we propose a statistical method for obtaining the interval estimate of allele frequency using ΔΔCq-based qPCR analyses forr multiple bulk samples taken from a population. We first introduced the random error structure to approximate the relative abundance of the two alleles (wild type and mutant) and their ratios in the bulk DNA sample. Thereafter, we formulated how the relative amounts of the two alleles in a sample solution impacted the Cq measurements through ΔΔCq-based qPCR analysis. Finally, we combined the models and developed a maximum likelihood estimation procedure to estimate an allele frequency implemented using the R language. The package source is available on the Internet (https://github.com/sudoms/freqpcr).

## Model

### Approximation of allele quantities contained in a bulk DNA sample

When DNA is directly extracted from the whole body of a living organism, the DNA yield is roughly proportional to its body weight (Chen et al., 2010). For insects, the intra-population frequency distribution of body weight is often approximated using a unimodal and right-skewed continuous distribution, typically lognormal or gamma distribution (May, 1976; Rakovski et al., 2011; Knapp, 2016). In fact, it has been suggested that body weights are distributed lognormally in many non-social insect species (Gouws et al., 2011).

In this study, we adopted a gamma, rather than lognormal, distribution to approximate the DNA amount per individual organism for two reasons. First, it is difficult to distinguish which distribution a real population obeys when the sample size is small. The two distributions are considered interchangeable (Wiens, 1999; Kundu and Manglick, 2005). Second, the sum and proportion of independent gamma distributions have closed forms under certain conditions. Using Eq. 1, let *X* (*X* ≥ 0) be the DNA yield per single locus per individual:

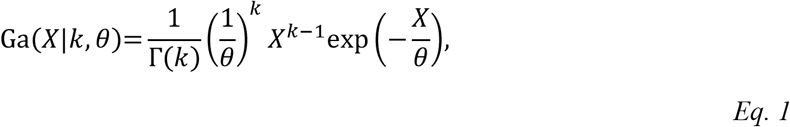

where Γ(.) denotes the gamma function. The parameters *k* and *θ* (*k, θ* > 0) are the shape and scale parameters of the gamma distribution, respectively. The mean is given by *kθ*.

Using Eq. 1, let us consider the amounts of allelic DNA in the sample extracted from multiple individuals at once, hereafter referred to as “a bulk sample.” Table 1 lists the variables and parameters of the model structure. For simplicity, we model the case of haploidy in the main text, while Appendix S1 describes the approximated formulation for diploids. Now, we have *n* insects, of which *m* (*m* = 0,1,,…, *n*) are the genotypes resistant to an insecticide (hereafter denoted by R). The rest *n — m* carried S, the susceptible allele. When we capture insects from a wild population, the size of *n* is obvious, however *m* is usually unknown (Figure 1A). Assuming random sampling from an infinite population with the R allele at the frequency *p, m* follows a binomial distribution (Eq. 2):

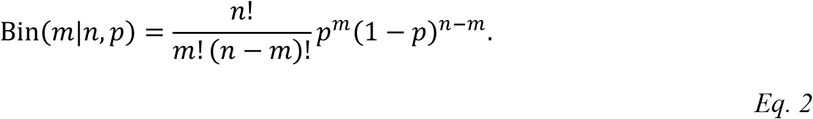

**Table 1.**
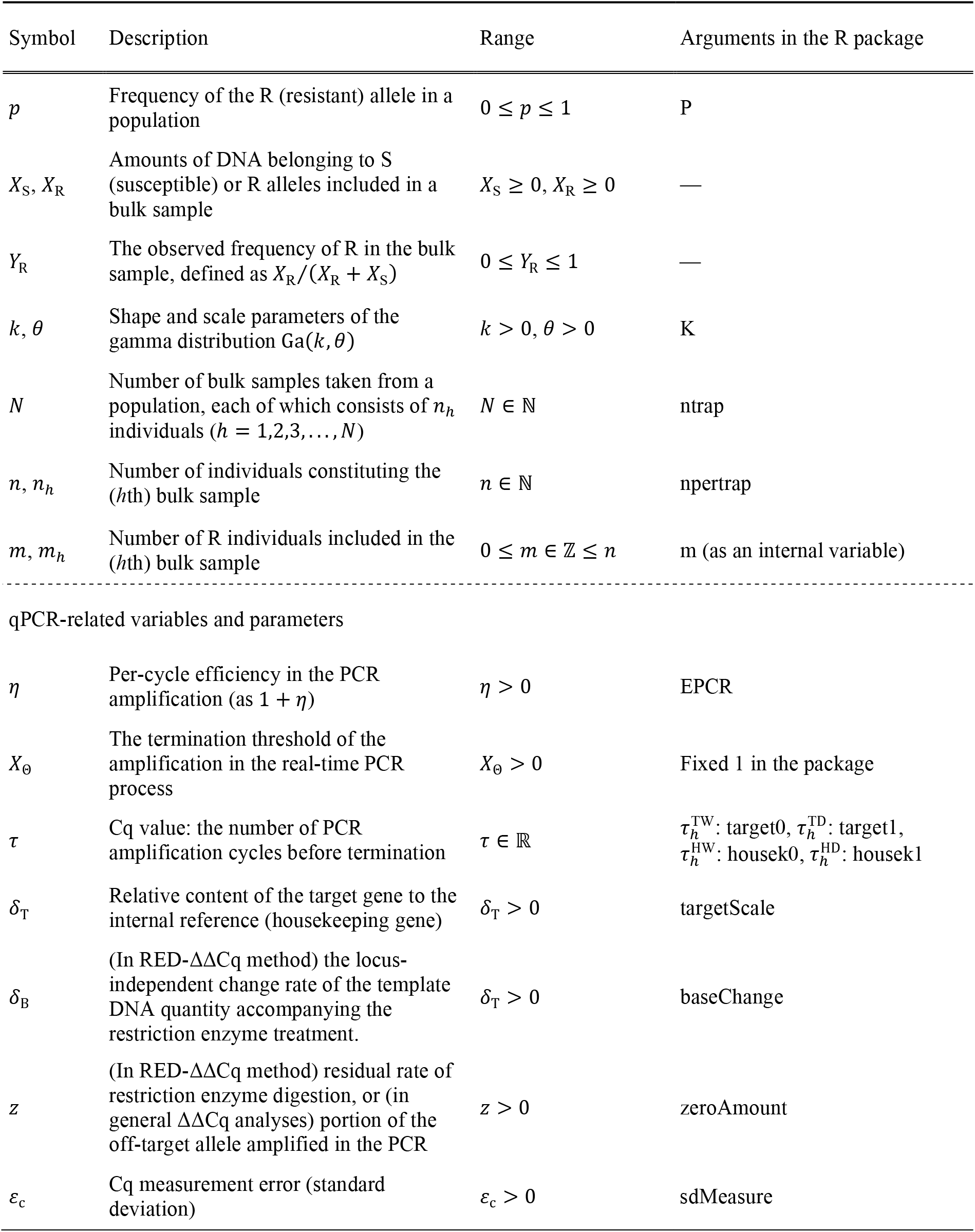
Description of variables and parameters

**Figure 1.**
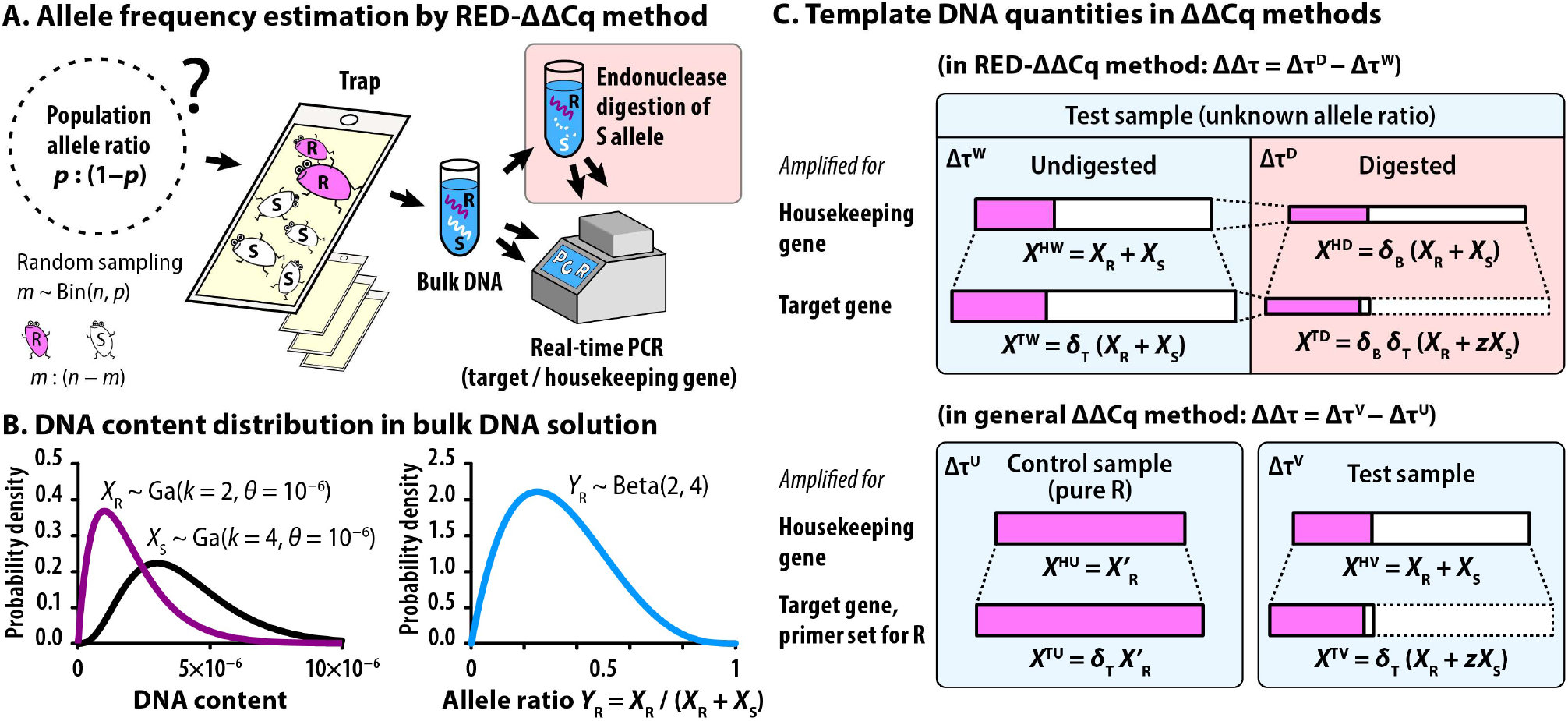
Scheme of population allele frequency estimation based on qPCR analyses. A: Insect sampling and subsequent qPCR analysis using the restriction enzyme digestion (RED)-ΔΔCq method. B: Probability distributions of the DNA amounts of the resistant (R) and susceptible (S) alleles and their ratio in the bulk sample. C: Template DNA involved in the RED-ΔΔCq analysis and a general ΔΔCq analysis using an R-specific primer set. In either method, the frequency of *X*_R_ in a test sample is quantified as *X*_R_ + *zX*_S_ (≅ *X*_R_) measured on the target gene, divided by *X*_R_+ *X*_S_ measured on a housekeeping gene in the sample. As the copy numbers may differ between genes, the relative content *δ*_T_ is also quantified using a control sample.

When the bulk sample contains at least one resistant individual, 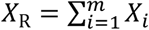 denotes the total R content. If there is no systematic error in the efficiency of DNA extraction between the genotypes, and if *X_i_*, the individual DNA yield obeys the gamma distribution of Eq. 1, then *X*_R_ follows the gamma distribution with the shape parameter *mk* and scale parameter *θ* based on the reproductive property. Conversely, the amount of S allele is denoted by 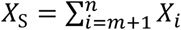, which follows the gamma distribution with (*n* — *m*)*k* and *θ* (Figure 1B).

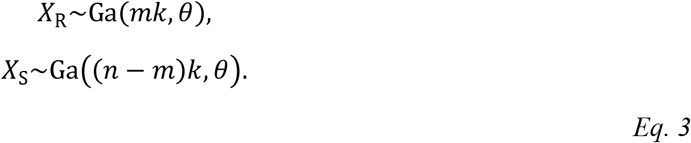

When *X*_R_ and *X*_S_ independently follow gamma distributions with the same scale parameter, the observed allele frequency *Y*_R_ = *X*_R_/(*X*_S_ + *X*_S_) follows a beta distribution with the shape parameters *mk* and (*n — m*)*k*: Eq.

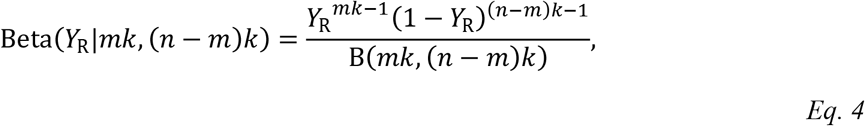

where B(.) is a beta function. This error structure was originally developed to model allele frequencies measured via quantitative sequencing (Sudo et al., in press).

### Relative quantification of DNA by real-time PCR

#### Allele frequency estimation from a single bulk sample: RED-ΔΔCq method

The ΔΔCq (quantification cycle) method (Livak, 1997) is the most common method for relative quantification using qPCR, in which the quantities of complementary cDNA libraries are compared between samples to determine the relative expression levels of the genes of interest. Osakabe *et al*. (2017) expanded this concept and proposed the “RED-ΔΔCq method” (RED, restriction enzyme digestion), a derivative method that can measure the allele frequency from a single sample solution, to diagnose the regional prevalence of an acaricide-resistant point mutation in a *T. urticae* population.

In the RED-ΔΔCq method, the control was prepared as an intact sample containing total DNA (= *X*_R_ + *X*_S_) on the target locus. The sample in question was the same DNA extract, however, was digested with restriction endonucleases prior to qPCR analysis (Figure 1A). The restriction site is designed to recognize the S allele on the target locus so that the operation digests the major part of S (denoted by 1 — *z: z* is a small yet, positive variable giving the residual rate). Consequently, we obtained the template amount *X*_R_ + *zX_S_* at the target locus after digestion. To calibrate the template DNA amounts, the samples before and after digestion were also amplified using the primer set for a housekeeping gene as an internal reference.

Taken together, the single bulk sample results in a quartet of Cq measurements differentiating at the target loci (resistance-associated and housekeeping genes) × restriction enzyme digestion (undigested and digested). We can then formulate the allele frequencies by letting *X^HW^* and *X^TW^* represent the total amounts of template DNA at the housekeeping (H) and target (T) loci, respectively, included in the sample without digestion, the state denoted by W (Figure 1C).

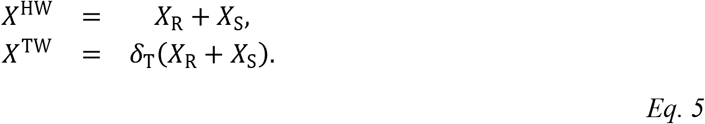

The coefficient *δ_T_* (*δ_T_* > 0) provides the relative content of the target gene to the housekeeping gene in genomic DNA (the difference in the DNA extraction efficiencies is also included). After digestion (state D), *X^HD^* and *X^TD^* denote the DNA amounts at the H and T loci, respectively:

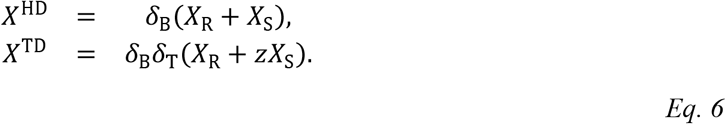

The common coefficient *δ*_B_ (*δ*_B_ > 0) provides the rate of certain locus-independent changes in the quantities of template DNA accompanying the restriction enzyme treatment.

As a result of qPCR, the Cq quartet, *τ*^HW^, *τ*^TW^, *τ*^HD^, and *τ*^TD^ were obtained as:

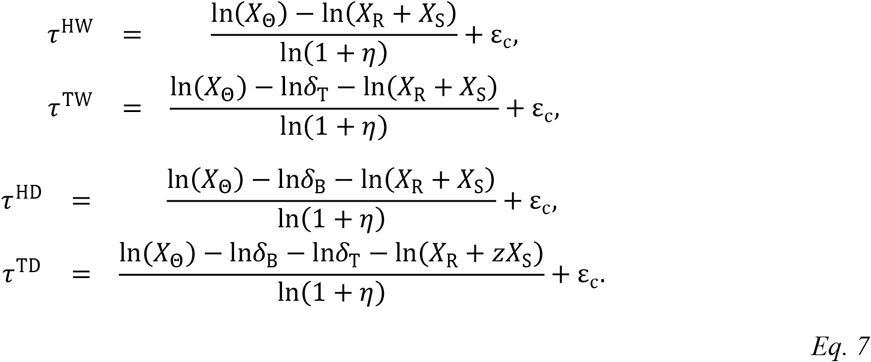

Here, 1 + *η* (*η* > 0) and *X_Θ_* denote the amplification efficiency per PCR cycle and its threshold, respectively. According to Livak and Schmittgen (2001), we assume an ideal amplification, where *X_Θ_* is set within the early exponential amplification phase. The actual Cq data contain measurement errors in addition to uncertainty due to experimental operations, such as sample dispensation or PCR amplification. We express these using the common error term 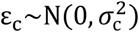, following the normal distribution of mean = 0 and 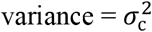 in the scale of raw Cq values. The validity of this error structure is verified later.

The two ΔCq values were then defined for the undigested and digested samples, as *Δτ^W^* = *τ^TW^* — *τ^HW^* and Δ*τ*^D^ = *τ*^TD^ — *τ*^HD^, respectively. Their ΔΔCq are:

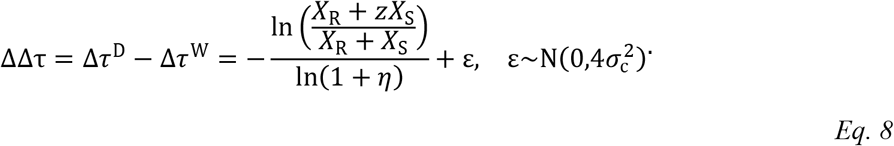

From Eq. 8, the expected value of (*X*_R_ + *zX*_S_)/(*X*_R_ + *X*_S_) is calculated as (1 + *η*)^−ΔΔτ^. The coefficients *δ*_B_ and *δ*_T_ in Eq. 5 and Eq. 6 vanished by subtracting the Cq values and ΔCq values, respectively.

The point estimate of the resistance allele frequency, *Ŷ*_R_, is defined as *X*_R_/(*X*_R_ + *X*_S_) for each bulk sample. When *z* is much smaller than *Ŷ*_R_, the quantity (*X*_R_ + *zX*_S_)/(*X*_R_ + *X*_S_ = *Ŷ*_R_ + z(1 — *Ŷ*_R_) itself can approximate the frequency, which will be the case with enough digestion time before qPCR. However, the use of the point estimate may introduce a problem in that the size of *Ŷ*_R_ often exceeds 1 when the R frequency is high and a larger error exists in the Cq measurement (also see the result of Experiment 2).

Although the value of 1 + *η* may vary on the primer sets, both target and housekeeping loci share the same amplification efficiency in Eq. 7, because practical PCR protocols were designed to be 1 + *η* ≅ 2. We can also approximately cancel the effect of heterogeneous amplification efficiencies by fitting the *δ*_T_ size of the sample sets with known allele ratios (Experiment 1).

#### Measurement of ΔΔCq using allele-specific primer sets

While the RED-ΔΔCq method enabled us to measure allele frequency from the bulk sample, enzyme availability is a prerequisite to digest the S-allele-specific restriction site at the target locus. A longer digestion period (3 h) was also required to quantify etoxazole resistance in the protocol by Osakabe *et al*. (2017).

Maeoka *et al*. (2020) demonstrated that a general ΔΔCq method without restriction enzyme treatment could be used for allele-frequency measurement if a specific primer set were to be designed to amplify only the R allele at the target locus. Similar to the RED-ΔΔCq method, DNA samples with unknown mixing ratios were dispensed and amplified using primer sets corresponding to T and H loci, respectively. Unlike the RED-ΔΔCq method, the control sample was not taken from the test sample solution, but rather was prepared as a DNA solution containing 100% R, hereafter denoted as U (= pUre R line) (Figure 1C).

X^HU^ and X^TU^ then denote the template DNA quantities:

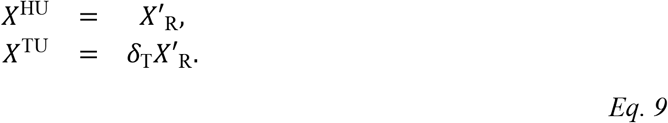

Though the definition of *δ*_T_ is the same as Eq. 5, the quantity is denoted by *X’*_R_ instead of *X*_S_ + *X*_R_ as it no longer originates from the R portion of the test sample itself (i.e., not internal).

For the test sample (denoted as V), the template DNA quantities amplified at the housekeeping (*X*^HV^) and target (*X*^TV^) loci are expressed as follows:

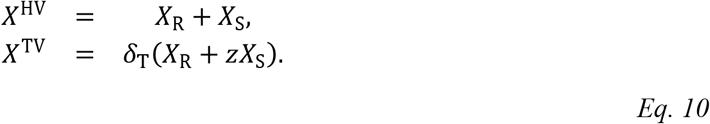

In the PCR process of the modified ΔΔCq method, the small positive number *z* provides the template quantity of S, which is non-specifically amplified even with the R-specific primer set. As the primer set for the housekeeping gene was nonspecific, *X*^HV^ was fully amplified. Assuming that all four template DNAs are amplified with efficiency 1 + *η*, we define the two ΔCq values as Δ*τ*^U^ = *τ*^TU^ — *τ*^HU^ and Δ*τ*^V^ = *τ*^TV^ — *τ*^HV^. Finally, their ΔΔCq values are ΔΔ*τ* = Δ*τ*^V^ — Δ*τ*^U^, which yields a formula identical to Eq. 8.

### Interval estimation of allele frequency and experimental parameters based on qPCR over multiple bulk samples

Finally, we consider the likelihood model to obtain the interval estimate of the allele frequency based on the (RED-)ΔΔCq analysis over multiple bulk samples. Assume that the population has the R allele at the frequency *p* from which *N* bulk samples are taken. The *h*th sample (*h* = 1,2,3,…, *N*) consists of *n_h_* haploid individuals, of which *m_h_* are resistant mutants. As shown in Eq. 7, the Cq values (denoted as 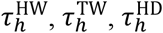, and 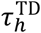 for each bulk sample) are determined not only by the DNA quantities, denoted as *X_h,R_* and *X_h,S_*, but also by parameters such as *δ*_T_ or 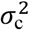 accompanying the experimental operation. We can simultaneously estimate these if we have multiple bulk samples, for which the likelihood function of obtaining the Cq values under the parameters is defined.

We propose the joint likelihood for the two ΔCq values, 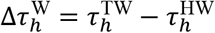 and 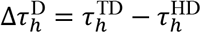, for the convenience of numerical calculation:

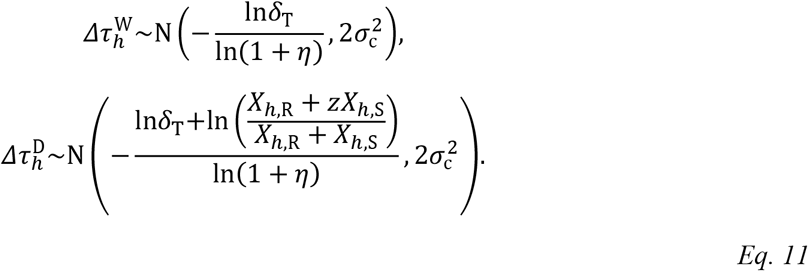

Although Eq. 11 is defined for the RED-ΔΔCq method, it is also applicable to the ΔΔCq method by Maeoka *et al*. (2020) by substituting 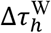 and 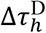 to 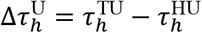 and 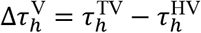, respectively.

#### Formulation of likelihood based on gamma or beta distribution

Using the relationship between *m_h_*, *n_h_*, and *p* in Eq. 2, we proceed to the likelihood function defined as the probability of observing the set of 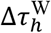 and 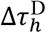 under the given values of *p, n_h_*, and other experimental parameters. In Eq. 11, 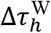 is not affected by the R: S ratio in the bulk sample; it is only affected by the experimental parameters, *δ*_T_, *η*, and 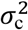. In addition, by taking the differences, there is no need to estimate as *X*_Θ_ and *δ*_B_ appear in Eq. 7. Moreover, cancelation of *δ*_B_ also ensures that we can apply the model of Eq. 11 to the general ΔΔCq method of Eq. 9 and Eq. 10.

Conversely, we must consider the amount of DNA in the bulk sample to calculate the probability of obtaining 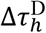. When the size of *m_h_* is specified under the binomial assumption, the quantities of DNA in the *h*th bulk sample, *X*_*h*,R|*m_h_*_ and *X*_h__S_|*m_h_*, can independently take any positive values following the gamma distribution of Eq. 3, and their proportions *X*_*h*,R|*m_h_*_ =*X*_*h*,R|*m_h_*_ / (*X*_*h*,R|*m_h_*_ + *X*_h__S_|*m_h_*) are Beta(*m_h_k*, (*n_h_* — *m_h_*)*k*) as shown in Eq. 4. If the sample contains only S or R, then 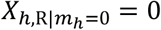 or 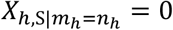 is guaranteed.

The likelihood function for the observed ΔCq values on the *h*th bulk sample *L_h_* is defined as follows:

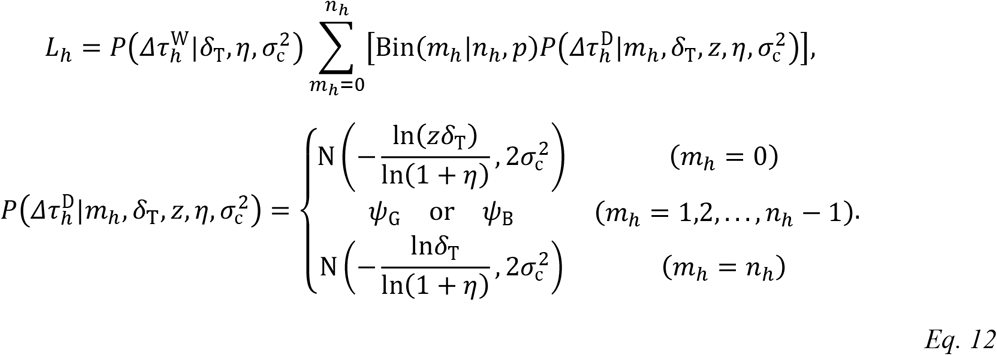

In Eq. 12, *Ψ*_G_ or *Ψ*_B_ denotes the probability of obtaining 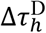 under the template DNA quantities of 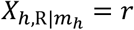 and 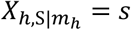 if we formularize the two quantities by gamma distribution, or if we formularize their mixing ratio by the single beta distribution, respectively. We must consider not only the possible cases of *m_h_*, but also the entire range of the DNA amounts. If we use the gamma distributions, for every case *m_h_* = 1,2,…, *n_h_* — 1, we need to calculate the double integration for *Ψ*_G_ under the whole region of *X*_*h*,R|*m_h_*_ = *r* and *X*_*h*,S|*m_h_*_ = *S* for the interval {*D*: 0 ≤ *r* < ∞, 0 ≤ *s* < ∞}.

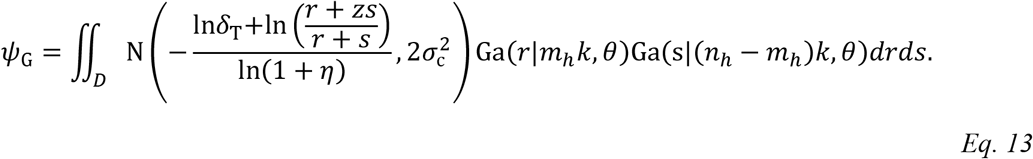

The common scale parameter of the gamma distributions, *θ*, is not identifiable from the data, although we can substitute arbitrary values *θ* = 1 for it because it is canceled in ln[(*r* + *zs*)/(*r* + *s*)] in Eq. 13.

Since the computational burden for the double integration is large, we simplified the likelihood model with the beta distribution. By introducing *y* = *r*/(*r* + *s*), the probability of obtaining 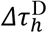 is replaced with *ψ*_B_ defined as follows:

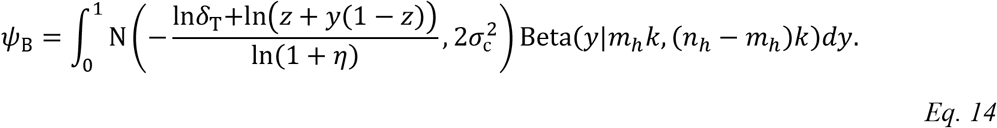

We provide an R function “freqpcr()” to estimate the parameters *p, k, δ*_T_, and *σ*_c_ simultaneously when the set of Cq measurements (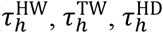, and 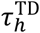) and *n_h_* are given for each of the *N* bulk samples. The default is freqpcr(…, beta = TRUE), where the beta distribution model of Eq. 14 was used instead of gamma. Regardless of the algorithms, the asymptotic confidence intervals are calculated using the inverse of the Hessian matrix evaluated at the last iteration. The functions nlm() of R and cubintegrate() in the R package “cubature” (Narasimhan et al., 2019) are used for the iterative optimization and the (double) integration, respectively.

### Identification of auxiliary parameters using DNA samples with known allele-mixing ratios

The likelihood introduced above ensures that we can estimate the sizes of p and *k* together with other experimental parameters if we have conducted a (RED-)ΔΔCq analysis on multiple bulk samples. However, the size of *z*, the residue rate of the S allele, is not identified and must be specified as a fixed parameter. The amplification efficiency, p, is estimated in theory over the iterative calculation of Eq. 11, but in fact, simultaneous estimation sometimes fails when p is set as unknown.

Therefore, the experimenter should identify the sizes of these auxiliary parameters. To estimate their plausible sizes, one can conduct (RED-)ΔΔCq analysis using DNA solutions with known allele ratios; for instance, DNA can be extracted from each of the pure breeding lines of S and R and mix the solutions at multiple ratios, or make a dilution series of R by S. As the ratio of *X*_R_ to *X*_S_ is strictly fixed, Eq. 7 is directly applicable to express the relationship between DNA quantities and the four Cq measurements. The R functions knownqpcr() and knownqpcr_unpaired() appearing in the package provide the maximum likelihood estimation for *δ*_B_, *δ*_T_, *σ*_c_, *z*, and *η*. These values can be used as fixed parameters in the freqpcr() function. The “knownqpcr_unpaired” function was developed to handle incomplete data (i.e., the observations of *τ*^HW^, *τ*^TW^, *τ*^HD^, and *τ*^TD^ have different data lengths). If the four Cq measures are available for all samples, then “knownqpcr” is used.

Another objective of the analysis with known-ratio samples is to test the homoscedasticity of the qPCR data at the scale of Cq measures. Regarding the relationship between the etoxazole-R allele frequency in *T. urticae* and the corresponding 2^−ΔΔCq^ measures (the approximate point estimate of the frequency), Osakabe *et al*. (2017) demonstrated linearity using a sample series of DNA with multiple mixing ratios on CHS1 (I1017F). In the next section, we recycled the same data to compare whether the Cq measurements in the RED-ΔΔCq analysis obey the homoscedasticity in the scale of ΔΔCq or (1 + *η*)^−ΔΔCq^.

## Materials and Methods

### Experiment 1: estimation of auxiliary parameters and verification of homoscedasticity in Cq measurements based on mite DNA samples with known allele-mixing ratios

#### Experimental setup

In the experiment by Osakabe *et al*. (2017), the resistant mite strain (SoOm1-etoR strain) originated from a field population collected in Omaezaki City, Shizuoka, Japan (34.7°N, 138.1°E) in January 2012. The susceptible strain was obtained from Kyoyu Agri Co., Ltd. (Kanagawa, Japan) (Kyoyu-S strain). For each strain, two pairs of females and males were used separately. Each pair was allowed to mate and oviposit on a kidney bean leaf square (2 × 2 cm) for four days. The mites were then confirmed to be homozygous on the CHS1 locus using sequence analysis. Genomic DNA extracted from the offspring of each pair was used for qPCR analysis. For each pair, the DNA extracts were prepared twice, each of which was a mixture from 50 adult females homogenized together, that is, four extracts (replicates) for each strain.

To verify the validity of the RED-ΔΔCq method, qPCR analysis was performed with heterogeneous DNA solutions with ten mixing ratios of *X*_R_/(*X*_R_ + *X*_S_) = {0, 0.001, 0.005, 0.01, 0.05, 0.1, 0.25, 0.5, 0.75, 1}. The net DNA concentration of each mixed solution was adjusted to 1 ng μl^−1^, from which 15 ng was dispensed into each of the two tubes. Only one was digested with the restriction enzymes before qPCR. For digestion, the samples were treated with a mixture of two enzymes, *MluC* I (10 units) and *Taq*^α^I (20 units; New England BioLabs, Ipswich, MA, USA), at 37 °C for 3 h, followed by incubation at 65 °C for 3 h. This is due to the polymorphism of the CHS1 loci; the 1017 codon of *T. urticae* displays ATT (Kyoyu-S strain) or TTT (SoOm1-etoR) sequences, whereas the upstream 1016 codon displays a synonymous TCG or TCA independent of the strains (Van Leeuwen et al., 2012). Therefore, we need to digest both TC<u>GATT</u> (underline shows the restriction site of *Taq*^α^I) and TC<u>AATT</u> (*MluC* I) to diminish the entire S allele.

qPCR analysis using the intercalator method was performed using the LightCycler Nano System (Roche Diagnostics, Basel, Switzerland) with SYBR Fast qPCR Mix (Takara, Kusatsu, Japan) as described previously (Osakabe et al., 2017). The primer sets were *tu03CHS1* (forward: 5′-GGCACTGCTTCATCCACAAG-3′ and reverse: 5′-GTGTTCCCCAAGTAACAACGTTC-3′) and *tu25GAPDH* (forward: 5′-GCACCAAGTGCTAAAGCATGGAG-3′ and reverse: 5′-GAACTGGAACACGGAAAGCCATAC-3′).

#### Statistical analysis

The maximum likelihood of *δ*_B_, *δ*_T_, *σ*_c_, *z*, and *η* was conducted with the “knownqpcr_unpaired” function of the freqpcr package (version 0.3.2). The raw Cq data are available as ESM 1 along with a step-by-step guide for statistical analyses (ESM 2). Due to the limitation of the handling capacity of the thermal cycler, qPCR analysis was not conducted on undigested samples of the nine mixing ratios other than *X*_R_/(*X*_R_ + *X*_S_) = 1 (i.e., pure R solution). Thus, in each replicate, Osakabe *et al*. (2017) used the observed Δ*τ*^W^ value when the ratio = 1 for other ratios to calculate the conventional ΔΔCq indices. As we have shown in Eq. 7, this operation does not affect the point estimates of *p*, although the size of the Cq measurement error (σ_c_) will be underestimated if we recycle the observed Cq value multiple times.

Regarding the relationship between the true mixing ratio and the RED-ΔΔCq measures in the sample, the linearity was analyzed using a linear model via the function “lm” running on R version 3.6.1 (R Core Team, 2019), where the response variables were put into the model at the scale of Cq or (1 + η)^−ΔΔCq^. Based on the linear models, we tested heteroscedasticity using the Breusch-Pagan test via the bptest() function of the R library “lmtest” (Hothorn et al., 2019).

### Experiment 2: evaluation of the simultaneous estimation method with randomly generated data

Since the experiment by Osakabe *et al*. (2017) used a sample series with strict mixing ratios, the effect of individual differences in DNA yield was not evaluated. Instead, we conducted a numerical experiment to verify the accuracy of the simultaneous parameter estimation under uncertainty in the individual DNA yield. The frequency of the R allele in the population, *p*, was set to 0.01, 0.05, 0.1, 0.25, 0.5, or 0.75.

For the sampling strategy, *N* bulk samples (the parameter ‘ntrap’ in the R source code), each comprising *n* individuals (*n* was fixed among the samples: the parameter ‘npertrap’ in the code), were generated by random sampling from a wild population of a haploid organism. To assess how the estimation interval responds to the sample sizes, we evaluated the combination of *N* = {2, 4, 8, 16, 32, 64} and *n* = {4, 8, 16, 32, 64}, though the combinations with *Nn >* 128 were excluded (*Nn* corresponds to ‘ntotal’ in the code). The DNA quantities (*δ*_R_ and *X*_S_) contained in each bulk sample were generated as random numbers that followed the gamma distributions of Eq. 3. To cover a plausible variability range of the DNA yield, the gamma shape parameter was varied as *k* = {1, 3, 9, 27}. Depending on the size of *k* the gamma scale parameter was set at *θ* = 1 × 10^−6^/*k* to fix the mean of the individual DNA yield to 1 × 10^−6^. The termination threshold for qPCR, *X*_Θ_ was fixed at 1.

We fixed the other parameters due to limitations of the computing resources. From the results of Experiment 1, *δ*_T_ = 1.2, *δ*_B_ = 0.24, *z* = 0.0016, and *η* = 0.97 were presupposed. As for the random errors in the PCR amplification process and/or the Cq measurement, *σ*_c_ = 0.2 was assumed regardless of the initial template quantity. For each of the 624 parameter regions, the dummy datasets comprising *N* bulk samples were generated 1,000 times independently with different random number seeds (i.e., 1,000 replicates), for which the parameter estimation with freqpcr(…, beta = TRUE) of the freqpcr package version 0.3.1 was run on the R 3.6.1 environment. The simulation code is available in ESM 3.

As we also implemented the gamma distribution model as freqpcr(…, beta = FALSE), a numerical experiment with the gamma model was also conducted for the first 250 replicates, and the estimation accuracy was compared between the two assumptions. Furthermore, we also fitted the function with the settings freqpcr(…, K = 1), that is, assuming the gamma shape parameter was fixed at 1 (a.k.a. exponential distribution), in addition to the default simulation with all parameters (*p, k, δ*_T_, and *σ*_c_) unknown. Further, the easiest way to estimate *p* derived from Eq. 8, we averaged the observed ΔΔCq values for *N* bulk samples and transformed them as 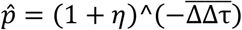.

### Results

#### Estimation of auxiliary parameters and verification of homoscedasticity

Based on the Cq measures, the auxiliary parameters were estimated based on the RED-ΔΔCq analysis of the I1017F mutation of *T. urticae*. As for the initial quantity of template DNA (the parameter “meanDNA” on the R code; defined as *X*/*X*_Θ_), the maximum-likelihood estimate was 1.256 × 10^-6^ (95% confidence interval: 7.722 × 10^-7^ to 2.041 × 10^-6^). The relative quantity of the target gene to the housekeeping gene *δ_T_* (targetScale) was estimated to be 1.170 (95% CI: 1.069-1.280). The locus-independent change rate in the template quantity accompanying the restriction enzyme treatment *δ*_B_ (baseChange) was 0.2361 (95% CI: 0.2040 to 0.2731). The measurement error in the scale of Cq *σ_c_* (SD) was 0.2376 (95% CI: 0.2050 to 0.2755). The residue rate of the S allele after digestion *z* (zeroAmount) was 0.001564 (95% CI: 0.001197-0.002044). The efficiency of amplification per PCR cycle *η* (EPCR) was 0.9712 (95% CI: 0.9231-1.022).

In the RED-ΔΔCq analysis of the etoxazole resistance of *T. urticae*, the relationship between the true R allele frequency (*Y*_R_ = *X*_R_/(*X*_R_ + *X*_S_) in the sample) and the corresponding Cq measures exhibited higher homoscedasticity in the scale of the measured ΔΔCq values rather than in (1 + η)^−ΔΔCq^, the transformation to *Ŷ*_R_ (Figure 2). The linear regression of the ΔΔCq values on —ln[0.001564 × (1 — *Y*_R_) + *Y*_R_]/ln(1 + 0.971) showed high linearity (intercept = −0.07694, coefficient = 1.025, adjusted R^2^ = 0.9936). The homoscedasticity of the coefficient of determination was not rejected at the 5% level of significance (Breusch-Pagan test: BP = 3.1577, *df* = 1, *p* = 0.07557) (Figure 2A). Conversely, the linear regression of 1.971^−ΔΔCq^ on [0.001564 × (1 — *Y*_R_) + *Y*_R_] showed a slightly lower linearity (intercept = −0.008625, coefficient = 1.092, adjusted R^2^ = 0.9709). The Breusch-Pagan test was highly significant (BP = 13.978, *df* = 1, *p* = 0.0001849), rejecting homoscedasticity (Figure 2B). These results suggest that it is easier to model the error structure of the RED-ΔΔCq method on the scale of Cq values (logarithm) rather than frequency (linear scale).

**Figure 2.**
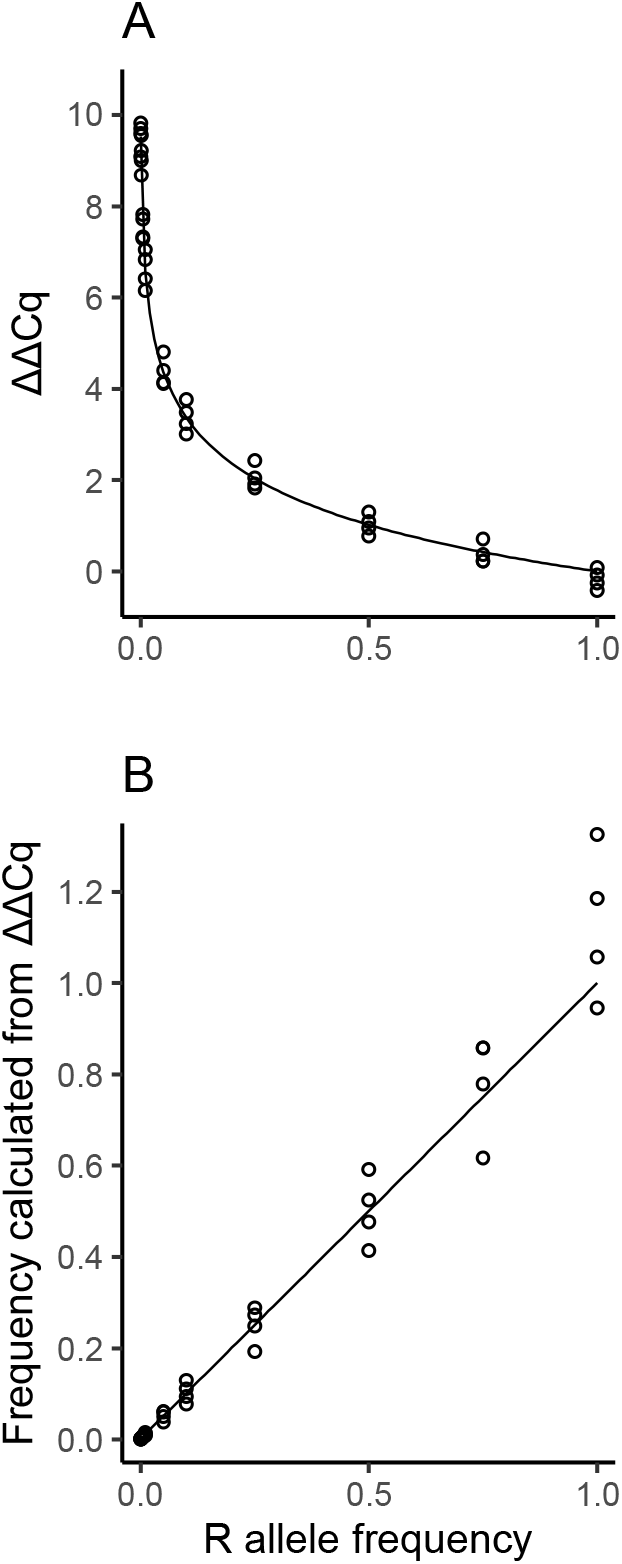
Relationship between the allele frequency in the sample and A: the RED-ΔΔCq measures, B: the observed frequency calculated as (1 + *η*)^(–ΔΔCq), showing the results of etoxazole resistance in the two-spotted spider mites. The lines are not the regression on the actual Cq measurement (shown as points), but the theoretical relationship between true frequency of the R allele and the quantity defined as A: – ln(*z* + *Y_R_*(1 – z))/ln(1 + *η*) or B: *z* + *Y_R_*(1 – *z*), where *Y*_R_ = *X*_R_/(*X*_R_ + *X*_S_). Parameters are *z* = 0.00156 and *η* = 0.971.

#### Evaluation of the simultaneous estimation method with randomly generated data

Among the 624 parameter regions of the numerical simulation with 1,000 replicates (250 for the gamma model), the total success rate of the interval estimation*p* using freqpcr(…, beta = TRUE) was 70.6% and 94.5% when all parameters were unknown, and when the gamma shape parameter was fixed as *k* = 1. The “success rate” here indicates the probability when the function returns certain values other than NA (i.e., the diagonal of the Hessian was not negative): no guarantee that the estimated confidence interval was accurate. The estimation success for the Cq measurement error, *σ*_c_, was 69.6% and 97.6% in the beta-distribution model with unknown *k* and *k* = 1, respectively. The relative quantity of the target gene, *δ*_T_, was 68.1% and 96.1%, respectively. However, the estimated success of *k* was 59.9% with the beta distribution model, showing a lower performance than the other parameters, implying that the likelihood is insensitive to the size of *k*. Conversely, the estimation of *p* is robust to the size of *k*, as we show later in this section.

The estimation success of freqpcr() largely depended on the total sample size (*Nn* corresponding to the facet ‘ntotal’ in the figures), as well as the level of *p* (Figure S1 and S2 for the beta and gamma models, with all parameters unknown). In each parameter region, the quantity Bin(0|*Nn, p*) generally gives the probability that the whole sample contains no R individuals. When *Nn* is larger enough, *Nn* > 3/*p* is approximately the requirement for the total sample size to contain at least one R individual with 95% confidence, called the “rule of three” (Eypasch et al., 1995). The gray backgrounds in the facets of Figures 3–4 and S1–S7 signify the regions where the total sample sizes are smaller than the thresholds (e.g., 60 haploid individuals are required when *p* = 0.05). As shown in Figures S1 and S2, the parameter estimation often failed when *Nn* did not meet the rule of three. Once we exclude the parameter regions of *Nn* ≤ 3/*p*, the estimation success rate of *p* with freqpcr(…, beta = TRUE) improved to 84.3% and 99.9% with all parameters unknown and assuming *k* = 1, respectively.

**Figure 3.**
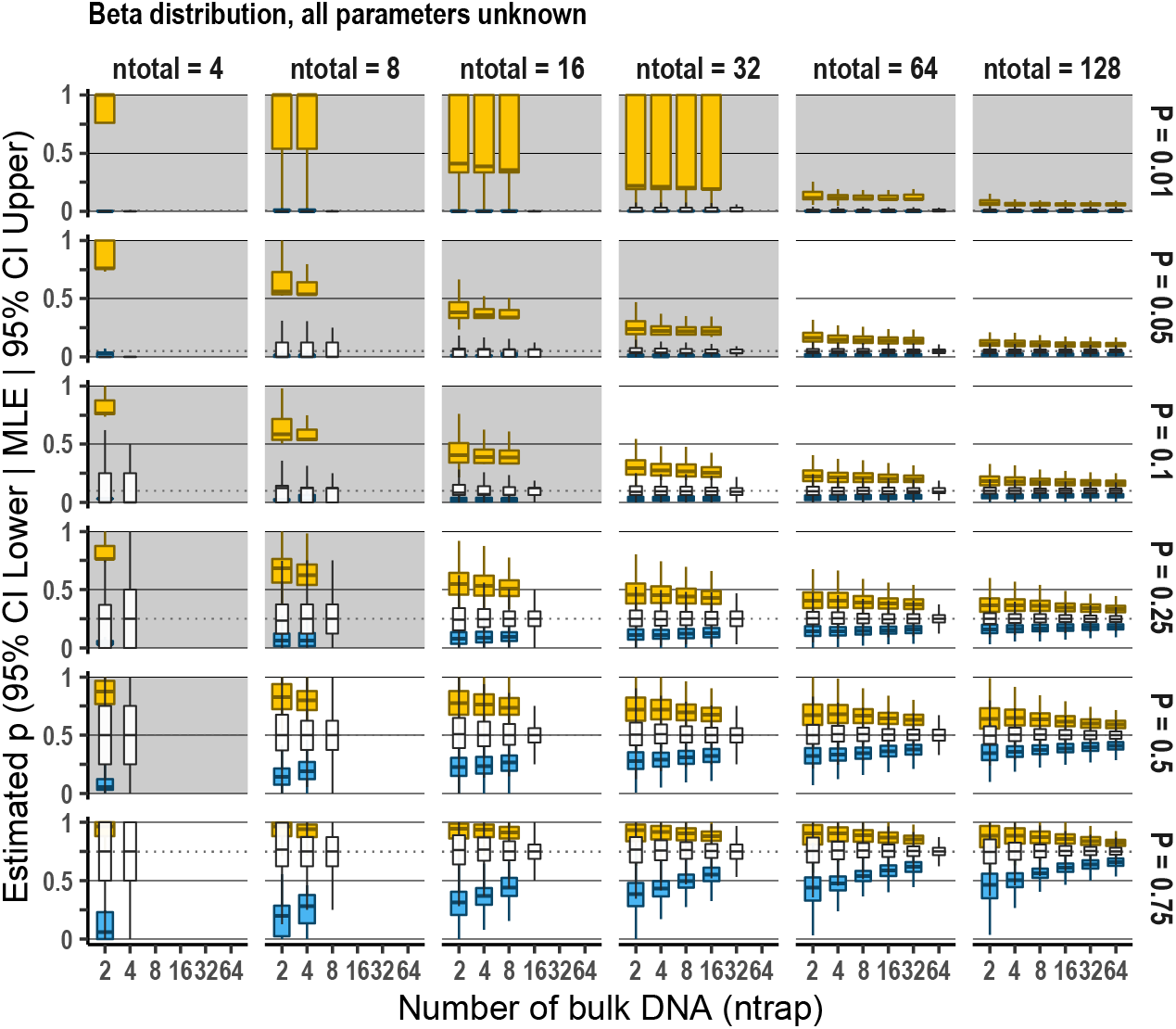
Estimation accuracy of the population allele frequency, *p*, with freqpcr() when the beta distribution was assumed, and all estimable parameters (P, K, targetScale, and sdMeasure) were set as unknown. The result of numerical experiments based on 1,000 dummy datasets per parameter region. The x-axes corresponds to *N*, or the “ntrap” parameter, the extent to which the collected individuals (ntotal) were divided to the bulk samples. The three box plots (white thin, blue, and yellow wide) in each region show the maximum likelihood estimates (MLE), lower bound of the 95% CI, and the upper bound, respectively. In each boxplot, the horizontal line signifies the median of the simulations, hinges of the box show 25 and 75 percentiles, and the upper/lower whiskers correspond to the 1.5 × interquartile ranges. The shaded facets show that the total sample sizes (ntotal) are smaller than 3/*p*.

**Figure 4.**
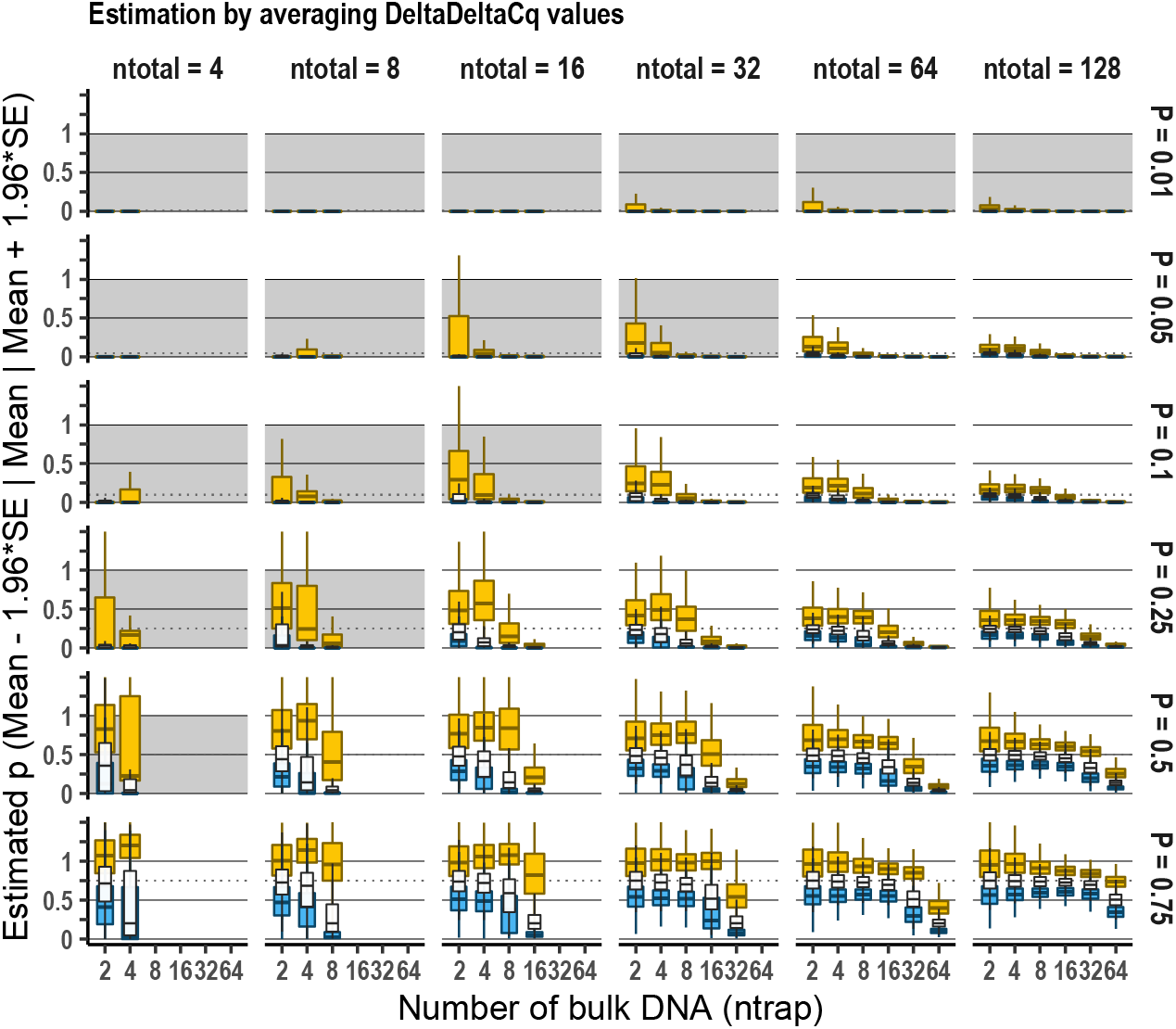
Estimation accuracy of the population allele frequency by simple averaging of ΔΔCq measures. The frequency was underestimated than its true value (horizontal broken line in each facet) as the samples were more divided. The Cq dataset was derived from the numerical experiment of “beta distribution, all parameters unknown.”

As for the estimation accuracy of *p*, the freqpcr() function assuming beta distribution provides an unbiased estimator. Figures 3 and S3 show the estimated sizes of *p* using the beta model with all parameters unknown and assuming *k* = 1, respectively. Both settings demonstrated that the estimator converged to the true R frequency; the upper/lower bounds of the estimated 95% confidence intervals (yellow/blue boxes in each plot) became narrower as we increased the total sample sizes (*Nn*) or included more bulk DNA samples (*N*). Fixing the size of the gamma shape parameter to *k* = 1scarcely affected the point estimates and intervals of p, as long as *Nn* > 3/p is satisfied (Figure S3). However, if every individual was analyzed separately, the interval estimation was only possible when *k* was fixed (see the regions of “sample division = ntotal” cases in Figure 3).

When we used the gamma distribution model, the interval estimation of *p* was also possible and unbiased (Figure S4). However, when we defined the point estimator of *p* as a simple average, that is, 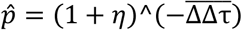, it was strongly underestimated as the samples were more divided (*N*/*Nn* was large) (Figure 4). The upper limit of 95% CI often violated 1, suggesting that the “simple average of ΔΔCq” ± 1.96 SE is inadequate for the interval estimation based on the RED-ΔΔCq method.

Although the freqpcr() function with the gamma and beta distributions both showed an unbiased estimation of *p*, the gamma model was disadvantageous regarding calculation time and the number of iterations before convergence. The time varied largely in the model settings and sample sizes (Figures S5–S7). Among the settings we tried, beta model with fixed *k* was the fastest and converged within a few seconds in most parameter regions (median and 75 percentile: 0.32 and 0.69 s: Figure S6). It was three and >10 times faster than the beta (0.91 and 2.4 seconds: Figure S5) and gamma (3.0 and 15 s: Figure S7) model, respectively with all parameters unknown. The calculation time increased as the dataset size increased - *Nn* and the sample was more divided (larger *N*/*Nn*) in the beta distribution model, because the marginal likelihood was calculated for each bulk sample (Figures S5 and S6). Conversely, the gamma distribution model (Figure S7) requires increased calculation time as the size of each bulk sample becomes larger (larger *n_h_*). This was considered because the combination of Bin(*m_h_*|*n_h_, p*) exploded when *n_h_* was large.

Furthermore, the estimation accuracy of the shape parameter, k, it was underestimated as the real size of the parameter increased (e.g., *k* = 27) when the gamma distribution model was applied (Figure S8B). Since the iterative fitting of the parameter in freqpcr() always starts internally from *k* = 1 (this was determined due to the calculation stability), this bias suggests the likelihood function of *ψ*_G_ (Eq. 13), with little information on the size of *k* compared with*p*. Then, *k* tends to stay at its initial value, suggesting that the gamma model is not suitable for the simultaneous estimation of *p* and *k*. Unlike the gamma version, the fitting of *k* with freqpcr(beta = TRUE) was satisfactory when we divided the total samples into more bulk samples (larger *N*/*Nn*), although the initial value dependence was still observed, especially when *p* or *N* was small (Figure S8A). This may be because the estimation of *k* via Beta(*m_h_k*, (*n_h_* – *m_h_*)/*k*) in Eq. 14 is comparable with measuring the overdispersion of *Y*_*h*,R|*m_h_*_, which is only possible when multiple bulk samples contain both R and S alleles.

## Discussion

In the present study, we developed a statistical model to estimate the population allele frequency based on qPCR across multiple bulk samples to address the issues facing the conventional point estimator for allele frequency which averages the observed ΔΔCq values 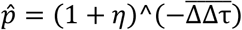. This conventional method sometimes exceeds 1 when the frequency of the target allele is close to 1. Furthermore, when one tries to quantify the rare mutant allele in a population, most bulk samples contain only the wild type allele. The conventional 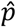 is vulnerable to many zero samples, which makes the frequency estimation more difficult when*p* is small. To circumvent these problems, our interval estimation explicitly models the number of individuals contained in each bulk sample (the binomial assumption) as well as the individual DNA yields (the gamma assumption), thereby obtaining the interval estimate over the entire range 0 < *p* < 1.

The explicit modeling of individuals also allows sample division to various degrees, which helps us to balance our sampling strategy on the cost-precision tradeoff. We can achieve higher precision (narrower confidence interval) by increasing the total sample size, 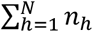 although it also increases the costs associated with sample collection and laboratory work, including library preparation and PCR analysis. Recent advances in molecular diagnosis have relieved sampling costs. However, although it is possible to now extract DNA from dead insect bodies obtained from sticky traps (Uesugi et al., 2016), a larger sample size still imposes a larger handling cost if we analyze the collected organisms individually via non-quantitative PCR.

The combination of mass trapping and bulk qPCR analysis solves the latter challenge by collecting more individuals and pooling them. This can result in higher precision with less work than individual PCR. For instance, we sampled 16 individuals from the population with an allele frequency of *p* = 0.05 and analyzed two individuals once in the numerical experiment (Figure 3: facet of ntotal = 16, sample division = 8). The lower and upper limits of the 95% confidence interval *p* were estimated to be 0.0087 and 0.34, respectively, using freqpcr(…, beta = TRUE) (as the medians of the 1,000 independent trials). We also simulated the case of ntotal = 64 and sample division = 4 (i.e., analyzed 16 individuals together) and found the upper and lower limits to be 0.015 and 0.15, respectively. Thus, we improved the precision of the interval estimate with half the handling effort.

Also, in non-quantitative PCR, sample pooling is considered as a tool for the detection of rare (c)DNA in the population with practical labor requirements, and has been used as high throughput pre-screening system for many samples e.g. in clinical examinations (Taylor et al., 2010; Yelin et al., 2020). However, in some fields, such as plant quarantine, it is important to guarantee that a product is not contaminated with pests or unapproved genetically modified seeds at a certain consumer risk. As the assumed frequency range is low (p ≈ 0.001), frequency estimation is not realistic (3,000 seeds are needed to meet the “rule of three” when *p* = 0.001) and is not required for the current inspection routine. Thus, group testing based on non-quantitative PCR has been conducted in these fields (Yamamura et al., 2019). Yamamura and Hino (2007) proposed a procedure to estimate the upper limit of the population allele frequency, in which they used the proportion of bulk samples detected as “positive.”

Overall, there has been a gap in methodology between the frequency estimation based on the individual PCR and the non- or semi-quantitative PCR based on the non-quantitative bulk PCR. Although it provides the highest estimation precision following binomial distribution, the former is only available at a higher *p*; it becomes labor-intensive once we try to quantify rare alleles. The latter can be applied to a lower range of *p*, but the precision is generally low or even non-quantitative. Bridging the gap, our qPCR-based procedure offers an allele frequency estimation in the mid-low range (*p* = 0.01 to 0.25), which is considered a critical range for decision making in some fields like pesticide resistance management (Takahashi et al., 2017; Sudo et al., 2018).

Although this study focused on resistance genes, the likelihood model in Eq. 11 can also be applied for other qPCR protocols based on ΔΔCq. If both the specific and nonspecific primer sets are available to amplify the “mutant” and “wild type + mutant” alleles at the target locus, they can be used for the test and control samples equivalent to *X^TV^* in Eq. 10 and *X^TU^* in Eq. 9, respectively. However, there is a caveat in determining which allele should be amplified with a specific primer set and which affects the estimation accuracy due to the intrinsic nature of (1 + η)^−ΔΔτ^. As shown, the 95% confidence intervals were broader when *p* = 0.75 than when *p* = 0.25 (Figure 3), the accuracy was not symmetric around 0.5, but more accurate when the frequency was low. That is, one should design a specific primer set to amplify the allele that would be rare in the population to improve the signal-to-noise ratio.

The maximum likelihood estimation with freqpcr() relies on the assumption that the quantities of the S and R alleles in each bulk sample independently follow gamma distribution and that their quotient is expressed using beta distribution. Fixing the size of the gamma shape parameter *k* further accelerated the optimization, which was owing to the robustness of *p* to the size of k. However, once the size of *k* was fixed much larger than the actual size of the gamma shape parameter (i.e., the individual DNA yield was regarded as almost a fixed value), the iterative optimization using the nlm() function sometimes returned an error. Therefore, one should start with a smaller shape parameter e.g., *k* = 1 (the exponential distribution: Figure S3), which is currently the default setting of the freqpcr package.

In qPCR applications for diagnostic use, ΔΔCq is often used with calibration. One of the popular methods is the involvement of technical replicates; each sample is dispensed and analyzed using qPCR multiple times, which negates the Cq measurement error. The measurement error obeys a homoscedastic normal distribution in the Cq scale, as shown in Experiment 1. Thus, a simple solution is to average the Cq values measured for each bulk sample before the estimation with freqpcr(), although the estimated size of *σ*_c_ changes from its original definition in Eq. 7. However, it is trivial if the number of technical replicates is unified between bulk samples.

Moreover, the comparison of Cq values is sometimes conducted on more than one internal reference as there is no guarantee that the expression level of a “housekeeping gene” is always constant (Vandesompele et al., 2002). Future updates of freqpcr() will handle multiple internal references. As long as qPCR is used to estimate population allele frequency, the use of statistical inferences on the bulk samples, as presented in this study, will continue to be a realistic option for regional allele monitoring and screening for practitioners, such as those in agricultural, food security, and public health sectors.

## Supporting information

Appendix S1 Formularization in the case of diploidy.

ESM1: mite Cq data, ESM2: R source for Figure 1, ESM3: simulation code.

## Acknowledgments

We appreciate Dr. Kohji Yamamura and Dr. Takehiko Yamanaka for earlier discussion on the gamma assumption of the individual DNA yield. The work was supported by a grant from the Ministry of Agriculture, Forestry, and Fisheries of Japan (Genomics-based Technology for Agricultural Improvement): PRM05 to M.O. and PRM07 to M.S.

## Data accessibility

The R package source is available at https://github.com/sudoms/freqpcr. The output data of the numerical experiment are available at https://figshare.com/collections/freqpcr/5258027. The source code for the figures, including the mite dataset from Osakabe *et al*. (2017), are available as electronic supplementary materials.

### ESM 1

RED-ΔΔCq dataset from Osakabe et al. (2017).

### ESM 2

R source code for Experiment 1 (Figure 2), including a brief guide to the “freqpcr” package.

### ESM 3

R source code for the numerical simulation (Experiment 2) and the codes for Figures 3 and after.

### Appendix S1

Formularization in the case of diploidy.

## Author contributions

M.S. designed research, made statistical models and R package, and analyzed data. M.O. conducted laboratory work. Both authors wrote the final manuscript.

## Figures

**Figure S1.**
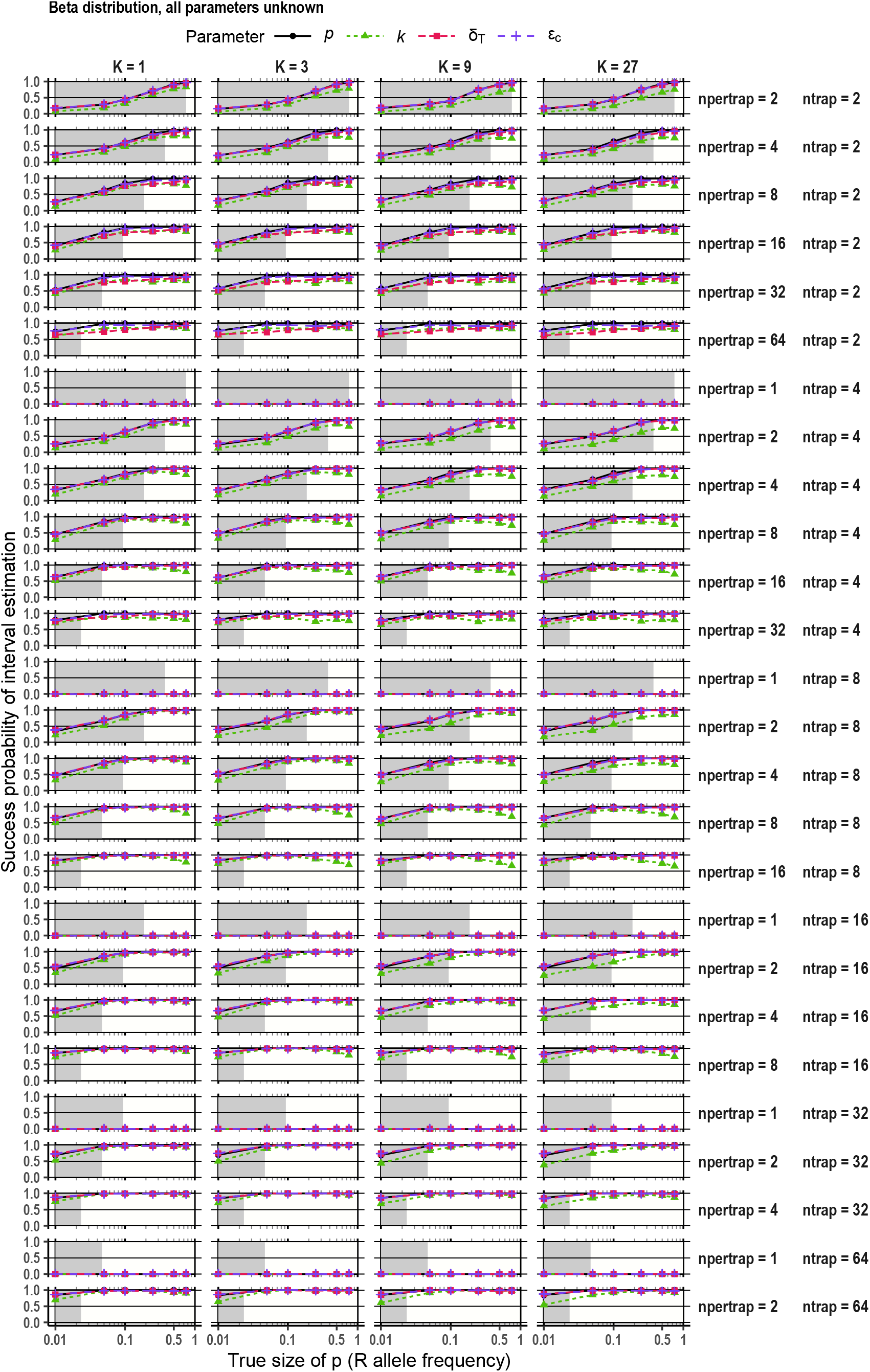
Probability of estimation success with freqpcr(). The beta distribution was assumed, and all estimable parameters (P, K, targetScale, and sdMeasure) were set as unknown. The shaded boxes in the background show the frequency ranges where the total sample sizes (ntotal) are smaller than 3/*p*.

**Figure S2.**
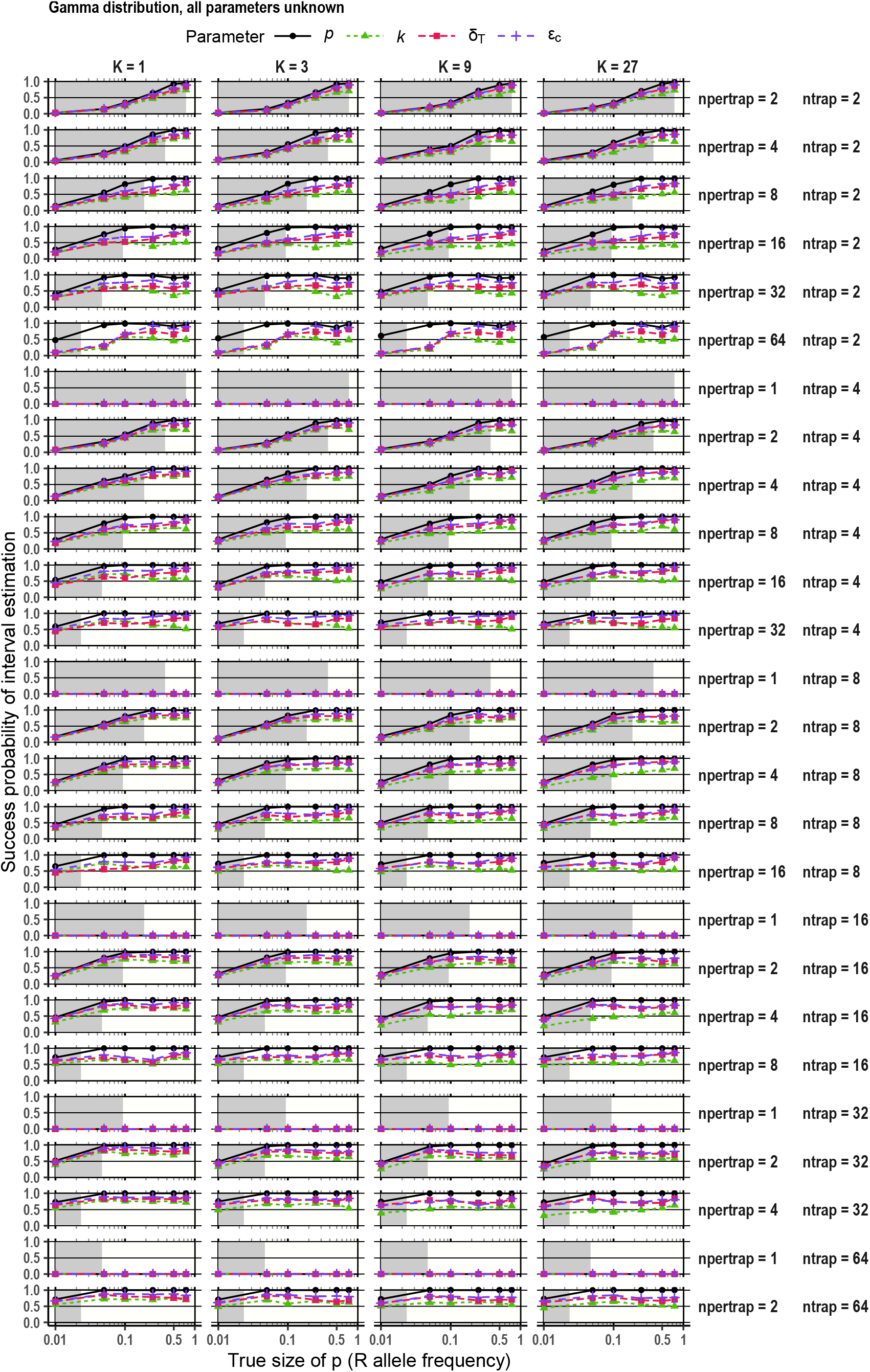
Probability of estimation success with freqpcr(). The gamma distributions were assumed, and all estimable parameters were set as unknown. The function often failed to calculate the CIs for *k* when npertrap (individuals in each bulk sample) were larger.

**Figure S3.**
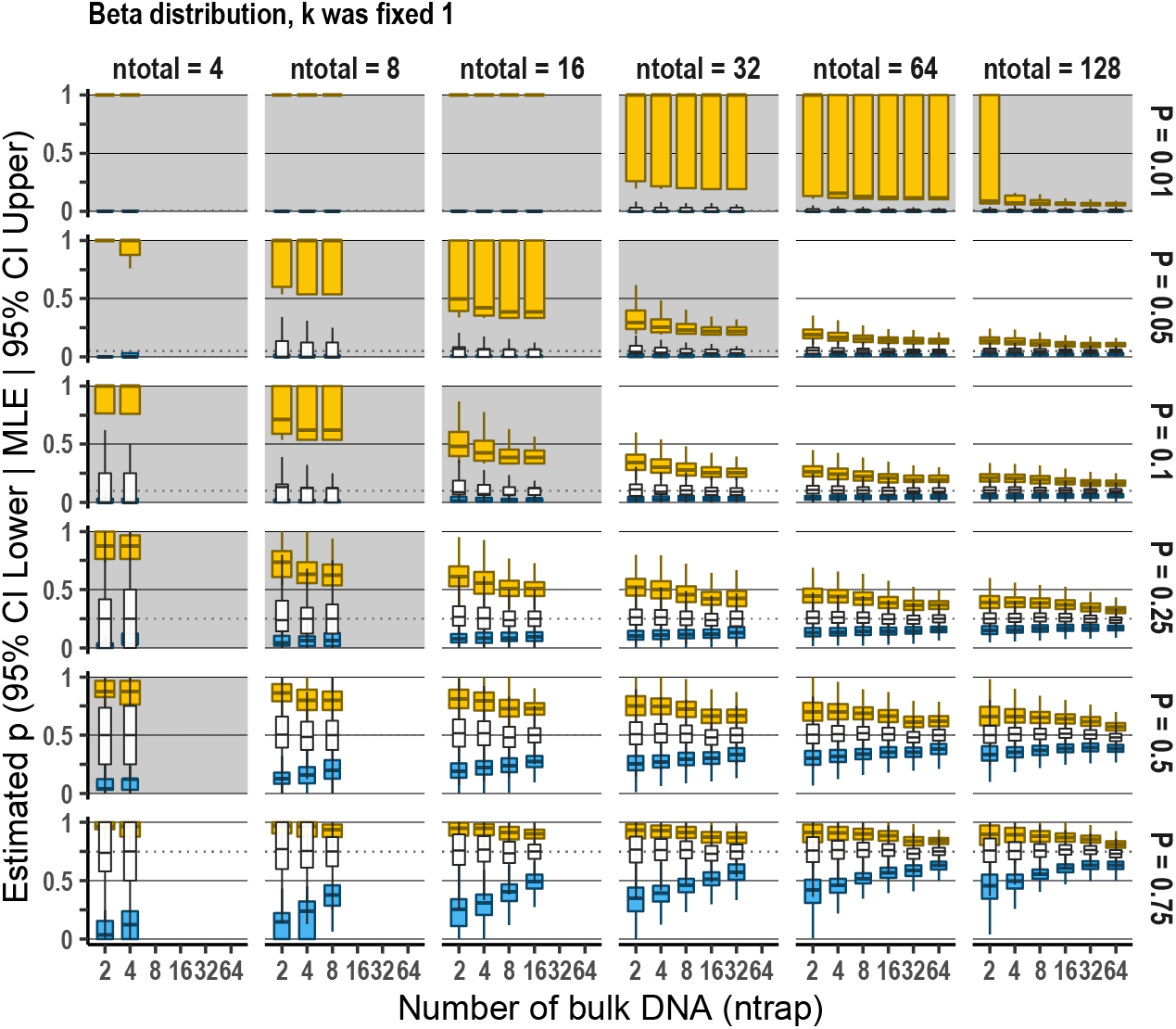
Estimation accuracy of the population allele frequency, *p*, with freqpcr() when the beta distribution was assumed, considering K = 1.

**Figure S4.**
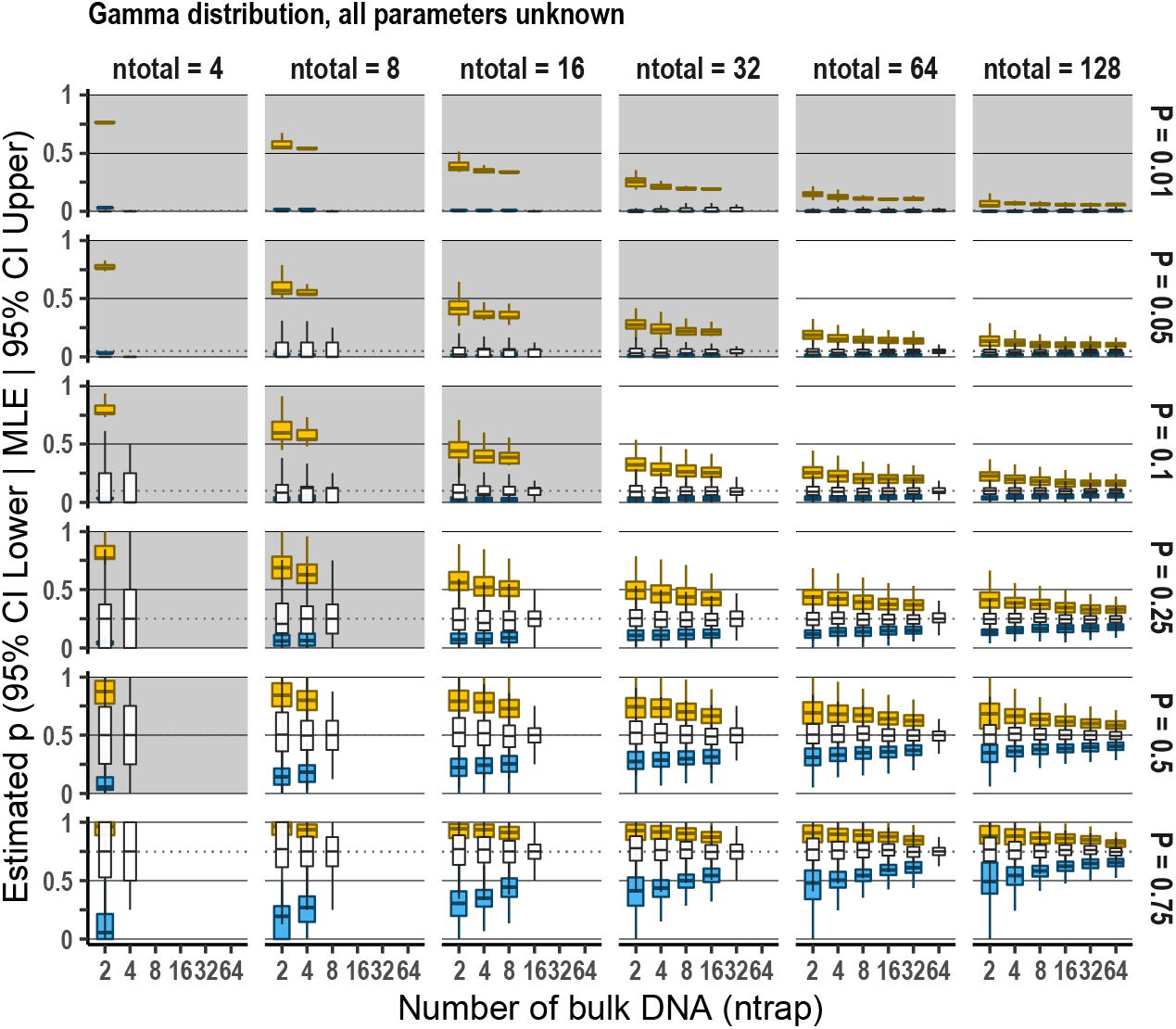
Estimation accuracy of *p* with freqpcr() when gamma distributions were assumed and all estimable parameters were set as unknown.

**Figure S5.**
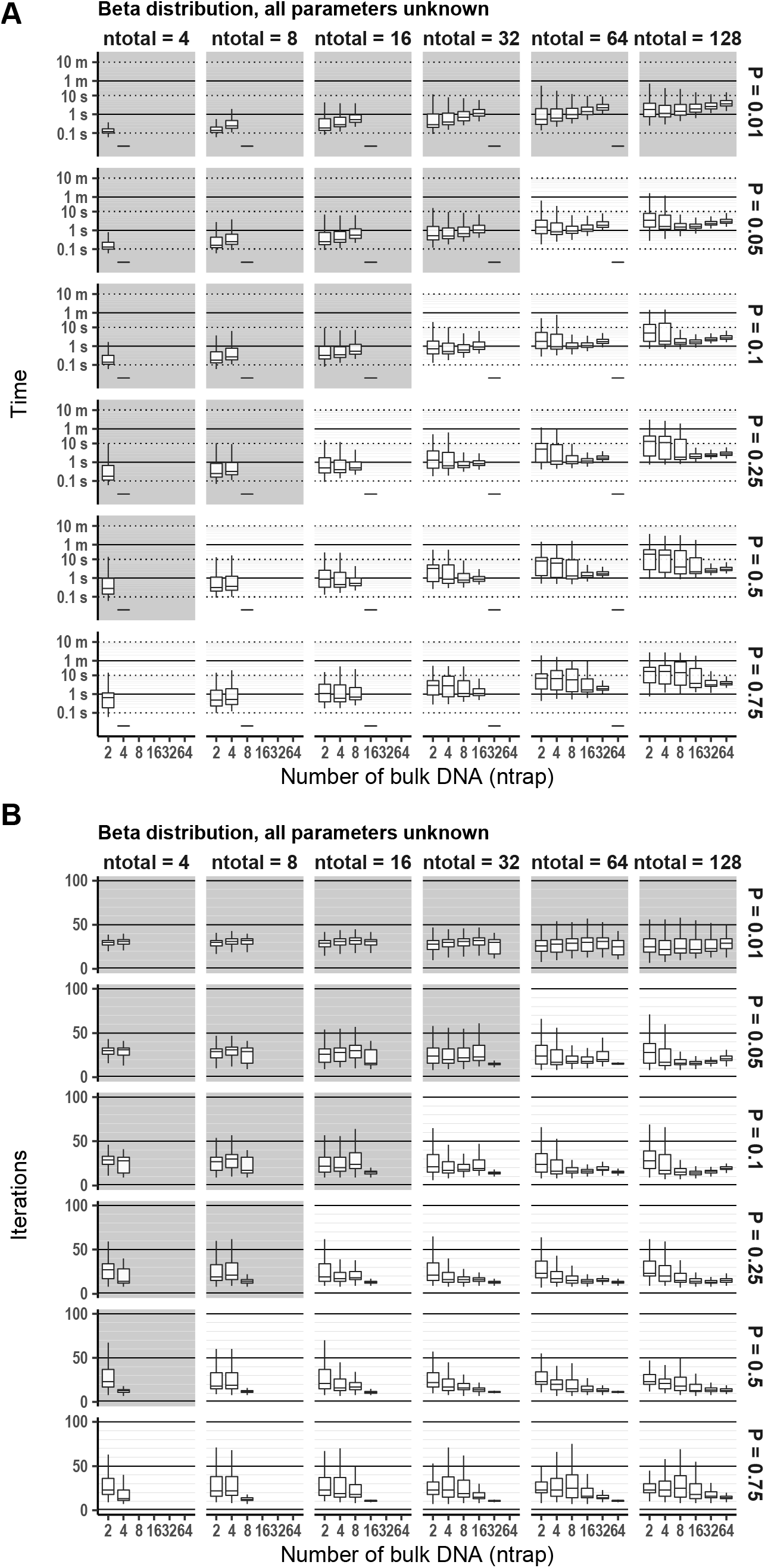
Calculation time (A) and number of iterations (B) until the freqpcr() function converges. The beta distribution was assumed, and all estimable parameters were set as unknown.

**Figure S6.**
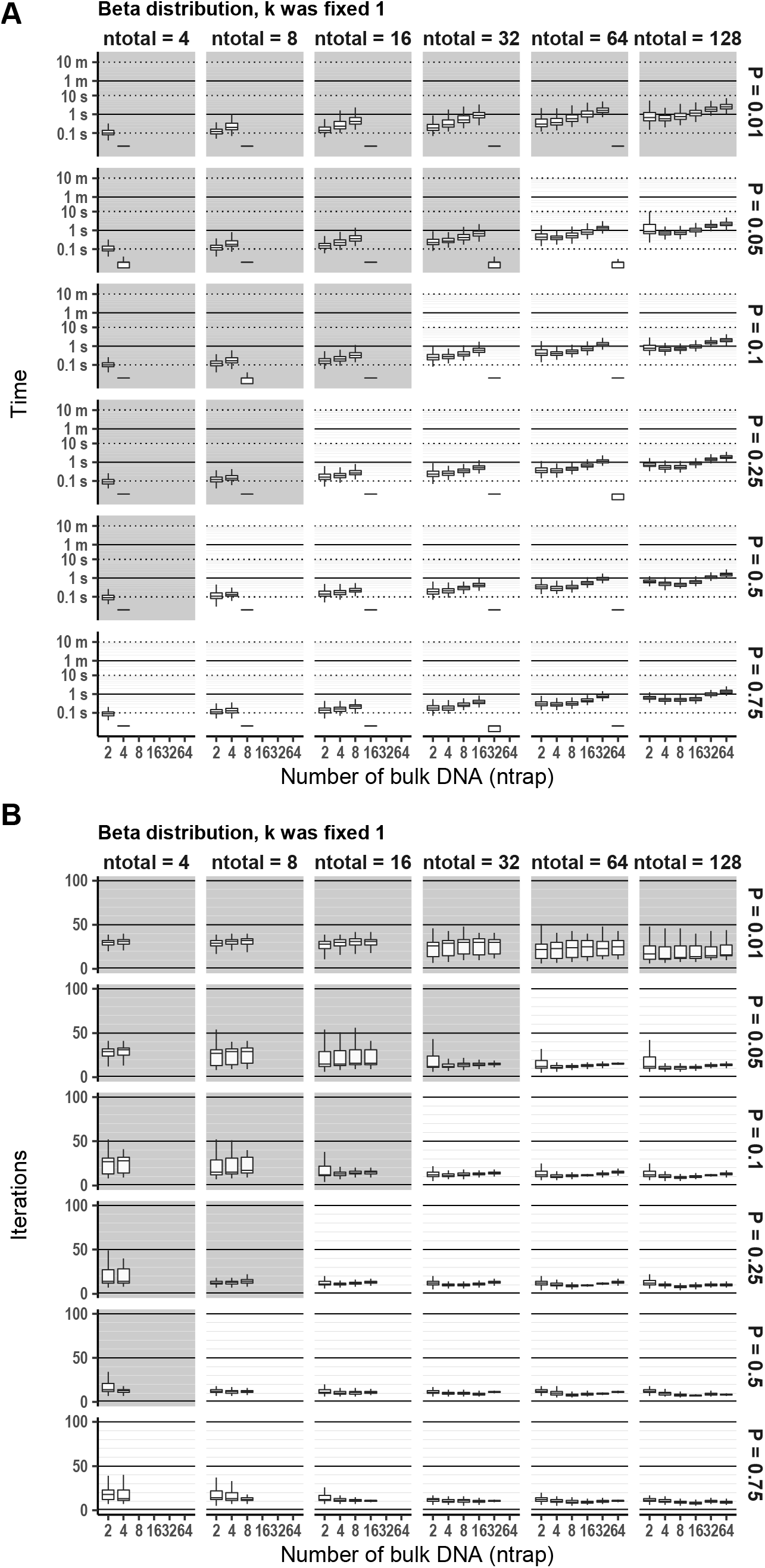
Calculation time (A) and number of iterations (B) until the freqpcr() function converges. The beta distribution was assumed, fixing the gamma shape parameter K = 1.

**Figure S7.**
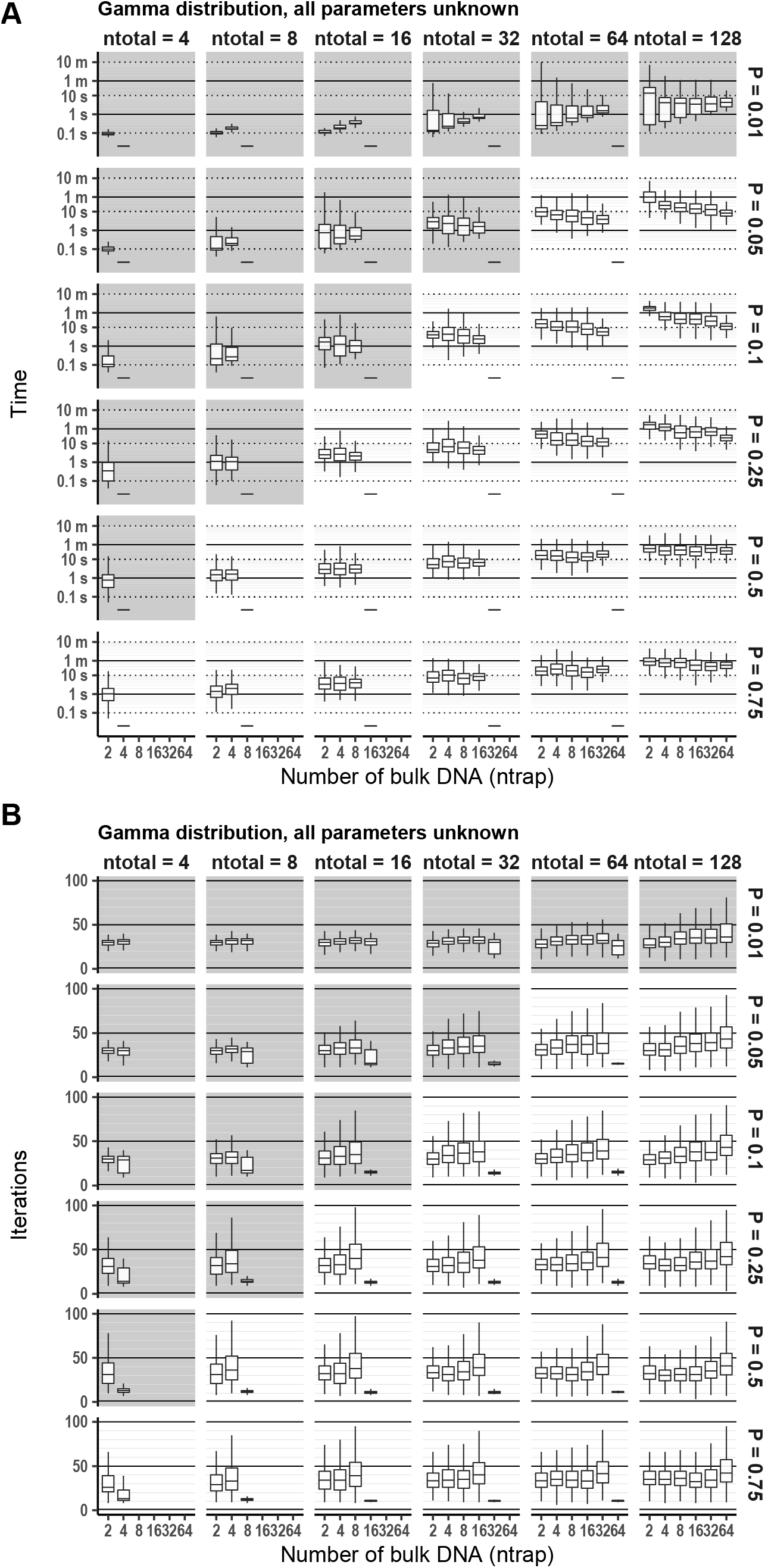
Calculation time (A) and number of iterations (B) until the freqpcr() function converges, assuming gamma distributions. All estimable parameters were set as unknown.

**Figure S8.**
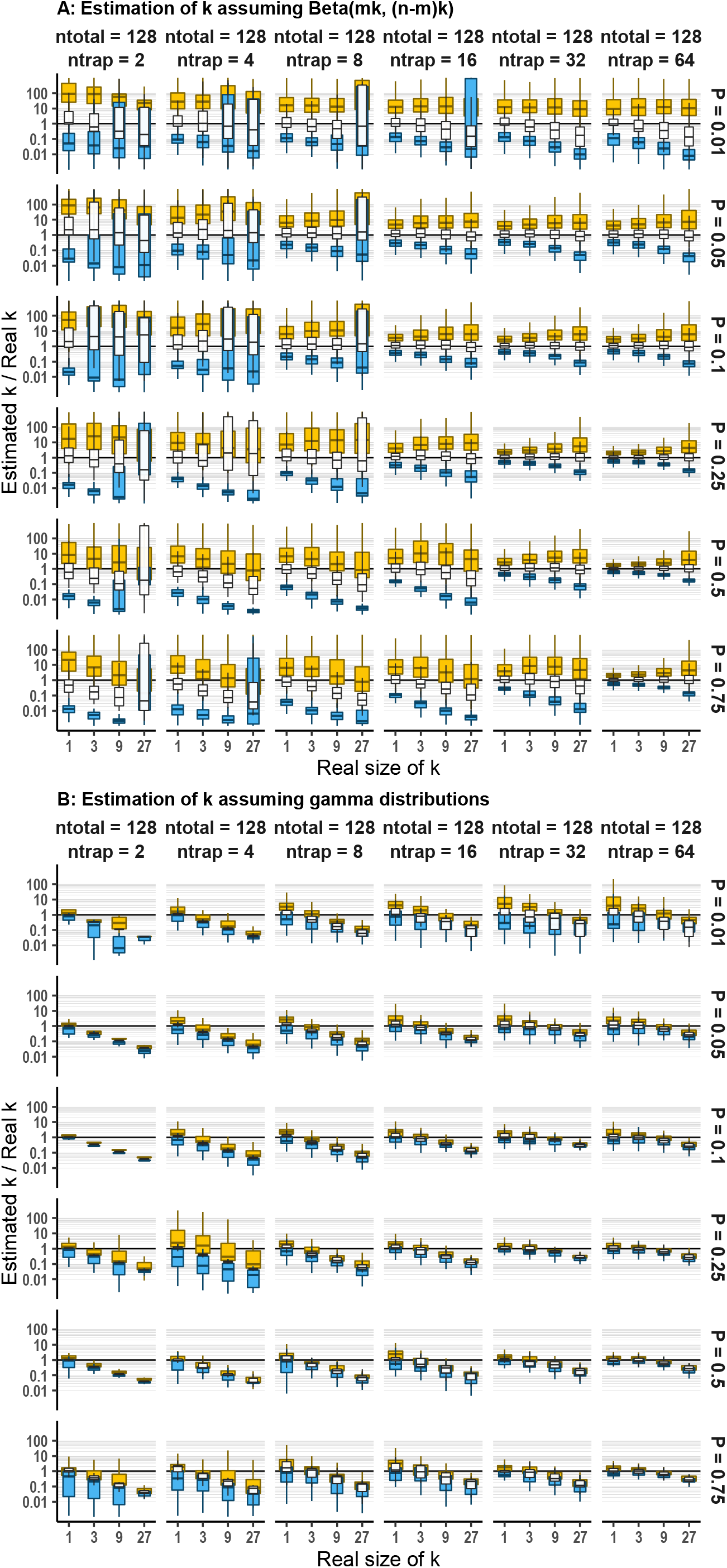
Estimation accuracy of *k* (the gamma shape parameter) in the simulation, showing the maximum likelihood estimate by freqpcr() divided by the actual parameter size.

