## Appendix S1 Formularization in the case of diploidy. for "freqpcr: estimation of population allele frequency using qPCR ΔΔCq measures from bulk samples"

### *Appendix S1: Case of Diploidy*

Although we considered sampling from haploid organisms, many insects and vertebrates are diploid. Let us consider that the population of a diploid insect species has the R allele frequency $p$, from which we collected $n$ individuals. The bulk sample then consists of $m_1$ ($m_1 = 0,1,,\ldots,n$) individuals of RR homozygotes, $n - m_1 - m_0$ RS heterozygotes, and $m_0$ ($m_0 = 0,1,,\ldots,n$) SS homozygotes ($m_1 + m_0 \leq n$). The joint probability of obtaining $\{m_1, m_0\}$ obeys the trinomial distribution with probabilities $p^2$ and $(1-p)^2$

$$\mathrm{Tri}(m_1, m_0 | n, p^2, (1-p)^2) = \frac{n!}{m_1!\, m_0!\, (n - m_1 - m_0)!} \cdot p^{2m_1} \cdot (1-p)^{2m_0} \cdot (2p - 2p^2)^{(n - m_1 - m_0)}.$$

*Eq. 15*

The total R allele in the bulk sample comes from two R/R sets contained in the $m_1$ homozygotes and a single set of R from the $n - m_1 - m_0$ heterozygotes. Likewise, two S/S sets from $m_0$ homozygotes and a single S set from the $n - m_1 - m_0$ heterozygotes constitute the total S body. Note that the yields of R and S from these heterozygotes would be the same unless there is a genotype-dependent systematic error in the extraction efficiency.

Let us define the amount of DNA copies per genome: the random variable $X_{*\in(\mathrm{S,R})|\mathrm{homo}}$ for the yield of S or R from the homozygotes, and $X_{*\in(\mathrm{S,R})|\mathrm{hetero}}$ for S or R from the heterozygotes. As in the case of haploidy, $X_\mathrm{R}$ and $X_\mathrm{S}$ denote the allele contents in the bulk sample; they are the linear combinations of $X_{*|\mathrm{homo}}$ and $X_{*|\mathrm{hetero}}$:

$$X_\mathrm{R} = 2 \times X_{\mathrm{R|homo}} + X_{\mathrm{R|hetero}}, \quad X_\mathrm{S} = X_{\mathrm{S|hetero}} + 2 \times X_{\mathrm{S|homo}},$$

$$2 \times X_{\mathrm{R|homo}} \sim \mathrm{Ga}(m_1 k, 2\theta), \quad X_{\mathrm{R|hetero}} \sim \mathrm{Ga}\big((n - m_1 - m_0)k, \theta\big),$$
$$X_{\mathrm{S|hetero}} = X_{\mathrm{R|hetero}}, \qquad 2 \times X_{\mathrm{S|homo}} \sim \mathrm{Ga}(m_0 k, 2\theta).$$

*Eq. 16*

#### *Parameter estimation*

There are $n - i + 1$ cases from $m_0 = 0$ to $m_0 = n - i$ when the number of RR homozygotes is given by $m_1 = i$. The segregation ratio in the bulk sample has $\sum_{i=0}^{n}(n - i + 1)$ total combinations. For each combination of $n$, $m_0$, and $m_1$, Eq. 16 gives the probability of obtaining the ΔCq measures in Eq. 11. However, a drawback arises from the constraint of the amounts of R and S possessed by heterozygotes. The applicability of the likelihood model (Eq. 13 or Eq. 14 in the main text) depends largely on the independence of $X_\mathrm{R}$ and $X_\mathrm{S}$. If we define the likelihood using Eq. 16 as it was, we must convolve the DNA amounts not on the two-dimensional parameter space spanned by $X_{\mathrm{R}}$ and $X_{\mathrm{S}}$, but a three-dimensional space by $X_{\mathrm{R|homo}}$, $X_{\mathrm{S|hetero}} = X_{\mathrm{R|hetero}}$, and $X_{\mathrm{S|homo}}$, which would increase the calculation time.

Therefore, we removed the constraint and assumed that $X_{\mathrm{R|*}}$ and $X_{\mathrm{S|*}}$ were distributed independently and identically; that is, instead of the heterozygotes, we captured $n - m_1 - m_0$ individuals of haploid R and another $n - m_1 - m_0$ individuals of haploid S separately. Regarding homozygotes, we also assumed that we captured $2m_1$ R haploids and $2m_0$ S haploids instead of $m_1$ RR and $m_0$ SS, respectively. Then,

$$X_{\mathrm{R|homo}} \sim \mathrm{Ga}(2m_1 k, \theta), \qquad X_{\mathrm{R|hetero}} \sim \mathrm{Ga}\big((n - m_1 - m_0)k, \theta\big),$$
$$X_{\mathrm{S|hetero}} \sim \mathrm{Ga}\big((n - m_1 - m_0)k, \theta\big)\ i.i.d., \qquad X_{\mathrm{S|homo}} \sim \mathrm{Ga}(2m_0 k, \theta).$$

*Eq. 17*

Finally, we can approximate the DNA amounts of a diploid organism in the bulk sample by simply substituting Eq. 3 in the main text:

$$X_{\mathrm{R}} \sim \mathrm{Ga}\big((n + m_1 - m_0)k, \theta\big), \quad X_{\mathrm{S}} \sim \mathrm{Ga}\big((n - m_1 + m_0)k, \theta\big).$$

*Eq. 18*

In addition, at probability $\mathrm{Bin}(0|2n_h, p)$, all (hypothetically haploid) individuals become S or R; in that case, there is no need to convolve the DNA amounts.
