## Supplementary material for "freqpcr: estimation of population allele frequency using qPCR ΔΔCq measures from bulk samples": ESM1: mite Cq data, ESM2: R source for Figure 1, ESM3: simulation code.: ESM2_code_for_figures.R

```
################################################################################################################################

# Name: freqpcr
# Short description: interval estimation of population allele frequency based on \u0394\u0394Cq measures of quantitative PCR over bulk samples
# Author: Masaaki Sudo (NARO, JAPAN)
# Maintainer: Masaaki Sudo
## https://github.com/sudoms/freqpcr
## https://orcid.org/0000-0001-9834-9857

################################################################################################################################

Tested environment:

> sessionInfo()
R version 3.6.1 (2019-07-05)
Platform: x86_64-w64-mingw32/x64 (64-bit)
Running under: Windows 10 x64 (build 18363)

Matrix products: default

Random number generation:
 RNG:     Mersenne-Twister
 Normal:  Inversion
 Sample:  Rounding

locale:
[1] LC_COLLATE=Japanese_Japan.932  LC_CTYPE=Japanese_Japan.932    LC_MONETARY=Japanese_Japan.932 LC_NUMERIC=C
[5] LC_TIME=Japanese_Japan.932

attached base packages:
[1] parallel  stats     graphics  grDevices utils     datasets  methods   base

other attached packages:
 [1] installr_0.22.0 furrr_0.1.0     future_1.16.0   forcats_0.4.0   stringr_1.4.0   dplyr_0.8.4     purrr_0.3.3
 [8] readr_1.3.1     tidyr_1.0.2     tibble_2.1.3    ggplot2_3.2.1   tidyverse_1.3.0

loaded via a namespace (and not attached):
 [1] tidyselect_1.0.0 listenv_0.8.0    haven_2.2.0      lattice_0.20-40  colorspace_1.4-1 vctrs_0.2.3      generics_0.0.2
 [8] utf8_1.1.4       rlang_0.4.5      pillar_1.4.3     glue_1.3.1       withr_2.1.2      DBI_1.1.0        dbplyr_1.4.2
[15] modelr_0.1.6     readxl_1.3.1     lifecycle_0.1.0  munsell_0.5.0    gtable_0.3.0     cellranger_1.1.0 rvest_0.3.5
[22] codetools_0.2-16 fansi_0.4.1      broom_0.5.4      Rcpp_1.0.3       scales_1.1.0     backports_1.1.5  jsonlite_1.6.1
[29] fs_1.3.1         hms_0.5.3        digest_0.6.25    stringi_1.4.6    grid_3.6.1       cli_2.0.1        tools_3.6.1
[36] magrittr_1.5     lazyeval_0.2.2   crayon_1.3.4     pkgconfig_2.0.3  xml2_1.2.2       reprex_0.3.0     lubridate_1.7.4
[43] assertthat_0.2.1 httr_1.4.1       rstudioapi_0.11  R6_2.4.1         globals_0.12.5   nlme_3.1-144     compiler_3.6.1


################################################################################################################################


# for the first time
library(devtools)
Sys.setenv(R_REMOTES_NO_ERRORS_FROM_WARNINGS="true") # If there are error (converted from warning)
install_github("sudoms/freqpcr")


# or install by hand
install.packages("freqpcr_0.3.2.tar.gz", repos=NULL, type="source")


# load the package
library(freqpcr); packageVersion("freqpcr");


################################################################################################################################
# Part I
# Procedure to determine parameters
################################################################################################################################


# Experiment 1
# Example dataset: DNA solution with known mixing ratio

# DNA was extracted from the etoxazole resistant (R) or susceptible (S) populations of
# the two-spotted spider mite, Tetranychus urticae.
# Copyright by Masahiro Osakabe (Kyoto University) 2017

# Osakabe, M., T. Imamura, R. Nakano, S. Kamikawa, M. Tadatsu, Y. Kunimoto, and M. Doi. 2017.
# Combination of restriction endonuclease digestion with the DeltaDeltaCt method in real-time PCR to monitor
# etoxazole resistance allele frequency in the two-spotted spider mite.
# Pesticide biochemistry and physiology 139:1–8.


# reading raw experiment data
library(readxl)
library(tidyverse)
mite.raw <- readxl::read_xlsx(  path="ESM1_mite.xlsx", sheet="Sheet1",
                                col_types=c(rep("text", 1), rep("numeric", 8))  )
data.frame(mite.raw)


> data.frame(mite.raw)
    Gene numR numS Digest FreqTrue     Cq.1     Cq.2     Cq.3     Cq.4
1   CHS1   50    0      1    1.000 21.76011 22.46276 21.67745 21.89234
2   CHS1   50   50      1    0.750 22.22055 22.73058 22.49938 22.13801
3   CHS1   50   50      1    0.500 22.55539 22.92551 23.13384 22.67460
4   CHS1   50   50      1    0.250 23.72164 24.22364 24.21292 23.96121
5   CHS1   50   50      1    0.100 25.02524 25.64115 25.37562 24.97376
6   CHS1   50   50      1    0.050 26.22723 26.48074 26.38646 25.98100
7   CHS1   50   50      1    0.010 28.86382 28.44911 28.68426 28.17910
8   CHS1   50   50      1    0.005 29.23326 29.45677 29.40738 28.98874
9   CHS1   50   50      1    0.001 30.36377 31.16006 30.66308 30.84527
10  CHS1    0   50      1    0.000 30.91856 31.39115 31.86532 31.48347
11  CHS1   50    0      0    1.000 19.76440 19.93363 19.81920 19.58301
12 GAPDH   50    0      1    1.000 22.25536 22.78204 22.21400 22.25800
13 GAPDH   50   50      1    0.750 22.09678 22.42058 22.39418 22.19650
14 GAPDH   50   50      1    0.500 21.70864 22.02676 22.48203 22.00767
15 GAPDH   50   50      1    0.250 21.91582 22.19761 22.41978 22.41461
16 GAPDH   50   50      1    0.100 21.78486 22.27926 22.26315 22.25005
17 GAPDH   50   50      1    0.050 22.06956 22.07313 22.36781 22.14638
18 GAPDH   50   50      1    0.010 22.27531 21.80223 22.39260 22.30971
19 GAPDH   50   50      1    0.005 21.75721 22.03858 22.23731 21.94482
20 GAPDH   50   50      1    0.001 21.92781 22.00806 21.78034 21.91004
21 GAPDH    0   50      1    0.000 22.07731 22.09051 22.16355 22.17278
22 GAPDH   50    0      0    1.000 20.00866 20.33493 19.94018 19.86679

# Cq. 1 ~ 4 are the replicates

################################################################################################################################


# data format 1: "unpaired"

# Desired data unit used in the RED-DeltaDeltaCq analysis is a 'quartet', 
# the 2x2 combination of (target- or housekeeping-gene) x (undigested or digested).
# A quartet is needed to calculate a conventional DeltaDeltaCq value.

# If all replicates of each experimental setting are fulfilled with the complete observations of the 2x2 combination,
# the dataset is easily treated in the "paired" format (see below).

# An "unpaired" format is used when some of the experimental settings (mixing ratio, replicates, etc.) lacks
# the complete combination of the 2x2 condition.

# For instance, the dataset from Osakabe et al. (2017) had 10 mixing ratios,
# R/(S+R) = c(0, 1, 0.001, 0.005, 0.01, 0.05, 0.1, 0.25, 0.5, 0.75), and four replicates. 
# However, qPCR on the undigested samples were only conducted with mixing ratio=1.
# The dataset is thus incomplete (i.e., the undigesed data must be recycled to calculate 
# the conventional DeltaDeltaCq values), though it is often the case in real operations.

# A dataset of this kind is often prepared as a "longer" formatted tidy data (https://r4ds.had.co.nz/tidy-data.html#pivoting)
# though we can also prepare it as a "wider" tidy data appearing in the next "paired" format, allowing missing values.


mite.long <- mite.raw %>%
    tidyr::pivot_longer(cols=starts_with("Cq"), names_to=c("Type", "Repl"),
                        values_to=c("Cq"), names_sep="\\.", names_ptypes=list(Repl=factor()))
mite.long # "mite.long" is an intermediate data format.

mite.unpaired <- mite.long %>%
    dplyr::mutate(Gene=as.numeric(factor(Gene, levels=c("GAPDH", "CHS1"))) - 1) %>%
    dplyr::arrange(desc(FreqTrue), Repl, Gene, Digest) %>%
    dplyr::select(FreqTrue, Repl, Gene, Digest, Cq)
mite.unpaired
readr::write_csv(mite.unpaired, "mite.unpaired.csv")


> mite.long
# A tibble: 88 x 8
   Gene   numR  numS Digest FreqTrue Type  Repl     Cq
   <chr> <dbl> <dbl>  <dbl>    <dbl> <chr> <fct> <dbl>
 1 CHS1     50     0      1     1    Cq    1      21.8
 2 CHS1     50     0      1     1    Cq    2      22.5
 3 CHS1     50     0      1     1    Cq    3      21.7
 4 CHS1     50     0      1     1    Cq    4      21.9
 5 CHS1     50    50      1     0.75 Cq    1      22.2
 6 CHS1     50    50      1     0.75 Cq    2      22.7
 7 CHS1     50    50      1     0.75 Cq    3      22.5
 8 CHS1     50    50      1     0.75 Cq    4      22.1
 9 CHS1     50    50      1     0.5  Cq    1      22.6
10 CHS1     50    50      1     0.5  Cq    2      22.9
# ... with 78 more rows

> mite.unpaired
# A tibble: 88 x 5
   FreqTrue Repl   Gene Digest    Cq
      <dbl> <fct> <dbl>  <dbl> <dbl>
 1        1 1         0      0  20.0
 2        1 1         0      1  22.3
 3        1 1         1      0  19.8
 4        1 1         1      1  21.8
 5        1 2         0      0  20.3
 6        1 2         0      1  22.8
 7        1 2         1      0  19.9
 8        1 2         1      1  22.5
 9        1 3         0      0  19.9
10        1 3         0      1  22.2
# ... with 78 more rows


################################################################################################################################

# data format 2: "paired," allowing missing values

# The combinations of (target- or housekeeping-gene) x (undigested or digested) are provided as separated data columns.
# Unobserved Cq data are filled as NA (not available).


mite.paired <- mite.long %>%
    tidyr::pivot_wider( names_from=c("Type", "Digest", "Gene"), names_sep=".",
                        values_from="Cq" ) %>%
    dplyr::select(FreqTrue, Repl, target0=Cq.0.CHS1, target1=Cq.1.CHS1, housek0=Cq.0.GAPDH, housek1=Cq.1.GAPDH) %>%
    dplyr::mutate(del.w=target0-housek0, del.d=target1-housek1) %>%
    dplyr::mutate(deldel=del.d-del.w) %>%
    dplyr::mutate(FreqCalc=2^-deldel)
mite.paired
readr::write_csv(mite.paired, "mite.paired.csv")


> mite.paired
# A tibble: 40 x 10
   FreqTrue Repl  target0 target1 housek0 housek1  del.w   del.d  deldel FreqCalc
      <dbl> <fct>   <dbl>   <dbl>   <dbl>   <dbl>  <dbl>   <dbl>   <dbl>    <dbl>
 1     1    1        19.8    21.8    20.0    22.3 -0.244 -0.495  -0.251     1.19
 2     1    2        19.9    22.5    20.3    22.8 -0.401 -0.319   0.0820    0.945
 3     1    3        19.8    21.7    19.9    22.2 -0.121 -0.537  -0.416     1.33
 4     1    4        19.6    21.9    19.9    22.3 -0.284 -0.366  -0.0819    1.06
 5     0.75 1        NA      22.2    NA      22.1 NA      0.124  NA        NA
 6     0.75 2        NA      22.7    NA      22.4 NA      0.310  NA        NA
 7     0.75 3        NA      22.5    NA      22.4 NA      0.105  NA        NA
 8     0.75 4        NA      22.1    NA      22.2 NA     -0.0585 NA        NA
 9     0.5  1        NA      22.6    NA      21.7 NA      0.847  NA        NA
10     0.5  2        NA      22.9    NA      22.0 NA      0.899  NA        NA
# ... with 30 more rows

####

# data format 2': "paired and recycled"

# If we adhere to calculate the conventional DeltaDelta Cq indices, 
# they need to be filled with ones of the correspinding experimental settings (data recycling).
# As for the mite dataset, unobserved Cq data are copied from the corresponding observations when FreqTrue == 1.


mite.paired.middle <- mite.long %>%
    dplyr::group_by(Gene, Repl) %>%
    tidyr::pivot_wider( names_from=c("Type", "Digest"), names_sep=".", values_from="Cq" ) %>%
    dplyr::mutate(Cq.0=mean(Cq.0, na.rm=TRUE))
mite.paired.middle

mite.paired.recycled <- mite.paired.middle %>%
    tidyr::pivot_wider( names_from=c("Gene"), names_sep=".", values_from=c("Cq.1", "Cq.0") ) %>%
    dplyr::select(FreqTrue, Repl, target0=Cq.0.CHS1, target1=Cq.1.CHS1, housek0=Cq.0.GAPDH, housek1=Cq.1.GAPDH) %>%
    dplyr::mutate(del.w=target0-housek0, del.d=target1-housek1) %>%
    dplyr::mutate(deldel=del.d-del.w) %>%
    dplyr::mutate(FreqCalc=2^-deldel)
mite.paired.recycled
readr::write_csv(mite.paired.recycled, "mite.paired.recycled.csv")


> mite.paired.middle
# A tibble: 80 x 7
# Groups:   Gene, Repl [8]
   Gene   numR  numS FreqTrue Repl   Cq.1  Cq.0
   <chr> <dbl> <dbl>    <dbl> <fct> <dbl> <dbl>
 1 CHS1     50     0     1    1      21.8  19.8
 2 CHS1     50     0     1    2      22.5  19.9
 3 CHS1     50     0     1    3      21.7  19.8
 4 CHS1     50     0     1    4      21.9  19.6
 5 CHS1     50    50     0.75 1      22.2  19.8
 6 CHS1     50    50     0.75 2      22.7  19.9
 7 CHS1     50    50     0.75 3      22.5  19.8
 8 CHS1     50    50     0.75 4      22.1  19.6
 9 CHS1     50    50     0.5  1      22.6  19.8
10 CHS1     50    50     0.5  2      22.9  19.9
# ... with 70 more rows

> mite.paired.recycled
# A tibble: 40 x 10
# Groups:   Repl [4]
   FreqTrue Repl  target0 target1 housek0 housek1  del.w   del.d  deldel FreqCalc
      <dbl> <fct>   <dbl>   <dbl>   <dbl>   <dbl>  <dbl>   <dbl>   <dbl>    <dbl>
 1     1    1        19.8    21.8    20.0    22.3 -0.244 -0.495  -0.251     1.19
 2     1    2        19.9    22.5    20.3    22.8 -0.401 -0.319   0.0820    0.945
 3     1    3        19.8    21.7    19.9    22.2 -0.121 -0.537  -0.416     1.33
 4     1    4        19.6    21.9    19.9    22.3 -0.284 -0.366  -0.0819    1.06
 5     0.75 1        19.8    22.2    20.0    22.1 -0.244  0.124   0.368     0.775
 6     0.75 2        19.9    22.7    20.3    22.4 -0.401  0.310   0.711     0.611
 7     0.75 3        19.8    22.5    19.9    22.4 -0.121  0.105   0.226     0.855
 8     0.75 4        19.6    22.1    19.9    22.2 -0.284 -0.0585  0.225     0.855
 9     0.5  1        19.8    22.6    20.0    21.7 -0.244  0.847   1.09      0.469
10     0.5  2        19.9    22.9    20.3    22.0 -0.401  0.899   1.30      0.406
# ... with 30 more rows


################################################################################################################################

# Maximum likelihood estimation on the Cq data taken from samples with known R ratios.

################################################################################################################################


1. Estimation of experimental parameters when the dataset was given in "unpaired" (i.e., long) format


library(freqpcr)
result <- knownqpcr_unpaired(   A=rep(1.0, nrow(mite.unpaired)), trueY=mite.unpaired$FreqTrue,
                                Gene=mite.unpaired$Gene, Digest=mite.unpaired$Digest, Cq=mite.unpaired$Cq,
                                method="BFGS", trace=1, report=10   )


initial  value 526.625717
iter  10 value 105.043851
iter  20 value 47.820456
iter  30 value -1.189728
final  value -1.601386
converged

Maximum-likelihood estimates with the two-sided 95% CIs
                                          Estimate         2.5%        97.5%
meanDNA (DNA content per unit)        1.255539e-06 7.721917e-07 2.041435e-06
targetScale (rel. target content)     1.170129e+00 1.069274e+00 1.280497e+00
baseChange (after digestion)          2.360702e-01 2.040327e-01 2.731381e-01
SD (Cq measurement error)             2.376223e-01 2.049847e-01 2.754564e-01
zeroAmount (S-target after digestion) 1.564246e-03 1.196945e-03 2.044261e-03
EPCR (PCR multiplification EPCRicacy) 9.712061e-01 9.231223e-01 1.021795e+00


####


2. Estimation of experimental parameters using the "paired" (i.e., wide) format dataset.

Although the undigested samples were quantified using qPCR only when FreqTrue == 1,
the "knownqpcr" function can deal with NA in the Cq data vectors (housek0, target0, housek1, and target1).

By contrast, no missing values are allowed for A (relative DNA contents between samples) and trueY (exact allele frequency).


result <- knownqpcr(A=rep(1.0, nrow(mite.paired)), trueY=mite.paired$FreqTrue,
                    housek0=mite.paired$housek0, target0=mite.paired$target0,
                    housek1=mite.paired$housek1, target1=mite.paired$target1,
                    method="BFGS", trace=1, report=10)


initial  value 526.625717
iter  10 value 105.043850
iter  20 value 47.820441
iter  30 value -1.189728
final  value -1.601386
converged

Maximum-likelihood estimates with the two-sided 95% CIs
                                          Estimate         2.5%        97.5%
meanDNA (DNA content per unit)        1.255539e-06 7.721917e-07 2.041435e-06
targetScale (rel. target content)     1.170129e+00 1.069274e+00 1.280497e+00
baseChange (after digestion)          2.360702e-01 2.040327e-01 2.731381e-01
SD (Cq measurement error)             2.376223e-01 2.049847e-01 2.754564e-01
zeroAmount (S-target after digestion) 1.564246e-03 1.196945e-03 2.044261e-03
EPCR (PCR multiplification EPCRicacy) 9.712061e-01 9.231223e-01 1.021795e+00


####


2'. Estimation of experimental parameters when the dataset was given in "paired" format, recycling the Cq values

"mite.paired.recycled" is the paired dataset, where the "undigested" Cq values other than FreqTrue == 1
were copied from the corresponding replicates of FreqTrue == 1.


result <- knownqpcr(A=rep(1.0, nrow(mite.paired.recycled)), trueY=mite.paired.recycled$FreqTrue,
                    housek0=mite.paired.recycled$housek0, target0=mite.paired.recycled$target0,
                    housek1=mite.paired.recycled$housek1, target1=mite.paired.recycled$target1,
                    method="BFGS", trace=1, report=10)


initial  value 811.268282
iter  10 value 189.797299
iter  20 value -5.694398
iter  30 value -26.773217
final  value -26.780544
converged

Maximum-likelihood estimates with the two-sided 95% CIs
                                          Estimate         2.5%        97.5%
meanDNA (DNA content per unit)        1.301427e-06 9.020759e-07 1.877571e-06
targetScale (rel. target content)     1.184654e+00 1.128954e+00 1.243103e+00
baseChange (after digestion)          2.379246e-01 2.203774e-01 2.568690e-01
SD (Cq measurement error)             2.046781e-01 1.834374e-01 2.283784e-01
zeroAmount (S-target after digestion) 1.583286e-03 1.267953e-03 1.977040e-03
EPCR (PCR multiplification EPCRicacy) 9.670477e-01 9.308465e-01 1.004657e+00


################################################################################################################################

# comparison of the estimates: "recycled" versus "allowing NA"
meanDNA     1.30e-6     1.26e-6
targetScale 1.18        1.17
baseChange  0.238       0.236
SD          0.205       0.238
zeroAmount  1.58e-3     1.56e-3
EPCR        0.967       0.971

You can see both results are almost same, but SD was underestimated when the Cq data are recycled.
As the original experimental design is "paired, allowing NA" in Osakabe et al. (2017), the parameter values calculated by:


knownqpcr(  A=rep(1.0, nrow(mite.paired)), trueY=mite.paired$FreqTrue,
            housek0=mite.paired$housek0, target0=mite.paired$target0,
            housek1=mite.paired$housek1, target1=mite.paired$target1,
            method="BFGS", trace=1, report=10  )

are used hereafter. That is,

meanDNA     1.26e-6
targetScale 1.17
baseChange  0.236
SD          0.238
zeroAmount  1.56e-3
EPCR        0.971


****
In the next section, however, DeltaDeltaCq and FreqCalc1971 are calculated on (2') "paird, recycled" dataset
because the calculation of each DeltaDeltaCq value needs complete observation of the four condition.
****

################################################################################################################################

# Show the relationship between the allele-mixing ratios and their Cq and DeltaDeltaCq values


# make the dataset including the calculated values of allele frequency based on EPCR = 0.971
mite <- mite.paired.recycled %>%
    dplyr::mutate(FreqCalc1971=1.971^-deldel)


# Test the heteroskedasticity around delta denta Cq

funcdeldel <- function (p, z=1.56e-03, EPCR=0.971) {
    -log(z*(1-p)+p)/log(1.0+EPCR) # predictor in the scale of Cq
}

funcfreq <- function (p, z=1.56e-03) {
    z*(1-p)+p # predictor in the scale of raw frequency
}


# Breusch-Pagan Test (function bptest) to test the heteroskedasticity
library(lmtest)
deldel.predictor <- funcdeldel(mite$FreqTrue, z=1.56e-03, EPCR=0.971)
freq.predictor <- funcfreq(mite$FreqTrue, z=1.56e-03)
deldel.lm <- lm(mite$deldel~deldel.predictor)
freq.lm <- lm(mite$FreqCalc1971~freq.predictor)
summary(deldel.lm)
summary(freq.lm)
lmtest::bptest(deldel.lm)
lmtest::bptest(freq.lm)


> summary(deldel.lm)

Call:
lm(formula = mite$deldel ~ deldel.predictor)

Residuals:
     Min       1Q   Median       3Q      Max
-0.60236 -0.19273 -0.00404  0.17686  0.61287

Coefficients:
                 Estimate Std. Error t value Pr(>|t|)
(Intercept)      -0.07694    0.07252  -1.061    0.295
deldel.predictor  1.02518    0.01319  77.703   <2e-16 ***
---
Signif. codes:  0 ‘***’ 0.001 ‘**’ 0.01 ‘*’ 0.05 ‘.’ 0.1 ‘ ’ 1

Residual standard error: 0.28 on 38 degrees of freedom
Multiple R-squared:  0.9937,	Adjusted R-squared:  0.9936
F-statistic:  6038 on 1 and 38 DF,  p-value: < 2.2e-16


> summary(freq.lm)

Call:
lm(formula = mite$FreqCalc1971 ~ freq.predictor)

Residuals:
      Min        1Q    Median        3Q       Max
-0.193350 -0.010329  0.007903  0.009899  0.242789

Coefficients:
                Estimate Std. Error t value Pr(>|t|)
(Intercept)    -0.008625   0.013154  -0.656    0.516
freq.predictor  1.091599   0.030257  36.078   <2e-16 ***
---
Signif. codes:  0 ‘***’ 0.001 ‘**’ 0.01 ‘*’ 0.05 ‘.’ 0.1 ‘ ’ 1

Residual standard error: 0.06555 on 38 degrees of freedom
Multiple R-squared:  0.9716,	Adjusted R-squared:  0.9709
F-statistic:  1302 on 1 and 38 DF,  p-value: < 2.2e-16


> lmtest::bptest(deldel.lm)

	studentized Breusch-Pagan test

data:  deldel.lm
BP = 3.1577, df = 1, p-value = 0.07557


> lmtest::bptest(freq.lm)

	studentized Breusch-Pagan test

data:  freq.lm
BP = 13.978, df = 1, p-value = 0.0001849

################################################################################################################################


########## code for Figure 2
# Error structure around the measured Cq values


deldel.unc <- function (p, scaleDNA=1.26e-06, zeroAmount=1.56e-03, EPCR=9.71e-01, trans=FALSE) {
    xR <- p*scaleDNA
    xS <- (1-p)*scaleDNA
    deldel <- -log((zeroAmount*xS+xR)/(xS+xR))/log(1+EPCR) # Eq. 8
    if (trans==FALSE) {
        ans <- deldel # As the scale of an expected Cq measure
    } else {
        ans <- (1+EPCR)^-deldel # As the transformation to the allele frequency
    }
    return(ans)
}


library(ggplot2)
library(patchwork)
LIM01 <- c(0, 1)
LIMY <- c(-1, 11)

p1 <- ggplot(mite) +
    stat_function(  fun=deldel.unc,
                    args=list(  scaleDNA=1.26e-06, zeroAmount=1.56e-03, EPCR=9.71e-01, 
                                trans=FALSE  ),
                    linetype=1, size=0.3  ) +
    geom_point(aes(y=deldel, x=FreqTrue), shape=1, size=1.0) +
    theme_classic(base_size=10) +
    coord_fixed(ratio=0.1) +
    scale_x_continuous(expand=c(0.02, 0.02), limit=LIM01, breaks=c(0, 0.5, 1)) +
    scale_y_continuous(expand=c(0, 0), limit=LIMY, breaks=seq(0, 10, by=2)) +
    labs( title="A", x="", y=expression(paste(italic(Delta), italic(Delta), "Cq", sep="")) )

p2 <- ggplot(mite) +
    stat_function(  fun=deldel.unc,
                    args=list(  scaleDNA=1.26e-06, zeroAmount=1.56e-03, EPCR=9.71e-01, 
                                trans=TRUE  ),
                    linetype=1, size=0.3  ) +
    geom_point(aes(y=FreqCalc1971, x=FreqTrue), shape=1, size=1.0) +
    theme_classic(base_size=10) +
    coord_fixed(ratio=1) +
    scale_x_continuous(expand=c(0.02, 0.02), limit=LIM01, breaks=c(0, 0.5, 1)) +
    scale_y_continuous(expand=c(0, 0), limit=c(-0.05, 1.35), breaks=seq(0, 1.3, by=0.2)) +
    labs( title="B",
          x=expression("R allele frequency"),
          y=expression(paste("Frequency calculated from ", italic(Delta), italic(Delta), "Cq", sep="")) )

p <- p1 / p2
ggsave(file="Figure2.pdf", device=cairo_pdf, plot=p, dpi=300, width=7/2.54, height=14/2.54)
ggsave(file="Figure2.eps", device=cairo_pdf, plot=p, dpi=300, width=7/2.54, height=14/2.54)
ggsave(file="Figure2.png", type="cairo", plot=p, dpi=300, width=7/2.54, height=14/2.54)

# save as the enhanced metafile 
library(devEMF)
emf(file="Figure2.emf", width=7/2.54, height=14/2.54, bg="transparent", pointsize=10, family="Arial", coordDPI=300)
p
dev.off()


################################################################################################################################
# Part II
# freqpcr: function of simultaneous parameter estimation
################################################################################################################################


# Make a dummy dataset for demonstration
# Test data are always generated in the "paired" format


library(freqpcr); packageVersion("freqpcr");
P <- 0.15
K <- 4
ntrap <- 4 # ntrap must be >= 1
npertrap <- 8 # npertrap must be >= 1
scaleDNA <- (1/K)*1e-06
targetScale <- 1.2
baseChange <- 0.2
EPCR <- 0.97
zeroAmount <- 1.6e-03
sdMeasure <- 0.2


# Make a dummy dataset (haploid)
dat.DNA <- make_dummy(  rand.seed=71, P, K, ntrap, npertrap, scaleDNA,
                        targetScale, baseChange, EPCR, zeroAmount, sdMeasure, diploid=FALSE  )
# Make a dummy dataset (diploid)
dat2.DNA <- make_dummy( rand.seed=71, P, K, ntrap, npertrap, scaleDNA,
                        targetScale, baseChange, EPCR, zeroAmount, sdMeasure, diploid=TRUE )
dat.DNA
dat2.DNA
(dat.gamma <- data.frame(   N=dat.DNA@N, housek0=dat.DNA@housek0, target0=dat.DNA@target0,
                            housek1=dat.DNA@housek1, target1=dat.DNA@target1,
                            RFreqMeasure=dat.DNA@RFreqMeasure, ObsP=dat.DNA@ObsP   ))
(dat2.gamma <- data.frame(  N=dat2.DNA@N, housek0=dat2.DNA@housek0, target0=dat2.DNA@target0,
                            housek1=dat2.DNA@housek1, target1=dat2.DNA@target1,
                            RFreqMeasure=dat2.DNA@RFreqMeasure, ObsP=dat2.DNA@ObsP  ))


> dat.DNA
An object of class "CqList"
Slot "N":
[1] 8 8 8 8

Slot "m":
     [,1] [,2] [,3] [,4]
[1,]    1    1    1    0
[2,]    7    7    7    8

Slot "xR":
[1] 6.654827e-07 1.540772e-06 7.998461e-07 0.000000e+00

Slot "xS":
[1] 5.541084e-06 7.909241e-06 8.052976e-06 1.004673e-05

Slot "housek0":
[1] 17.57120 16.90864 17.14117 16.88547

Slot "target0":
[1] 17.65665 16.84725 16.58306 16.90106

Slot "housek1":
[1] 19.90382 19.08542 19.59406 19.42896

Slot "target1":
[1] 23.13257 21.59854 22.97203 28.34278

Slot "DCW":
[1]  0.08544953 -0.06139074 -0.55811003  0.01559466

Slot "DCD":
[1] 3.228745 2.513126 3.377968 8.913814

Slot "deldel":
[1] 3.143296 2.574517 3.936078 8.898219

Slot "RFreqMeasure":
[1] 0.11868765 0.17453873 0.06933585 0.00239759

Slot "ObsP":
[1] 0.11868765 0.17453873 0.06933585 0.00239759

Slot "rand.seed":
[1] 71


> dat2.DNA
An object of class "CqList"
Slot "N":
[1] 8 8 8 8

Slot "m":
     [,1] [,2] [,3] [,4]
[1,]    0    0    0    0
[2,]    2    1    4    4
[3,]    6    7    4    4

Slot "xR":
[1] 1.599798e-06 1.249008e-06 4.770976e-06 3.342741e-06

Slot "xS":
[1] 1.513851e-05 1.706749e-05 1.564665e-05 9.480615e-06

Slot "housek0":
[1] 16.13263 16.32947 15.97995 16.30548

Slot "target0":
[1] 16.14822 15.66516 15.30663 16.40500

Slot "housek1":
[1] 18.67612 18.53178 18.06831 19.17237

Slot "target1":
[1] 21.53573 22.12281 20.64181 20.75895

Slot "DCW":
[1]  0.01559466 -0.66430459 -0.67331097  0.09952126

Slot "DCD":
[1] 2.859606 3.591031 2.573500 1.586582

Slot "deldel":
[1] 2.844011 4.255336 3.246811 1.487061

Slot "RFreqMeasure":
[1] 0.14539009 0.05584025 0.11064299 0.36484699

Slot "ObsP":
[1] 0.14539009 0.05584025 0.11064299 0.36484699

Slot "rand.seed":
[1] 71


> (dat.gamma <- data.frame(   N=dat.DNA@N, housek0=dat.DNA@housek0, target0=dat.DNA@target0,
+                             housek1=dat.DNA@housek1, target1=dat.DNA@target1,
+                             RFreqMeasure=dat.DNA@RFreqMeasure, ObsP=dat.DNA@ObsP   ))
  N  housek0  target0  housek1  target1 RFreqMeasure       ObsP
1 8 17.57120 17.65665 19.90382 23.13257   0.11868765 0.11868765
2 8 16.90864 16.84725 19.08542 21.59854   0.17453873 0.17453873
3 8 17.14117 16.58306 19.59406 22.97203   0.06933585 0.06933585
4 8 16.88547 16.90106 19.42896 28.34278   0.00239759 0.00239759

> (dat2.gamma <- data.frame(  N=dat2.DNA@N, housek0=dat2.DNA@housek0, target0=dat2.DNA@target0,
+                             housek1=dat2.DNA@housek1, target1=dat2.DNA@target1,
+                             RFreqMeasure=dat2.DNA@RFreqMeasure, ObsP=dat2.DNA@ObsP  ))
  N  housek0  target0  housek1  target1 RFreqMeasure       ObsP
1 8 16.13263 16.14822 18.67612 21.53573   0.14539009 0.14539009
2 8 16.32947 15.66516 18.53178 22.12281   0.05584025 0.05584025
3 8 15.97995 15.30663 18.06831 20.64181   0.11064299 0.11064299
4 8 16.30548 16.40500 19.17237 20.75895   0.36484699 0.36484699

################################################################################################################################


# Log likelihood by the gamma distribution model
system.time(
hoge <- freqpcr:::.freqpcr_loglike( X=c(P=-2, K=1, targetScale=0, sdMeasure=log(0.24), EPCR=0), N=dat.gamma$N,
                                    DCW=dat.gamma$target0-dat.gamma$housek0, DCD=dat.gamma$target1-dat.gamma$housek1,
                                    zeroAmount=0.0016, para.fixed=NULL, beta=FALSE, diploid=FALSE )
)
hoge


> hoge
[1] 7.049412


# formulation by the beta distribution model
system.time(
hoge <- freqpcr:::.freqpcr_loglike( X=c(P=-2, K=1, targetScale=0, sdMeasure=log(0.24), EPCR=0), N=dat.gamma$N,
                                    DCW=dat.gamma$target0-dat.gamma$housek0, DCD=dat.gamma$target1-dat.gamma$housek1,
                                    zeroAmount=0.0016, para.fixed=NULL, beta=TRUE, diploid=FALSE )
)
hoge


> hoge
[1] 7.049006


################################################################################################################################

(the following are preliminary implementation: not used in the paper)


Since freqpcr version 0.3.2, you can input Cq data with missing values (results are not shown)

#dat.gamma[1, "target1"] <- NA
#dat.gamma[1, "housek0"] <- NA


Since freqpcr version 0.3.2, you can use a "continuous" sample allele-ratio distribution,
in which the environmental allele ratio (p) follows directly a beta/gamma distribution, skipping the binomial sampling process. 
These direct DNA sampling may be consistent with environmental DNA though it has not been verified in field.


# Log likelihood by the gamma distribution model, ignoring binomial sampling (continuous gamma model)
system.time(
hoge <- freqpcr:::.freqpcr_loglike_cont(X=c(P=-2, K=1, targetScale=0, sdMeasure=log(0.24), EPCR=0), A=dat.gamma$N,
                                        DCW=dat.gamma$target0-dat.gamma$housek0, DCD=dat.gamma$target1-dat.gamma$housek1,
                                        zeroAmount=0.0016, para.fixed=NULL, beta=FALSE)
)
hoge


> hoge
[1] 86.18858


# Log likelihood by the beta distribution model, ignoring binomial sampling (continuous beta model)
system.time(
hoge <- freqpcr:::.freqpcr_loglike_cont(X=c(P=-2, K=1, targetScale=0, sdMeasure=log(0.24), EPCR=0), A=dat.gamma$N,
                                        DCW=dat.gamma$target0-dat.gamma$housek0, DCD=dat.gamma$target1-dat.gamma$housek1,
                                        zeroAmount=0.0016, para.fixed=NULL, beta=TRUE)
)
hoge


> hoge
[1] 14.08126


################################################################################################################################

Simultaneous parameter estimation with freqpcr()

# Assmmsing the gamma distributions and all parameters other than EPCR are unknown.

# print.level: 0) for no output during calculation, 1) for first/last iterations, 2) for all iterations

# if you feed some value for each of P, K, targetScale, sdMeasure,
# it is then treated as a fixed parameter value (not estimated).

# If a parameter is unknown, put NULL or simply ignore them.
# However, if you want to treat EPCR as unknown variable, you must explicitly write as "EPCR=NULL"
# or it is treated as a fixed parameter with default (= 0.99).


EPCR <- 0.97
result1 <- freqpcr( N=dat.gamma$N,
                    housek0=dat.gamma$housek0, target0=dat.gamma$target0,
                    housek1=dat.gamma$housek1, target1=dat.gamma$target1,
                    P=NULL, K=NULL, targetScale=NULL, sdMeasure=NULL, EPCR=EPCR,
                    zeroAmount=1.6e-03, beta=FALSE, diploid=FALSE,
                    pvalue=0.05, gradtol=1e-4, steptol=1e-9, iterlim=100, maxtime=600, print.level=1)
result1


> result1
An object of class "CqFreq"
Slot "report":
                                               Estimate Fixed   (scaled) (scaled.SE)       2.5%     97.5%
P (R-allele frequency)                        0.1141011     0 -2.0495184  0.58284394 0.03947276 0.2875803
K (gamma shape parameter)                     3.5678195     0  1.2719546         NaN        NaN       NaN
targetScale (relative amount of target locus) 1.1481927     0  0.1381891         NaN        NaN       NaN
Cq measurement error (SD)                     0.2187155     0 -1.5199833  0.06417172 0.19286641 0.2480291
EPCR (Duplication efficiency of PCR)          0.9700000     1         NA          NA         NA        NA

Slot "obj":
$minimum
[1] 6.808096

$estimate
[1] -2.0495184  1.2719546  0.1381891 -1.5199833

$gradient
[1]  2.704796e-01  7.025499e+04  8.934158e+04 -5.877667e+04

$hessian
              [,1]          [,2]          [,3]          [,4]
[1,]     2.9197547 -1.847117e+03 -2.345098e+03 -1.581763e-01
[2,] -1847.1167828 -1.104395e+09 -1.404740e+09 -2.500117e+04
[3,] -2345.0977397 -1.404740e+09 -1.786766e+09 -3.168260e+04
[4,]    -0.1581763 -2.500117e+04 -3.168260e+04  9.021054e+00

$code
[1] 3

$iterations
[1] 29


Slot "cal.time":
   user  system elapsed 
  11.44    0.65   12.14 


########

# Simultaneous parameter estimation with beta distribution (beta=TRUE). Parameters other than EPCR are set unknown.

EPCR <- 0.97
result1 <- freqpcr( N=dat.gamma$N,
                    housek0=dat.gamma$housek0, target0=dat.gamma$target0,
                    housek1=dat.gamma$housek1, target1=dat.gamma$target1,
                    P=NULL, K=NULL, targetScale=NULL, sdMeasure=NULL, EPCR=EPCR,
                    zeroAmount=1.6e-03, beta=TRUE, diploid=FALSE,
                    pvalue=0.05, gradtol=1e-4, steptol=1e-9, iterlim=100, maxtime=600, print.level=2)
result1


> result1
An object of class "CqFreq"
Slot "report":
                                                 Estimate Fixed  (scaled) (scaled.SE)       2.5%       97.5%
P (R-allele frequency)                         0.09773189     0 -2.222684  0.60914921 0.03178086   0.2633220
K (gamma shape parameter)                     20.92728468     0  3.041054  1.83522498 0.57354375 763.5882087
targetScale (relative amount of target locus)  1.11922896     0  0.112640  0.08911953 0.93985371   1.3328388
Cq measurement error (SD)                      0.20973065     0 -1.561931  0.32845068 0.11017528   0.3992451
EPCR (Duplication efficiency of PCR)           0.97000000     1        NA          NA         NA          NA

Slot "obj":
$minimum
[1] 6.094915

$estimate
[1] -2.222684  3.041054  0.112640 -1.561931

$gradient
[1] -3.400973e-05 -8.275977e-05 -5.170087e-05  8.878366e-05

$hessian
            [,1]        [,2]       [,3]       [,4]
[1,]  2.71023737  0.05094365   1.168535 -0.1766755
[2,]  0.05094365  0.37630061   2.045198 -0.6539722
[3,]  1.16853469  2.04519835 147.389559  6.4578191
[4,] -0.17667547 -0.65397222   6.457819 11.1504638

$code
[1] 1

$iterations
[1] 12


Slot "cal.time":
   user  system elapsed 
   0.73    0.20    0.94 


Faster but K was overestimated.

########

# Simultaneous parameter estimation with beta distribution (beta=TRUE), assuming K=2 (true gamma shape is known).

EPCR <- 0.97
result1 <- freqpcr( N=dat.gamma$N,
                    housek0=dat.gamma$housek0, target0=dat.gamma$target0,
                    housek1=dat.gamma$housek1, target1=dat.gamma$target1,
                    P=NULL, K=4, targetScale=NULL, sdMeasure=NULL, EPCR=EPCR,
                    zeroAmount=1.6e-03, beta=TRUE, diploid=FALSE,
                    pvalue=0.05, gradtol=1e-4, steptol=1e-9, iterlim=100, maxtime=600, print.level=1)
result1


> result1
An object of class "CqFreq"
Slot "report":
                                               Estimate Fixed   (scaled) (scaled.SE)       2.5%     97.5%
P (R-allele frequency)                        0.1035739     0 -2.1581308  0.60872713 0.03385543 0.2758687
K (gamma shape parameter)                     4.0000000     1         NA          NA         NA        NA
targetScale (relative amount of target locus) 1.1400313     0  0.1310557  0.08538882 0.96434774 1.3477206
Cq measurement error (SD)                     0.2025401     0 -1.5968176  0.30962382 0.11039736 0.3715893
EPCR (Duplication efficiency of PCR)          0.9700000     1         NA          NA         NA        NA

Slot "obj":
$minimum
[1] 6.678446

$estimate
[1] -2.1581308  0.1310557 -1.5968176

$gradient
[1] -7.381558e-06 -5.636824e-05  1.196814e-05

$hessian
           [,1]       [,2]       [,3]
[1,]  2.7089129   1.038771 -0.1281927
[2,]  1.0387709 139.379620  4.3497087
[3,] -0.1281927   4.349709 10.5764221

$code
[1] 1

$iterations
[1] 9


Slot "cal.time":
   user  system elapsed 
   0.44    0.13    0.56 


Pretty fast and accurate. If the shape parameter is known, this option is the best.


################################################################################################################################

# case of diploidy

################################################################################################################################


# freqpcr() also deals with diploidy (diploid=TRUE). This is an experimental implementation.
# We assume i.i.d. on the DNA quantities of S and R owned by heterozygotes.
# Both beta=TRUE/FALSE can be used.


system.time(
hoge <- freqpcr:::.freqpcr_loglike( X=c(P=-2, K=1, targetScale=0, sdMeasure=log(0.24), EPCR=0), N=dat2.gamma$N,
                                    DCW=dat2.gamma$target0-dat2.gamma$housek0, DCD=dat2.gamma$target1-dat2.gamma$housek1,
                                    zeroAmount=0.0016, para.fixed=NULL, beta=FALSE, diploid=TRUE )
)
hoge


> hoge
[1] 9.009644


# Simultaneous parameter estimation for diploids
# diploid, assuming i.i.d, use beta distribution, exact size of K is known.

EPCR <- 0.97
result1 <- freqpcr( N=dat2.gamma$N,
                    housek0=dat2.gamma$housek0, target0=dat2.gamma$target0,
                    housek1=dat2.gamma$housek1, target1=dat2.gamma$target1,
                    P=NULL, K=4, targetScale=NULL, sdMeasure=NULL, EPCR=EPCR,
                    zeroAmount=1.6e-03, beta=TRUE, diploid=TRUE,
                    pvalue=0.05, gradtol=1e-4, steptol=1e-9, iterlim=100, maxtime=600, print.level=1)
result1


> result1
An object of class "CqFreq"
Slot "report":
                                               Estimate Fixed   (scaled) (scaled.SE)       2.5%     97.5%
P (R-allele frequency)                        0.1581764     0 -1.6718598   0.4275540 0.07517052 0.3028276
K (gamma shape parameter)                     4.0000000     1         NA          NA         NA        NA
targetScale (relative amount of target locus) 1.2088993     0  0.1897103   0.1209760 0.95370738 1.5323752
Cq measurement error (SD)                     0.2526398     0 -1.3757906   0.3402709 0.12967689 0.4921992
EPCR (Duplication efficiency of PCR)          0.9700000     1         NA          NA         NA        NA

Slot "obj":
$minimum
[1] 6.791649

$estimate
[1] -1.6718598  0.1897103 -1.3757906

$gradient
[1] 0.0001637775 0.0006179155 0.0001780152

$hessian
           [,1]      [,2]       [,3]
[1,]  6.0753247  6.669574 -0.2452993
[2,]  6.6695739 76.380224  2.2547991
[3,] -0.2452993  2.254799  8.7389088

$code
[1] 1

$iterations
[1] 8


Slot "cal.time":
   user  system elapsed 
   0.81    0.13    0.94 


########


# diploid, use gamma distribution and K is known.

# Caution: the combination of beta=FALSE (the gamma model) and diploid=TRUE is very slow!


result1 <- freqpcr( N=dat2.gamma$N,
                    housek0=dat2.gamma$housek0, target0=dat2.gamma$target0,
                    housek1=dat2.gamma$housek1, target1=dat2.gamma$target1,
                    P=NULL, K=4, targetScale=NULL, sdMeasure=NULL, EPCR=0.97,
                    zeroAmount=1.6e-03, beta=FALSE, diploid=TRUE,
                    pvalue=0.05, gradtol=1e-4, steptol=1e-9, iterlim=100, maxtime=600, print.level=2)
result1

 
> result1
An object of class "CqFreq"
Slot "report":
                                               Estimate Fixed   (scaled) (scaled.SE)       2.5%     97.5%
P (R-allele frequency)                        0.1499250     0 -1.7351892   0.4224756 0.07154295 0.2875829
K (gamma shape parameter)                     4.0000000     1         NA          NA         NA        NA
targetScale (relative amount of target locus) 1.2011939     0  0.1833159         NaN        NaN       NaN
Cq measurement error (SD)                     0.2940397     0 -1.2240404   0.2615032 0.17612220 0.4909055
EPCR (Duplication efficiency of PCR)          0.9700000     1         NA          NA         NA        NA

Slot "obj":
$minimum
[1] 6.899955

$estimate
[1] -1.7351892  0.1833159 -1.2240404

$gradient
[1] -4.424957e-01  8.848356e+04 -7.228854e+04

$hessian
              [,1]          [,2]          [,3]
[1,]  5.516096e+00 -1.536161e+04 -2.011408e-01
[2,] -1.536161e+04 -1.769565e+09 -1.188202e+05
[3,] -2.011408e-01 -1.188202e+05  6.766996e+00

$code
[1] 3

$iterations
[1] 36


Slot "cal.time":
   user  system elapsed 
 162.90    1.19  164.25 


################################################################################################################################

# Test the function of parameter estimation from seed

################################################################################################################################

# sim_dummy() adopts the result of make_dummy() and returns the estimates of p and other parameters.
# Note that a CqList object, output of make_dummy(), in no way contains the value of the original parameters like p.


dat.DNA
EPCR <- 0.97
zeroAmount <- 0.0016
result <- sim_dummy(CqList=dat.DNA, EPCR=EPCR, zeroAmount=zeroAmount,
                    K=K, # only if some of the parameters e.g. K is given a priori
                    beta=TRUE, diploid=FALSE, maxtime=60, print.level=1, aux="Beta TRUE")
result


> result
An object of class "CqFreq"
Slot "report":
                                               Estimate Fixed   (scaled) (scaled.SE)       2.5%     97.5%
P (R-allele frequency)                        0.1035739     0 -2.1581308  0.60872713 0.03385543 0.2758687
K (gamma shape parameter)                     4.0000000     1         NA          NA         NA        NA
targetScale (relative amount of target locus) 1.1400313     0  0.1310557  0.08538882 0.96434774 1.3477206
Cq measurement error (SD)                     0.2025401     0 -1.5968176  0.30962382 0.11039736 0.3715893
EPCR (Duplication efficiency of PCR)          0.9700000     1         NA          NA         NA        NA

Slot "obj":
$minimum
[1] 6.678446

$estimate
[1] -2.1581308  0.1310557 -1.5968176

$gradient
[1] -7.381558e-06 -5.636824e-05  1.196814e-05

$hessian
           [,1]       [,2]       [,3]
[1,]  2.7089129   1.038771 -0.1281927
[2,]  1.0387709 139.379620  4.3497087
[3,] -0.1281927   4.349709 10.5764221

$code
[1] 1

$iterations
[1] 9


Slot "cal.time":
   user  system elapsed 
   0.40    0.07    0.49 


# case of diploidy
result <- sim_dummy(CqList=dat2.DNA, EPCR=EPCR, zeroAmount=zeroAmount,
                    K=K, # only if some of the parameters e.g. K is given a priori
                    beta=TRUE, diploid=TRUE, maxtime=60, print.level=1, aux="just a test")
result


# If you specified diploid=TRUE for the dataset generated as haploids...
result <- sim_dummy(CqList=dat.DNA, EPCR=EPCR, zeroAmount=zeroAmount,
                    K=K, # only if some of the parameters e.g. K is given a priori
                    beta=TRUE, diploid=TRUE, maxtime=60, print.level=1, aux="just a test")
# Actually, not so bad.
result


################################################################################################################################
```
