## Supplementary material for "freqpcr: estimation of population allele frequency using qPCR ΔΔCq measures from bulk samples": ESM1: mite Cq data, ESM2: R source for Figure 1, ESM3: simulation code.: ESM3_simulation.R

```
################################################################################################################################

# Name: freqpcr
# Short description: interval estimation of population allele frequency based on \u0394\u0394Cq measures of quantitative PCR over bulk samples
# Author: Masaaki Sudo (NARO, JAPAN)
# Maintainer: Masaaki Sudo
## https://github.com/sudoms/freqpcr
## https://orcid.org/0000-0001-9834-9857


################################################################################################################################
# Part III
# Numerical experiment to evaluate the functions for simultaneous parameter estimation
################################################################################################################################


# For each of the 624 parameter regions, the estimation on a randomly-generated dataset was replicated 1000 times.
# The dummy datasets for the nth replicate were generated using freqpcr::make_dummy() with the RNG seed == n.
# (624 regions) * (five replicates) were simulated in a batch. This is what called below an "iter".
# Therefore, the simulation comprised 200 batches (iterations). Those batches can be run on separate machines.

# Furthermore, the procedure was evaluated in four different configurations.
# 1. Assuming Beta distribution, fixing K = 1 (and P, targetScale, and sdMeasure are unknown).
# 2. Assuming Beta distribution, treating K (and all other parameters) as unknown.
# 3. Assuming Gamma distribution, fixing K = 1
# 4. Assuming Gamma distribution, treating K as unknown.

# As the configurations 3 and 4 requires more time, they were only replicated 250 times (first 50 batches).

# Note: those simulations were conducted on haploidy. The function also has a limited ability to deal with diploids,
# but it has not been evaluated in the paper because it requires considerable machine resource.


library(tidyverse)
library(parallel) # >= R 2.14.0
library(future)
library(furrr) # if not installed, install.packages("furrr")
library(installr) # if not installed, install.packages("installr")
sessionInfo()
library(freqpcr)
options(future.makeNodePSOCK.connectTimeout=30*24*60*60); getOption("future.makeNodePSOCK.connectTimeout");
options(future.makeNodePSOCK.timeout=24*60*60); getOption("future.makeNodePSOCK.timeout");


# Set information for the session.
set.seed(NULL)
trialid <- 172308L # rename it as you like
analysis_name <- paste("200824", trialid, sep=".")
#repl.total <- 2; repl.each <- 1; # for small demo
repl.total <- 1000
repl.each <- 5 # real code: number of replicates run in a single iteration

# The following code execute the first 50 iterations (50 iterations * 5 repl.each = 250 replicates).
# For the first time, put "iter.skip <- 0" and "iter.to <- 50" to execute the first 50 iterations.
# For the second time, put "iter.skip <- 50" and "iter.to <- 100" to execute the next 50 iterations.
iter.skip <- 0 # skip first n iterations (default 0 = no skip)
#iter.to <- 50 # the last iteration actually calculated
iter.to <- 2 # for small demos; calculate only the first two iterations (624 regions * 10 replicates)
iter <- (iter.skip+1):iter.to # the iterator values which are actually used in the session.

P <- c(0.01, 0.05, 0.1, 0.25, 0.5, 0.75)
K <- c(1, 3, 9, 27)
ntrap <- 2^seq(0, 6, by=1) # N: we do not consider ntrap=1 (DNA from all individuals are extracted together)
npertrap <- 2^seq(0, 6, by=1) # n_h: npertrap must be >= 1
diploid <- c(FALSE)

# om: the tibble that stores the parameter combination (and the results).
om <- tidyr::expand_grid(P, K, ntrap, npertrap, replicate=c(1:repl.total), diploid) %>%
    dplyr::mutate(  scaleDNA=(1/K)*1e-06, targetScale=1.2, baseChange=0.24,
                    EPCR=0.97, zeroAmount=1.6e-03, sdMeasure=0.2  ) %>% # regardless of "iter.skip", all rows are generated.
    dplyr::mutate(ntotal=ntrap*npertrap) %>%
    dplyr::filter(ntotal<=128, ntotal>=4, ntrap>=2)
om$LID <- c(1:nrow(om))
set.seed(1729)
om$rseed <- sample(nrow(om), replace=FALSE) # rseed is affected by the size of "repl.total"
set.seed(NULL)
om <- om %>%
    dplyr::filter(replicate > floor(iter.skip*repl.each), replicate <= ceiling(iter.to*repl.each)) %>%
    dplyr::select(LID, everything())
print(om)
print(nrow(om))
print(xtabs( ~ ntotal + ntrap, data=om))

# Generate the dummy data of DNA yield for each bulk sample (this operation returns lists in the tibble).
# This process should not be parallelized due to the consistency of RNG seeds. Calculation time is not proportional:
# 18000 regions -> 17 sec, 78,000 -> 271 sec, 156,000 -> 1668 sec
ptime0 <- proc.time()
om <- om %>%
    dplyr::group_by(LID) %>%
    tidyr::nest() %>%
    dplyr::mutate(  DNA=purrr::map( .x=data,
                                    .f=~make_dummy( rand.seed=.$rseed, P=.$P, K=.$K,
                                                    ntrap=.$ntrap, npertrap=.$npertrap, scaleDNA=.$scaleDNA,
                                                    targetScale=.$targetScale, baseChange=.$baseChange,
                                                    EPCR=.$EPCR, zeroAmount=.$zeroAmount, sdMeasure=.$sdMeasure,
                                                    diploid=.$diploid ) )  ) %>%
    dplyr::mutate(  ave_P_mean=purrr::map_dbl(.x=DNA, .f=~mean(pluck(., "deldel"))),
                    ave_P_sd=purrr::map_dbl(.x=DNA, .f=~sd(pluck(., "deldel")))  ) %>%
    tidyr::unnest(cols=data) %>%
    dplyr::ungroup() %>%
    dplyr::arrange(rseed) %>% # arrange() should comes after mutate(), or calculation time explodes.
    dplyr::mutate(ave_P_est=(1.0+EPCR)^(-ave_P_mean)) # estimation by simple averaging of Cq
cal.time <- proc.time()-ptime0; print(cal.time);
invisible(gc()); invisible(gc());
print(om)


om[1, "DNA"][[1]][[1]]; # The data looks like that


# simulation 1. Beta distribution, K = 1
session <- "Beta_K1" ################################################
param.beta <- TRUE ################################################
param.K <- 1 ################################################


# the following simulation code will be recycled.
for (i in iter) {
    iter.num <- formatC( as.integer(i), width=4, flag="0" )
    cat(paste( "\nStarting iteration No. ", i, " (min:", iter.skip+1, ", max: ", iter.to, ")", " / ",
                ceiling(repl.total/repl.each), "\n\n", sep="" ))
    om.each <- om %>%
        dplyr::filter(replicate > (i-1)*repl.each, replicate <= i*repl.each) %>%
#        dplyr::filter(npertrap==8) %>% # for small demos
#        dplyr::filter(ntrap==2) %>%
        dplyr::mutate(beta=param.beta, Session=session) ################################################
    print(om.each); cat("\n"); print(xtabs( ~ ntrap + npertrap, data=om.each)); cat("\n");
    flush.console()

    fout <- file(paste("o", analysis_name, session, iter.num, "txt", sep="."), "w") # Initialize log output

    cl <- future::makeClusterPSOCK(availableCores()-1, connectTimeout=24*60*60, timeout=24*60*60)
    cl.pid <- purrr::map_int(cl, ~pluck(., "session_info", "process", "pid"))
    cat("Cluster cl was initialized with the following PID:\n"); print(cl.pid);
    invisible(clusterEvalQ(cl, library(freqpcr))) # load the package on cluster
    future::plan(cluster, workers=cl);
    cat(paste("furrr: plan cluster, worker = cl, number of workers = ", nbrOfWorkers(), "\n", sep=""));

    # Estimation with freqpcr()
    ptime0 <- proc.time()
    e1 <- try( {
        cat(paste("Calculation started in ", Sys.time(), "\n", sep="")); flush.console();
        capture.output( temp <- om.each %>%
            dplyr::group_by(LID) %>%
            tidyr::nest() %>%
            furrr::future_map(  .x=.$data,
                                .f=~sim_dummy(  CqList=pluck(.$DNA, 1), EPCR=.$EPCR, zeroAmount=.$zeroAmount,
                                                K=param.K, ################################################
                                                beta=.$beta, diploid=.$diploid,
                                                maxtime=120, print.level=1, # 120
                                                aux=c(  replicate=.$replicate, rseed=.$rseed,
                                                        P=.$P, K=.$K,
                                                        ntrap=.$ntrap, npertrap=.$npertrap, beta=.$beta  )  )  ),
            file=fout, append=FALSE, type=c("output") ) # "NUL" or /dev/null (unix) to supress the output
        cat(paste("Calculation ended in   ", Sys.time(), "\n", sep=""));
    }, silent=FALSE )
    print(proc.time()-ptime0); flush.console();

    if (class(e1)=="try-error") {
        while (class(e1)=="try-error") {
            cat("All 'Rscript.exe' actually exist on your machine:\n"); print(installr::get_Rscript_PID());
            cat("Recognized by R host:\n"); print(purrr::map_int(cl, ~pluck(., "session_info", "process", "pid")));
            cat("freqpcr() failed in some parameter regions. Retry.\n")
            result <- om.each

            close(fout)
            future::plan(sequential)
            try({parallel::stopCluster(cl)})
            installr::kill_pid(cl.pid)
            closeAllConnections()
            gc(); gc();

            fout <- file(paste("o", analysis_name, session, iter.num, "txt", sep="."), "w") # Re-initialize log output
            # use fewer cluster cores if connection errors occurred.
            cl <- future::makeClusterPSOCK(min(8, floor(availableCores()/2)), connectTimeout=24*60*60, timeout=24*60*60)
            cl.pid <- purrr::map_int(cl, ~pluck(., "session_info", "process", "pid"))
            cat("\nCluster cl was re-initialized with the following PID:\n"); print(cl.pid);
            invisible(clusterEvalQ(cl, library(freqpcr))) # load the package on cluster
            future::plan(cluster, workers=cl)

            # Recalculation with freqpcr()
            ptime0 <- proc.time()
            e1 <- try( {
                cat(paste("Re-calculation started in ", Sys.time(), "\n", sep="")); flush.console();
                capture.output( temp <- om.each %>%
                    dplyr::group_by(LID) %>%
                    tidyr::nest() %>%
                    furrr::future_map(  .x=.$data,
                                        .f=~sim_dummy(  CqList=pluck(.$DNA, 1), EPCR=.$EPCR, zeroAmount=.$zeroAmount,
                                                        K=param.K, ################################################
                                                        beta=.$beta, diploid=.$diploid,
                                                        maxtime=120, print.level=1, # 120
                                                        aux=c(  replicate=.$replicate, rseed=.$rseed,
                                                                P=.$P, K=.$K,
                                                                ntrap=.$ntrap, npertrap=.$npertrap, beta=.$beta  )  )  ),
                    file=fout, append=FALSE, type=c("output") ) # "NUL" or /dev/null (unix) to supress the output
                cat(paste("Re-calculation ended in   ", Sys.time(), "\n", sep=""));
            }, silent=FALSE )
            print(proc.time()-ptime0); flush.console();
        }
    }

    # First, close the log file. if fout has not been initialized, this operation returns error.
    try( {
        close(fout); cat("The log object 'fout' was closed successfully. ");
    } )
    future::plan(sequential);
    cat(paste("Switched to plan sequential, number of workers = ", nbrOfWorkers(), "\n", sep=""));
    stopcl <- try( {
        parallel::stopCluster(cl); cat("Cluster cl stopped successfully.\n");
    } )
    # If even a part of cluster nodes has connection failure (when "freqpcr failed."), stopCluster(cl) returns
    # error such as "Error in serialize(data, node$con)"
    # Then you can still see the rest of the non-removable nodes with showConnection(); they must be closed.
    if (class(stopcl)=="try-error") {
        cat("'stopCluster(cl)' failed; force 'kill_pid' and 'closeAllConnections()';\n")
        cat("Available connections on R host:\n"); print(showConnections(all=TRUE));
        installr::kill_pid(cl.pid) # Kill Rscript.exe by PID.
        # You can also kill all processes named "Rscript.exe" but this goes wrong when you are running multiple R.
#        installr::kill_process("Rscript")
        cat("Cluster cl was forced to close.\n")
    }
    closeAllConnections() # close connections except for stdin, stdout, stderr
    rm(stopcl); rm(cl.pid); rm(cl); gc(); gc();

    # recall the rows of "time-over":
    # "maxtime" set above (120 s) was smaller than the official (600 s) in order to reduce idle nodes.
    temp.iter <- unlist(purrr::map(.x=temp, .f=~pluck(., "obj", "iterations", 1)))
    over.LID <- is.na(temp.iter)
    if (sum(over.LID)>0) {
        cat(paste("\n", sum(over.LID), " rows ended in timeover.\n", sep=""))
        fout <- file(paste("o", analysis_name, session, iter.num, "txt", sep="."), "a") # Reopen log output as append mode.
        ptime0 <- proc.time()
        e1 <- try( {
            cat(paste("Re-calculation for timeover started in ", Sys.time(), "\n", sep="")); flush.console();
            capture.output( temp.over <- om.each %>%
                dplyr::filter(over.LID) %>%
                dplyr::group_by(LID) %>%
                tidyr::nest() %>%
                purrr::map( .x=.$data,
                            .f=~sim_dummy(  CqList=pluck(.$DNA, 1), EPCR=.$EPCR, zeroAmount=.$zeroAmount,
                                            K=param.K, ################################################
                                            beta=.$beta, diploid=.$diploid,
                                            maxtime=600, print.level=1, #600
                                            aux=c(  replicate=.$replicate, rseed=.$rseed,
                                                    P=.$P, K=.$K,
                                                    ntrap=.$ntrap, npertrap=.$npertrap, beta=.$beta  )  ) ),
                file=fout, append=TRUE, type=c("output") ) # "NUL" or /dev/null (unix) to supress the output
            cat(paste("Re-calculation for timeover ended in   ", Sys.time(), "\n", sep=" "));
        }, silent=FALSE )
        print(proc.time()-ptime0); flush.console();
        temp[over.LID] <- temp.over
        try( {
            close(fout); cat(paste("The log object 'fout' was closed successfully.", "\n", sep=""));
        } )
    }

    result <- om.each %>%
        dplyr::mutate(res=temp) %>%
        tidyr::hoist(.col=res, Time.obj="cal.time", obj="obj", report="report") %>%
        dplyr::group_by(LID) %>%
        dplyr::mutate(  Time=pluck(Time.obj, 1, 3), Iter=pluck(obj, 1, "iterations", 1)  ) %>%
        dplyr::mutate(
                        P_lwr=pluck(report, 1)[1, 5], P_est=pluck(report, 1)[1, 1], P_upr=pluck(report, 1)[1, 6],
                        K_lwr=pluck(report, 1)[2, 5], K_est=pluck(report, 1)[2, 1], K_upr=pluck(report, 1)[2, 6],
                        T_lwr=pluck(report, 1)[3, 5], T_est=pluck(report, 1)[3, 1], T_upr=pluck(report, 1)[3, 6],
                        C_lwr=pluck(report, 1)[4, 5], C_est=pluck(report, 1)[4, 1], C_upr=pluck(report, 1)[4, 6]
        ) %>%
        dplyr::select(-Time.obj, -obj, -report) %>%
        dplyr::ungroup()
    rm(temp)
    cat("Iteration succeeded.\n\n")
    rm(e1); gc(); gc();

    result <- result %>% arrange(LID)
    result.df <- select(result, -DNA, -starts_with("res"))
    print(summary(result.df)) # $DNA is a list; summary() got weird.
    assign(paste("r", analysis_name, session, iter.num, sep="."), value=result)
    save(result, file=paste("r", analysis_name, session, iter.num, "rda", sep="."))
    readr::write_csv(result.df, paste("r", analysis_name, session, iter.num, "csv", sep="."))
    cat(paste("Writing", paste("r", analysis_name, session, iter.num, "csv", sep="."), "completed.\n", sep=" "))
    flush.console()
    rm(result, result.df)
}


# Estimation with freqpcr() with Core i9-7920X
# (Beta , K=1   :   170 sec / full parameter region x 5 replicates (3120 rows) / 16 threads)
# (Beta , K=NULL:  1200 sec / full parameter region x 5 replicates (3120 rows) / 16 threads) 16.5 h / 50 iters
# (Gamma, K=1   :  1500 sec / full parameter region x 5 replicates (3120 rows) / 16 threads) 16.5 h / 50 iters
# (Gamma, K=NULL:  3500 sec / full parameter region x 5 replicates (3120 rows) / 16 threads)
# (Gamma, K=NULL: 40000 sec / full parameter region x 5 replicates (3120 rows) /  1 thread )


# 2. Beta distribution, all parameters unknown
session <- "Beta_all" ################################################
param.beta <- TRUE ################################################
param.K <- NULL ################################################


for (i in iter) {

    (replicate the code of the session "Beta_all")

}


# 3. Gamma distribution, K = 1
session <- "Gamma_K1" ################################################
param.beta <- FALSE ################################################
param.K <- 1 ################################################


for (i in iter) {

    (replicate the code of the session "Beta_all")

}


# 4. Gamma distribution, all parameters unknown
session <- "Gamma_all" ################################################
param.beta <- FALSE ################################################
param.K <- NULL ################################################


for (i in iter) {

    (replicate the code of the session "Beta_all")

}


################################################################################################################################
# Part IV
# Figures 3-
################################################################################################################################


# First, run the code of Part III. It takes a few MONTHS, depending on your environment.
# The summary of the calculation results are written to a csv file for each chunk.

# You can instead download the results from the Internet (see below).


# Re-import and bind the result csv files.
library(tidyverse)
trialid <- 172308L
analysis_name <- paste("200824", trialid, sep=".")

# Beta distribution model: 200 replicates
iter <- 1:200
iter_ab <- paste(   formatC( as.integer(min(iter)), width=4, flag="0" ),
                    formatC( as.integer(max(iter)), width=4, flag="0" ), sep="_"   )
sessions <- c("Beta_K1", "Beta_all")
for (k in sessions) {
    objects <- paste("r", analysis_name, k, formatC( as.integer(iter), width=4, flag="0" ), sep=".")
    try( {
        for (j in 1:length(iter)) {
            assign(objects[j], readr::read_csv(paste(objects[j], "csv", sep="."),
                                                col_types="iddddildddddddidddlcdidddddddddddd"), envir=.GlobalEnv)
            print(get(objects[j]))
        }
        bind <- dplyr::bind_rows(mget(objects))
        assign(paste("r", analysis_name, k, iter_ab, sep="."), bind)
        save(bind, file=paste("r", analysis_name, k, iter_ab, "rda", sep=".")) # 60 MB of the R binary file.
        readr::write_csv(bind, paste("r", analysis_name, k, iter_ab, "csv", sep=".")) # 210 MB of the combined csv file.
    } )

}

# Gamma distribution model: first 50 replicates
iter <- 1:50
iter_ab <- paste(   formatC( as.integer(min(iter)), width=4, flag="0" ),
                    formatC( as.integer(max(iter)), width=4, flag="0" ), sep="_"   )
sessions <- c("Gamma_all", "Gamma_K1")
for (k in sessions) {
    objects <- paste("r", analysis_name, k, formatC( as.integer(iter), width=4, flag="0" ), sep=".")
    try( {
        for (j in 1:length(iter)) {
            assign(objects[j], readr::read_csv(paste(objects[j], "csv", sep="."),
                                                col_types="iddddildddddddidddlcdidddddddddddd"), envir=.GlobalEnv)
            print(get(objects[j]))
        }
        bind <- dplyr::bind_rows(mget(objects))
        assign(paste("r", analysis_name, k, iter_ab, sep="."), bind)
        save(bind, file=paste("r", analysis_name, k, iter_ab, "rda", sep=".")) # 15 MB of the R binary file.
        readr::write_csv(bind, paste("r", analysis_name, k, iter_ab, "csv", sep=".")) # 50 MB of the combined csv file.
    } )

}
ls()
get(paste("r", analysis_name, sessions[1], iter_ab, sep="."))


################################################################################################################################


# Download the data from
https://figshare.com/collections/freqpcr/5258027
# (links are found on the bottom of the page) 


# load the combined R binary files
library(tidyverse)
trialid <- 172308L
analysis_name <- paste("200824", trialid, sep=".")
load(paste("r", analysis_name, "Beta_K1", "0001_0200", "rda", sep=".")); bind.Beta_K1 <- bind;
load(paste("r", analysis_name, "Beta_all", "0001_0200", "rda", sep=".")); bind.Beta_all <- bind;
load(paste("r", analysis_name, "Gamma_K1", "0001_0050", "rda", sep=".")); bind.Gamma_K1 <- bind;
load(paste("r", analysis_name, "Gamma_all", "0001_0050", "rda", sep=".")); bind.Gamma_all <- bind;
rm(bind)
ls()


# The combined data seem like that.

> bind.Beta_K1
# A tibble: 624,000 x 34
     LID     P     K ntrap npertrap replicate diploid scaleDNA targetScale baseChange  EPCR zeroAmount sdMeasure ntotal  rseed
   <int> <dbl> <dbl> <dbl>    <dbl>     <int> <lgl>      <dbl>       <dbl>      <dbl> <dbl>      <dbl>     <dbl>  <dbl>  <int>
 1     1  0.01     1     2        2         1 FALSE   0.000001         1.2       0.24  0.97     0.0016       0.2      4 365190
 2     2  0.01     1     2        2         2 FALSE   0.000001         1.2       0.24  0.97     0.0016       0.2      4 272934
 3     3  0.01     1     2        2         3 FALSE   0.000001         1.2       0.24  0.97     0.0016       0.2      4 357485
 4     4  0.01     1     2        2         4 FALSE   0.000001         1.2       0.24  0.97     0.0016       0.2      4 220405
 5     5  0.01     1     2        2         5 FALSE   0.000001         1.2       0.24  0.97     0.0016       0.2      4 537430
 6  1001  0.01     1     2        4         1 FALSE   0.000001         1.2       0.24  0.97     0.0016       0.2      8 460547
 7  1002  0.01     1     2        4         2 FALSE   0.000001         1.2       0.24  0.97     0.0016       0.2      8  33196
 8  1003  0.01     1     2        4         3 FALSE   0.000001         1.2       0.24  0.97     0.0016       0.2      8 546004
 9  1004  0.01     1     2        4         4 FALSE   0.000001         1.2       0.24  0.97     0.0016       0.2      8 552619
10  1005  0.01     1     2        4         5 FALSE   0.000001         1.2       0.24  0.97     0.0016       0.2      8 434526
# ... with 623,990 more rows, and 19 more variables: ave_P_mean <dbl>, ave_P_sd <dbl>, ave_P_est <dbl>, beta <lgl>,
#   Session <chr>, Time <dbl>, Iter <int>, P_lwr <dbl>, P_est <dbl>, P_upr <dbl>, K_lwr <dbl>, K_est <dbl>, K_upr <dbl>,
#   T_lwr <dbl>, T_est <dbl>, T_upr <dbl>, C_lwr <dbl>, C_est <dbl>, C_upr <dbl>


########## code for Figure 3--


# Estimation accuracy of p


print(analysis_name)

library(magrittr)
library(extrafont)
library(ggplot2)
library(ggthemes)
library(cowplot)
library(devEMF)


caption <- "est_P"
sessions <- c("Beta_all", "Beta_K1", "Gamma_all", "Gamma_K1")
sessions.label <- c(Beta_all="Beta distribution, all parameters unknown", Beta_K1="Beta distribution, k was fixed 1",
                    Gamma_all="Gamma distribution, all parameters unknown", Gamma_K1="Gamma distribution, k was fixed 1")
for (k in sessions) {
    dat.bind <- paste("bind", k, sep=".") %>% get()
    dat.nest <- dat.bind %>%
        dplyr::arrange(LID) %>%
        dplyr::group_by(P, K, ntrap, npertrap, ntotal, diploid) %>%
        tidyr::nest() %>%
        dplyr::mutate(  zeroci=binom.test(0, as.integer(ntotal))$conf.int[2]  ) %>%
        dplyr::mutate(  used.binom=P>=zeroci,
                        used.three=ntotal>=3.0/P  )
    print(dat.bind)
    print(dat.nest)
    levels_p <- tibble(P=c(0.01, 0.05, 0.1, 0.25, 0.5, 0.75), level=c(0.01, 0.05, 0.1, 0.25, 0.5, 0.75))
    THREE <- c(hsv(0.4, 0, 0.8), hsv(0.4, 0, 1)) # color chart: is the sample size enough
    p <-ggplot(data=dat.bind) +
        geom_rect(  data=dat.nest, aes(xmin=0, xmax=Inf, ymin=0, ymax=1, fill=used.three), color=NA ) +
        geom_hline( yintercept=c(0, 0.5, 1), linetype=1, size=0.1 ) +
        geom_hline( data=levels_p, aes(yintercept=level), linetype=3, color=gray(0.4), size=0.25 ) +
        geom_boxplot(   aes(y=P_upr, x=ntrap, group=npertrap), notch=FALSE,
                        color=hsv(0.13, 0.99, 0.5), fill=hsv(0.13, 0.99, 0.99), outlier.shape=NA, size=0.3,
                        width=1.6/length(unique(dat.bind$npertrap)), fatten=1.5   ) + # fatten: width of median
        geom_boxplot(   aes(y=P_lwr, x=ntrap, group=npertrap), notch=FALSE,
                        color=hsv(0.56, 0.9, 0.4), fill=hsv(0.56, 0.7, 0.95), outlier.shape=NA, size=0.3,
                        width=1.3/length(unique(dat.bind$npertrap)), fatten=1.5   ) +
        geom_boxplot(   aes(y=P_est, x=ntrap, group=npertrap), notch=FALSE,
                        color=hsv(0.5, 0, 0.15), fill=hsv(0.5, 0, 1, 0.95), outlier.shape=NA, size=0.2,
                        width=1/length(unique(dat.bind$npertrap)), fatten=1.5   ) +
        facet_grid(P ~ ntotal,  margins=F, labeller=purrr::partial(label_both, sep=" = ")) +
        theme_classic(base_size=9) +
        scale_fill_manual(values=THREE) +
        theme(  panel.background=element_blank(), plot.background=element_rect(colour=NA, fill=NA),
                plot.margin=unit(c(0.25, 0, 0.25, 0.25), "lines")  ) + # top, right, bottom, left
        theme(  text=element_text(family="Arial"),# plot.title=element_blank(),
                plot.title=element_text(family="Arial", face="bold", size=8, angle=0, hjust=0),
                strip.background=element_blank(),
                strip.text=element_text(family="Arial Narrow", face="bold", size=8, margin=margin(b=2, t=1, l=4, r=4, unit="pt")),
                axis.text.x=element_text(family="Arial Narrow", face="bold", size=7, angle=0, hjust=0.5),
                axis.text.y=element_text(family="Arial Narrow", face="bold", size=7, angle=0, hjust=1)  ) + # hjust=0.5: center
        theme(legend.position="none") +
        scale_x_log10( breaks=c(1, 2, 4, 8, 16, 32, 64, 128), labels=c(1, 2, 4, 8, 16, 32, 64, 128)) +
        scale_y_continuous( limit=c(0, 1), breaks=c(0, 0.25, 0.5, 0.75, 1), labels=c("0", "", "0.5", "", "1")) +
        labs(   title=sessions.label[k], x="Number of bulk DNA (ntrap)",
                y="Estimated p (95% CI Lower | MLE | 95% CI Upper)")
    outer.title <- paste("Fig", caption, k, sep=".")
    assign(paste("ggp", outer.title, sep="."), value=p)
    ggsave(paste(outer.title, ".pdf", sep=""), device=cairo_pdf, plot=p, dpi=300, width=11/2.54, height=9.6/2.54)
    ggsave(paste(outer.title, ".eps", sep=""), device=cairo_pdf, plot=p, dpi=300, width=11/2.54, height=9.6/2.54)
    ggsave(paste(outer.title, ".png", sep=""), type="cairo", plot=p, dpi=300, width=11/2.54, height=9.6/2.54)
    # save as the enhanced metafile
    emf(file=paste(outer.title, ".emf", sep=""), width=11/2.54, height=9.6/2.54)
    print(p)
    dev.off()
}
warnings()

# combine pdf files
library(pdftools)
pdf_combine(paste("Fig", caption, sessions, "pdf", sep="."), output=paste("Fig", caption, "combined", "pdf", sep="."))

# Figure files for submission
file.copy(from="Fig.est_P.Beta_all.eps", to="Figure3.eps", overwrite=TRUE, copy.date=TRUE)
file.copy(from="Fig.est_P.Beta_all.pdf", to="Figure3.pdf", overwrite=TRUE, copy.date=TRUE)
file.copy(from="Fig.est_P.Beta_all.emf", to="Figure3.emf", overwrite=TRUE, copy.date=TRUE)

file.copy(from="Fig.est_P.Beta_K1.eps", to="FigureS3.eps", overwrite=TRUE, copy.date=TRUE)
file.copy(from="Fig.est_P.Beta_K1.pdf", to="FigureS3.pdf", overwrite=TRUE, copy.date=TRUE)
file.copy(from="Fig.est_P.Beta_K1.emf", to="FigureS3.emf", overwrite=TRUE, copy.date=TRUE)

file.copy(from="Fig.est_P.Gamma_all.eps", to="FigureS4.eps", overwrite=TRUE, copy.date=TRUE)
file.copy(from="Fig.est_P.Gamma_all.pdf", to="FigureS4.pdf", overwrite=TRUE, copy.date=TRUE)
file.copy(from="Fig.est_P.Gamma_all.emf", to="FigureS4.emf", overwrite=TRUE, copy.date=TRUE)


################################################################################################################################


# Estimation of P by simple averaging of DeltaDeltaCt values (Figure 4)


caption <- "est_averaging"
sessions <- c("Beta_all")
sessions.label <- c(Beta_all="Estimation by averaging DeltaDeltaCq values")

for (k in sessions) {
    dat.bind <- paste("bind", k, sep=".") %>% get() %>%
        dplyr::mutate(  ave_P_se=ave_P_sd/sqrt(ntrap),
                        ave_P_est_recalc=(1+EPCR)^(-ave_P_mean),
                        ave_P_lwr=(1+EPCR)^(-ave_P_mean-1.96*ave_P_se),
                        ave_P_upr=(1+EPCR)^(-ave_P_mean+1.96*ave_P_se)  )
    dat.nest <- dat.bind %>%
        dplyr::arrange(LID) %>%
        dplyr::group_by(P, K, ntrap, npertrap, ntotal, diploid) %>%
        tidyr::nest() %>%
        dplyr::mutate(  zeroci=binom.test(0, as.integer(ntotal))$conf.int[2]  ) %>%
        dplyr::mutate(  used.binom=P>=zeroci,
                        used.three=ntotal>=3.0/P  )
    levels_p <- tibble(P=c(0.01, 0.05, 0.1, 0.25, 0.5, 0.75), level=c(0.01, 0.05, 0.1, 0.25, 0.5, 0.75))
    THREE <- c(hsv(0.4, 0, 0.8), hsv(0.4, 0, 1)) # color chart if the sample size is enough

    p <-ggplot(dat.bind) +
        geom_rect(  data=dat.nest, aes(xmin=0, xmax=Inf, ymin=0, ymax=1, fill=used.three), color=NA ) +
        geom_hline( yintercept=c(0, 0.5, 1), linetype=1, size=0.1 ) +
        geom_hline( data=levels_p, aes(yintercept=level), linetype=3, color=gray(0.4), size=0.25 ) +
        geom_boxplot(   aes(y=ave_P_upr, x=ntrap, group=npertrap), notch=FALSE,
                        color=hsv(0.13, 0.99, 0.5), fill=hsv(0.13, 0.99, 0.99), outlier.shape=NA, size=0.3,
                        width=1.6/length(unique(dat.bind$npertrap)), fatten=1.5   ) + # fatten: width of median
        geom_boxplot(   aes(y=ave_P_lwr, x=ntrap, group=npertrap), notch=FALSE,
                        color=hsv(0.56, 0.9, 0.4), fill=hsv(0.56, 0.7, 0.95), outlier.shape=NA, size=0.3,
                        width=1.3/length(unique(dat.bind$npertrap)), fatten=1.5   ) +
        geom_boxplot(   aes(y=ave_P_est, x=ntrap, group=npertrap), notch=FALSE,
                        color=hsv(0.5, 0, 0.15), fill=hsv(0.5, 0, 1, 0.95), outlier.shape=NA, size=0.2,
                        width=1/length(unique(dat.bind$npertrap)), fatten=1.5   ) +
        facet_grid(P ~ ntotal,  margins=F, labeller=purrr::partial(label_both, sep=" = ")) +
        theme_classic(base_size=9) +
        scale_fill_manual(values=THREE) +
        theme(  panel.background=element_blank(), plot.background=element_rect(colour=NA, fill=NA),
                plot.margin=unit(c(0.25, 0, 0.25, 0.25), "lines")  ) + # top, right, bottom, left
        theme(  text=element_text(family="Arial"),# plot.title=element_blank(),
                plot.title=element_text(family="Arial", face="bold", size=8, angle=0, hjust=0),
                strip.background=element_blank(),
                strip.text=element_text(family="Arial Narrow", face="bold", size=8, margin=margin(b=2, t=1, l=4, r=4, unit="pt")),
                axis.text.x=element_text(family="Arial Narrow", face="bold", size=7, angle=0, hjust=0.5),
                axis.text.y=element_text(family="Arial Narrow", face="bold", size=7, angle=0, hjust=1)  ) + # hjust=0.5: center
        theme(legend.position="none") +
        scale_x_log10( breaks=c(1, 2, 4, 8, 16, 32, 64, 128), labels=c(1, 2, 4, 8, 16, 32, 64, 128)) +
        scale_y_continuous( limit=c(0, 1.5), breaks=c(0, 0.25, 0.5, 0.75, 1, 1.25, 1.5), 
                            labels=c("0", "", "0.5", "", "1", "", "") ) +
        labs(   title=sessions.label[k], x="Number of bulk DNA (ntrap)",
                y="Estimated p (Mean - 1.96*SE | Mean | Mean + 1.96*SE)")
    outer.title <- paste("Fig", caption, sep=".")
    assign(paste("ggp", outer.title, sep="."), value=p)
    ggsave(paste(outer.title, ".pdf", sep=""), device=cairo_pdf, plot=p, dpi=300, width=11/2.54, height=9.6/2.54)
    ggsave(paste(outer.title, ".eps", sep=""), device=cairo_pdf, plot=p, dpi=300, width=11/2.54, height=9.6/2.54)
    ggsave(paste(outer.title, ".png", sep=""), type="cairo", plot=p, dpi=300, width=11/2.54, height=9.6/2.54)
    # save as the enhanced metafile
    emf(file=paste(outer.title, ".emf", sep=""), width=11/2.54, height=9.6/2.54)
    print(p)
    dev.off()
}
warnings()


# Figure files for submission
file.copy(from="Fig.est_averaging.eps", to="Figure4.eps", overwrite=TRUE, copy.date=TRUE)
file.copy(from="Fig.est_averaging.pdf", to="Figure4.pdf", overwrite=TRUE, copy.date=TRUE)
file.copy(from="Fig.est_averaging.emf", to="Figure4.emf", overwrite=TRUE, copy.date=TRUE)


################################################################################################################################


# Time/iterations to converge with fitqpcr()


caption <- c("time", "iters")
sessions <- c("Beta_all", "Beta_K1", "Gamma_all", "Gamma_K1")
sessions.label <- c(Beta_all="Beta distribution, all parameters unknown", Beta_K1="Beta distribution, k was fixed 1",
                    Gamma_all="Gamma distribution, all parameters unknown", Gamma_K1="Gamma distribution, k was fixed 1")
for (k in sessions) {
    dat.bind <- paste("bind", k, sep=".") %>% get()
    dat.nest <- dat.bind %>%
        dplyr::arrange(LID) %>%
        dplyr::group_by(P, K, ntrap, npertrap, ntotal, diploid) %>%
        tidyr::nest() %>%
        dplyr::mutate(  zeroci=binom.test(0, as.integer(ntotal))$conf.int[2]  ) %>%
        dplyr::mutate(  used.binom=P>=zeroci,
                        used.three=ntotal>=3.0/P  )
#    levels_p <- tibble(P=c(0.01, 0.05, 0.1, 0.25, 0.5, 0.75), level=c(0.01, 0.05, 0.1, 0.25, 0.5, 0.75))
    THREE <- c(hsv(0.4, 0, 0.8), hsv(0.4, 0, 1)) # color chart if the sample size is enough

    p <-ggplot(dat.bind) +
        geom_rect(  data=dat.nest,
                    aes(xmin=0, xmax=Inf, ymin=0, ymax=Inf, fill=used.three), color=NA ) +
        geom_hline( yintercept=c(seq(0.1, 1, by=0.1), seq(1, 10, by=1), seq(10, 60, by=10), seq(60, 600, by=60)),
                    linetype=1, color=gray(0.88), size=0.05) +
        geom_hline( yintercept=c(1, 60, 3600), linetype=1, size=0.25 ) +
        geom_hline( yintercept=c(0.1, 10, 600), linetype=3, size=0.25 ) +
        geom_boxplot(   aes(y=Time, x=ntrap, group=npertrap), notch=FALSE,
                        color=hsv(0.5, 0, 0.15), fill=hsv(0.5, 0, 1, 0.9), size=0.25,
                        outlier.shape=NA, outlier.size=0.05, outlier.color=gray(0.2),
                        width=1.6/length(unique(dat.bind$npertrap)), fatten=1.0   ) +
        facet_grid(P ~ ntotal,  margins=F, labeller=purrr::partial(label_both, sep=" = ")) +
        theme_classic(base_size=9) +
        scale_fill_manual(values=THREE) +
        theme(  panel.background=element_blank(), plot.background=element_rect(colour=NA, fill=NA),
                plot.margin=unit(c(0.25, 0, 0.25, 0.25), "lines")  ) + # top, right, bottom, left
        theme(  text=element_text(family="Arial"),
                plot.title=element_text(family="Arial", face="bold", size=8, angle=0, hjust=0),
                strip.background=element_blank(),
                strip.text=element_text(family="Arial Narrow", face="bold", size=8, margin=margin(b=2, t=1, l=4, r=4, unit="pt")),
                axis.text.x=element_text(family="Arial Narrow", face="bold", size=7, angle=0, hjust=0.5),
                axis.text.y=element_text(family="Arial Narrow", face="bold", size=7, angle=0, hjust=1)  ) + # hjust=0.5: center
        theme(legend.position="none") +
        scale_x_log10( breaks=c(1, 2, 4, 8, 16, 32, 64, 128), labels=c(1, 2, 4, 8, 16, 32, 64, 128)) +
        scale_y_log10( limit=c(0.01, 1200), breaks=c(0.1, 1, 10, 60, 600, 3600),
                                        labels=c("0.1 s", "1 s", "10 s", "1 m", "10 m", "1 h")) +
        labs(title=sessions.label[k], x="Number of bulk DNA (ntrap)", y="Time\n") #
    outer.title <- paste("Fig", caption[1], k, sep=".")
    assign(paste("ggp", outer.title, sep="."), value=p)

    p <-ggplot(dat.bind) +
        geom_rect(  data=filter(dat.nest),
                    aes(xmin=0, xmax=Inf, ymin=-Inf, ymax=Inf, fill=used.three), color=NA ) +
        geom_hline( yintercept=seq(0, 100, by=10), linetype=1, color=gray(0.88), size=0.1) +
        geom_hline(yintercept=c(1, 50, 100), linetype=1, size=0.25) +
        geom_boxplot(   aes(y=Iter, x=ntrap, group=npertrap), notch=FALSE,
                        color=hsv(0.5, 0, 0.15), fill=hsv(0.5, 0, 1, 0.9), size=0.25,
                        outlier.shape=NA, outlier.size=0.05, outlier.color=gray(0.2),
                        width=1.6/length(unique(dat.bind$npertrap)), fatten=1.0   ) +
        facet_grid(P ~ ntotal,  margins=F, labeller=purrr::partial(label_both, sep=" = ")) +
        theme_classic(base_size=9) +
        theme(aspect.ratio=1) + # keep the aspect ratio regardless of margin.
        scale_fill_manual(values=THREE) +
        theme(  panel.background=element_blank(), plot.background=element_rect(colour=NA, fill=NA),
                plot.margin=unit(c(0.25, 0, 0.25, 0.25), "lines")  ) + # top, right, bottom, left
        theme(  text=element_text(family="Arial"),
                plot.title=element_text(family="Arial", face="bold", size=8, angle=0, hjust=0),
                strip.background=element_blank(),
                strip.text=element_text(family="Arial Narrow", face="bold", size=8, margin=margin(b=2, t=1, l=4, r=4, unit="pt")),
                axis.text.x=element_text(family="Arial Narrow", face="bold", size=7, angle=0, hjust=0.5),
                axis.text.y=element_text(family="Arial Narrow", face="bold", size=7, angle=0, hjust=1)  ) + # hjust=0.5: center
        theme(legend.position="none") +
        scale_x_log10( breaks=c(1, 2, 4, 8, 16, 32, 64, 128), labels=c(1, 2, 4, 8, 16, 32, 64, 128)) +
        scale_y_continuous( limit=c(0, 100), breaks=c(0, 50, 100)) +
        labs(title=sessions.label[k], x="Number of bulk DNA (ntrap)", y="Iterations\n") #\n
    outer.title <- paste("Fig", caption[2], k, sep=".")
    assign(paste("ggp", outer.title, sep="."), value=p)

    # Making conbined figure via cowplot
    library(cowplot)
    ps2 <- cowplot::plot_grid(  get(paste("ggp.Fig.time", k, sep=".")),
                                get(paste("ggp.Fig.iters", k, sep=".")),
                                align="v", rel_heights=c(1, 1), scale=c(1, 1), ncol=1,
                                labels=c("A", "B"), label_size=12, hjust=0.05  )
    outer.title <- paste("Fig", paste(caption, collapse="."), k, sep=".")
    ggsave(paste(outer.title, ".eps", sep=""), device=cairo_pdf, plot=ps2, dpi=300, width=11/2.54, height=22.8/2.54)
    ggsave(paste(outer.title, ".pdf", sep=""), device=cairo_pdf, plot=ps2, dpi=300, width=11/2.54, height=22.8/2.54)
    emf(file=paste(outer.title, ".emf", sep=""), width=11/2.54, height=22.8/2.54)
    print(ps2)
    dev.off()

}
warnings()


# combine pdf files
library(pdftools)
pdf_combine(paste("Fig", paste(caption, collapse="."), sessions, "pdf", sep="."),
            output=paste("Fig", paste(caption, collapse="."), "combined", "pdf", sep="."))


# Figure files for submission
file.copy(from="Fig.time.iters.Beta_all.eps", to="FigureS5.eps", overwrite=TRUE, copy.date=TRUE)
file.copy(from="Fig.time.iters.Beta_all.pdf", to="FigureS5.pdf", overwrite=TRUE, copy.date=TRUE)
file.copy(from="Fig.time.iters.Beta_all.emf", to="FigureS5.emf", overwrite=TRUE, copy.date=TRUE)
file.copy(from="Fig.time.iters.Beta_K1.eps", to="FigureS6.eps", overwrite=TRUE, copy.date=TRUE)
file.copy(from="Fig.time.iters.Beta_K1.pdf", to="FigureS6.pdf", overwrite=TRUE, copy.date=TRUE)
file.copy(from="Fig.time.iters.Beta_K1.emf", to="FigureS6.emf", overwrite=TRUE, copy.date=TRUE)
file.copy(from="Fig.time.iters.Gamma_all.eps", to="FigureS7.eps", overwrite=TRUE, copy.date=TRUE)
file.copy(from="Fig.time.iters.Gamma_all.pdf", to="FigureS7.pdf", overwrite=TRUE, copy.date=TRUE)
file.copy(from="Fig.time.iters.Gamma_all.emf", to="FigureS7.emf", overwrite=TRUE, copy.date=TRUE)


################################################################################################################################


# success probability of the simultaneous parameter estimation

bind.Beta_all


> bind.Beta_all
# A tibble: 624,000 x 34
     LID     P     K ntrap npertrap replicate diploid scaleDNA targetScale baseChange  EPCR zeroAmount sdMeasure ntotal  rseed
   <int> <dbl> <dbl> <dbl>    <dbl>     <int> <lgl>      <dbl>       <dbl>      <dbl> <dbl>      <dbl>     <dbl>  <dbl>  <int>
 1     1  0.01     1     2        2         1 FALSE   0.000001         1.2       0.24  0.97     0.0016       0.2      4 365190
 2     2  0.01     1     2        2         2 FALSE   0.000001         1.2       0.24  0.97     0.0016       0.2      4 272934
 3     3  0.01     1     2        2         3 FALSE   0.000001         1.2       0.24  0.97     0.0016       0.2      4 357485
 4     4  0.01     1     2        2         4 FALSE   0.000001         1.2       0.24  0.97     0.0016       0.2      4 220405
 5     5  0.01     1     2        2         5 FALSE   0.000001         1.2       0.24  0.97     0.0016       0.2      4 537430
 6  1001  0.01     1     2        4         1 FALSE   0.000001         1.2       0.24  0.97     0.0016       0.2      8 460547
 7  1002  0.01     1     2        4         2 FALSE   0.000001         1.2       0.24  0.97     0.0016       0.2      8  33196
 8  1003  0.01     1     2        4         3 FALSE   0.000001         1.2       0.24  0.97     0.0016       0.2      8 546004
 9  1004  0.01     1     2        4         4 FALSE   0.000001         1.2       0.24  0.97     0.0016       0.2      8 552619
10  1005  0.01     1     2        4         5 FALSE   0.000001         1.2       0.24  0.97     0.0016       0.2      8 434526
# ... with 623,990 more rows, and 19 more variables: ave_P_mean <dbl>, ave_P_sd <dbl>, ave_P_est <dbl>, beta <lgl>,
#   Session <chr>, Time <dbl>, Iter <int>, P_lwr <dbl>, P_est <dbl>, P_upr <dbl>, K_lwr <dbl>, K_est <dbl>, K_upr <dbl>,
#   T_lwr <dbl>, T_est <dbl>, T_upr <dbl>, C_lwr <dbl>, C_est <dbl>, C_upr <dbl>


# example code of the data processing to calculate success probabilities (no need to run here)

Beta_all.nest <- bind.Beta_all %>%
    dplyr::arrange(LID) %>%
    dplyr::group_by(P, K, ntrap, npertrap, ntotal, diploid) %>%
    tidyr::nest() %>%
    dplyr::mutate(  zeroci=binom.test(0, as.integer(ntotal))$conf.int[2]  ) %>%
    dplyr::mutate(  used.binom=P>=zeroci,
                    used.three=ntotal>=3.0/P,
                    ns.P_lwr=purrr::map_dbl(.x=data, .f=~{sum(!is.na(.$P_lwr))/length(.$P_lwr)}),
                    ns.K_lwr=purrr::map_dbl(.x=data, .f=~{sum(!is.na(.$K_lwr))/length(.$K_lwr)}),
                    ns.T_lwr=purrr::map_dbl(.x=data, .f=~{sum(!is.na(.$T_lwr))/length(.$T_lwr)}),
                    ns.C_lwr=purrr::map_dbl(.x=data, .f=~{sum(!is.na(.$C_lwr))/length(.$C_lwr)})  )
Beta_all.nest
summary(select(Beta_all.nest, -data)) # Mean is the success probability
summary(select(filter(Beta_all.nest, used.three), -data)) # The success probability when ntotal satisfies 3/P


> summary(select(Beta_all.nest, -data)) # Mean is the success probability
       P                K            ntrap       npertrap       diploid            ntotal           zeroci
 Min.   :0.0100   Min.   : 1.0   Min.   : 2   Min.   : 1.000   Mode :logical   Min.   :  4.00   Min.   :0.02841
 1st Qu.:0.0500   1st Qu.: 2.5   1st Qu.: 4   1st Qu.: 2.000   FALSE:624       1st Qu.: 16.00   1st Qu.:0.05601
 Median :0.1750   Median : 6.0   Median : 8   Median : 4.000                   Median : 32.00   Median :0.10888
 Mean   :0.2767   Mean   :10.0   Mean   :14   Mean   : 9.423                   Mean   : 54.15   Mean   :0.16106
 3rd Qu.:0.5000   3rd Qu.:13.5   3rd Qu.:16   3rd Qu.: 8.000                   3rd Qu.: 64.00   3rd Qu.:0.20591
 Max.   :0.7500   Max.   :27.0   Max.   :64   Max.   :64.000                   Max.   :128.00   Max.   :0.60236
 used.binom      used.three         ns.P_lwr         ns.K_lwr         ns.T_lwr         ns.C_lwr
 Mode :logical   Mode :logical   Min.   :0.0000   Min.   :0.0000   Min.   :0.0000   Min.   :0.0000
 FALSE:268       FALSE:224       1st Qu.:0.4407   1st Qu.:0.2767   1st Qu.:0.4273   1st Qu.:0.4355
 TRUE :356       TRUE :400       Median :0.9730   Median :0.7600   Median :0.8725   Median :0.9340
                                 Mean   :0.7061   Mean   :0.5988   Mean   :0.6806   Mean   :0.6964
                                 3rd Qu.:0.9990   3rd Qu.:0.8745   3rd Qu.:0.9790   3rd Qu.:0.9840
                                 Max.   :1.0000   Max.   :0.9980   Max.   :1.0000   Max.   :1.0000
> summary(select(filter(Beta_all.nest, used.three), -data)) # The success probability when ntotal satisfies 3/P
       P                K            ntrap          npertrap      diploid            ntotal           zeroci
 Min.   :0.0500   Min.   : 1.0   Min.   : 2.00   Min.   : 1.00   Mode :logical   Min.   :  4.00   Min.   :0.02841
 1st Qu.:0.1000   1st Qu.: 2.5   1st Qu.: 4.00   1st Qu.: 2.00   FALSE:400       1st Qu.: 32.00   1st Qu.:0.02841
 Median :0.3750   Median : 6.0   Median : 8.00   Median : 4.00                   Median : 64.00   Median :0.05601
 Mean   :0.3905   Mean   :10.0   Mean   :16.32   Mean   :11.31                   Mean   : 66.48   Mean   :0.10602
 3rd Qu.:0.7500   3rd Qu.:13.5   3rd Qu.:16.00   3rd Qu.:16.00                   3rd Qu.:128.00   3rd Qu.:0.10888
 Max.   :0.7500   Max.   :27.0   Max.   :64.00   Max.   :64.00                   Max.   :128.00   Max.   :0.60236
 used.binom      used.three        ns.P_lwr         ns.K_lwr         ns.T_lwr         ns.C_lwr
 Mode :logical   Mode:logical   Min.   :0.0000   Min.   :0.0000   Min.   :0.0000   Min.   :0.0000
 FALSE:44        TRUE:400       1st Qu.:0.9760   1st Qu.:0.7758   1st Qu.:0.8882   1st Qu.:0.9400
 TRUE :356                      Median :0.9965   Median :0.8460   Median :0.9660   Median :0.9735
                                Mean   :0.8434   Mean   :0.7413   Mean   :0.8083   Mean   :0.8271
                                3rd Qu.:1.0000   3rd Qu.:0.9313   3rd Qu.:0.9842   3rd Qu.:0.9890
                                Max.   :1.0000   Max.   :0.9980   Max.   :1.0000   Max.   :1.0000


################################################################################################################################


# Figure S1, S2 estimation success

# Succecc/fail of the estimation process for each parameter
# Note: "success" only matters some result value has been obtained: the result may deviate from its true value.


caption <- "success"
sessions <- c("Beta_all", "Beta_K1", "Gamma_all", "Gamma_K1")
sessions.label <- c(Beta_all="Beta distribution, all parameters unknown", Beta_K1="Beta distribution, k was fixed 1",
                    Gamma_all="Gamma distribution, all parameters unknown", Gamma_K1="Gamma distribution, k was fixed 1")
i <- 1
for (k in sessions) {
    dat.bind <- paste("bind", k, sep=".") %>% get()
    dat.nest <- dat.bind %>%
        dplyr::arrange(LID) %>%
        dplyr::group_by(P, K, ntrap, npertrap, ntotal, diploid) %>%
        tidyr::nest() %>%
        dplyr::mutate(  zeroci=binom.test(0, as.integer(ntotal))$conf.int[2]  ) %>%
        dplyr::mutate(  used.binom=P>=zeroci,
                        used.three=ntotal>=3.0/P,
                        ns.P_lwr=purrr::map_dbl(.x=data, .f=~{sum(!is.na(.$P_lwr))/length(.$P_lwr)}),
                        ns.K_lwr=purrr::map_dbl(.x=data, .f=~{sum(!is.na(.$K_lwr))/length(.$K_lwr)}),
                        ns.T_lwr=purrr::map_dbl(.x=data, .f=~{sum(!is.na(.$T_lwr))/length(.$T_lwr)}),
                        ns.C_lwr=purrr::map_dbl(.x=data, .f=~{sum(!is.na(.$C_lwr))/length(.$C_lwr)})  )
    dat.nest.longer <- dat.nest %>%
        tidyr::pivot_longer( cols=starts_with("ns."),
                             names_to=c("ns", "Parameter"), values_to=c("pSuccess"), names_sep="\\." )

    data <- dat.nest.longer
    data$Parameter <- factor(data$Parameter, levels=c("P_lwr", "K_lwr", "T_lwr", "C_lwr"))
    label_parameter <- c(   P_lwr=expression(italic(p)), K_lwr=expression(italic(k)), 
                            T_lwr=expression(delta[T]), C_lwr=expression(epsilon[c])   )
    levels_p <- tibble(P=c(0.01, 0.05, 0.1, 0.25, 0.5, 0.75), level=c(0.01, 0.05, 0.1, 0.25, 0.5, 0.75))
    RFREQ <- c(hsv(0, 0, 0), hsv(0.27, 0.95, 0.75, i), hsv(0.95, 0.9, 0.9), hsv(0.72, 0.74, 0.95), hsv(0.15, 0.99, 0.8))

    p <-ggplot(data, aes()) +
        geom_vline( xintercept=c(seq(0.01, 0.1, by=0.01), seq(0.1, 1, by=0.1)),
                    linetype=1, color=gray(0.5), size=0.2) +
        geom_vline(xintercept=c(0.01, 0.1, 1), linetype=1, size=0.4) +
        geom_rect(  aes(xmin=0.01, xmax=1, ymin=0, ymax=1), fill=hsv(0.15, 0.01, 1), color="black", size=0.25) +
        geom_rect(  data=dat.nest, aes(xmin=0.01, xmax=3.0/ntotal, ymin=0, ymax=1), color=NA, fill=gray(0.8) ) +
        geom_hline(yintercept=c(0, 0.5, 1), linetype=1, size=0.25) +
        geom_line(aes(y=pSuccess, x=P, group=interaction(K, Parameter), color=Parameter, linetype=Parameter), size=0.4) +
        geom_point(aes(y=pSuccess, x=P, group=interaction(K, Parameter), color=Parameter, shape=Parameter), size=1.05) +
        scale_color_manual(values=RFREQ, name="Parameter", labels=label_parameter) +
        scale_linetype(name="Parameter", labels=label_parameter) +
        scale_shape(name="Parameter", labels=label_parameter) +
        facet_grid(ntrap + npertrap ~ K,  margins=F, labeller=purrr::partial(label_both, sep=" = ")) +
        theme_classic(base_size=9) +
        theme(  panel.background=element_blank(), plot.background=element_rect(colour=NA, fill=NA),
                plot.margin=unit(c(0, 0, 0.25, 0.25), "lines")  ) + # top, right, bottom, left
        theme(  text=element_text(family="Arial"),
                plot.title=element_text(family="Arial", face="bold", size=8, angle=0, hjust=0),
                strip.background=element_blank(),
                strip.text.x=element_text(  family="Arial Narrow", face="bold", size=8,
                                            margin=margin(b=2, t=2, l=4, r=0, unit="pt")  ),
                strip.text.y=element_text(   family="Arial Narrow", face="bold", size=8, angle=0, hjust=0  ),
                axis.text.x=element_text(family="Arial Narrow", face="bold", size=7, angle=0, hjust=0.5),
                axis.text.y=element_text(family="Arial Narrow", face="bold", size=7, angle=0, hjust=1)  ) + # hjust=0.5: center
        theme(  legend.position="top", legend.spacing=unit(0, "lines"),
                legend.margin=margin(b=0, t=0, l=8, r=4, unit="pt"), # the margin around each legend
                legend.box="vertical",
                legend.box.margin=margin(b=0, t=0, l=0, r=0, unit="pt"), # margins around the full legend area
                legend.box.background=element_blank(),
                legend.box.spacing=unit(4, unit="pt"),
                legend.key.size=unit(1.5, "lines"), legend.key.height=unit(1, "pt"), # enlarge key but not increase margin
                legend.title=element_text(family="Arial", size=8),
                legend.text=element_text(family="Arial", size=8)  ) +
        scale_x_log10(  limit=c(0.01, 1), breaks=c(0.01, 0.1, 0.5, 1), labels=c(0.01, 0.1, 0.5, 1)  ) +
        scale_y_continuous( expand=c(0.1, 0), limit=c(0, 1), breaks=c(0, 0.5, 1) ) +
        labs(title=sessions.label[k], x="True size of p (R allele frequency)", y="Success probability of interval estimation")

    outer.title <- paste("Fig", caption, k, sep=".")
    assign(paste("ggp", outer.title, sep="."), value=p)
    ggsave(paste(outer.title, ".pdf", sep=""), device=cairo_pdf, plot=p, dpi=300, width=15/2.54, height=22.8/2.54)
    ggsave(paste(outer.title, ".eps", sep=""), device=cairo_pdf, plot=p, dpi=300, width=15/2.54, height=22.8/2.54)
    ggsave(paste(outer.title, ".png", sep=""), type="cairo", plot=p, dpi=300, width=15/2.54, height=22.8/2.54)

    # save as the enhanced metafile
    emf(file=paste(outer.title, ".emf", sep=""), width=15/2.54, height=22.8/2.54)
    print(p)
    dev.off()

    i <- (i+1)%%2
}
warnings()


# combine pdf files
library(pdftools)
pdf_combine(paste("Fig", caption, sessions, "pdf", sep="."), output=paste("Fig", caption, "combined", "pdf", sep="."))


# Figure files for submission
file.copy(from="Fig.success.Beta_all.eps", to="FigureS1.eps", overwrite=TRUE, copy.date=TRUE)
file.copy(from="Fig.success.Beta_all.pdf", to="FigureS1.pdf", overwrite=TRUE, copy.date=TRUE)
file.copy(from="Fig.success.Beta_all.emf", to="FigureS1.emf", overwrite=TRUE, copy.date=TRUE)
file.copy(from="Fig.success.Gamma_all.eps", to="FigureS2.eps", overwrite=TRUE, copy.date=TRUE)
file.copy(from="Fig.success.Gamma_all.pdf", to="FigureS2.pdf", overwrite=TRUE, copy.date=TRUE)
file.copy(from="Fig.success.Gamma_all.emf", to="FigureS2.emf", overwrite=TRUE, copy.date=TRUE)


################################################################################################################################


# Estimation accuracy of K (Figure S8)


caption <- "est_K"
sessions.label <- c(Beta_all="A: Estimation of k assuming Beta(mk, (n-m)k)",
                    Gamma_all="B: Estimation of k assuming gamma distributions")
sessions <- c("Beta_all", "Gamma_all")
for (k in sessions) {
    dat.bind <- paste("bind", k, sep=".") %>% get() %>%
        dplyr::filter(ntotal %in% c(128))
    p <-ggplot(dat.bind) +
        geom_hline( yintercept=c(seq(0.01, 0.1, by=0.01), seq(0.1, 1, by=0.1), seq(1, 10, by=1), seq(10, 100, by=10)),
                    linetype=1, color=gray(0.75), size=0.05) +
        geom_hline( yintercept=c(1), linetype=1, size=0.3 ) +
        geom_boxplot(   aes(y=K_upr/K, x=K, group=K), notch=FALSE,
                        color=hsv(0.13, 0.99, 0.5), fill=hsv(0.13, 0.99, 0.99), outlier.shape=NA, size=0.3,
                        width=1.6/length(unique(dat.bind$npertrap)), fatten=1.5   ) + # fatten: width of median
        geom_boxplot(   aes(y=K_lwr/K, x=K, group=K), notch=FALSE,
                        color=hsv(0.56, 0.9, 0.4), fill=hsv(0.56, 0.7, 0.95), outlier.shape=NA, size=0.3,
                        width=1.3/length(unique(dat.bind$npertrap)), fatten=1.5   ) +
        geom_boxplot(   aes(y=K_est/K, x=K, group=K), notch=FALSE,
                        color=hsv(0.5, 0, 0.15), fill=hsv(0.5, 0, 1, 0.95), outlier.shape=NA, size=0.2,
                        width=1/length(unique(dat.bind$npertrap)), fatten=1.5   ) +
        facet_grid(P ~ ntotal+ntrap, margins=F, labeller=purrr::partial(label_both, sep=" = ")) +
        theme_classic(base_size=9) +
        theme(  panel.background=element_blank(), plot.background=element_rect(colour=NA, fill=NA),
                plot.margin=unit(c(0.25, 0.25, 0.25, 0.25), "lines")  ) + # top, right, bottom, left
        theme(  text=element_text(family="Arial"),# plot.title=element_blank(),
                plot.title=element_text(family="Arial", face="bold", size=8, angle=0, hjust=0),
                strip.background=element_blank(),
                strip.text=element_text(family="Arial Narrow", face="bold", size=8, margin=margin(b=2, t=1, l=4, r=4, unit="pt")),
                axis.text.x=element_text(family="Arial Narrow", face="bold", size=7, angle=0, hjust=0.5),
                axis.text.y=element_text(family="Arial Narrow", face="bold", size=7, angle=0, hjust=1)  ) + # hjust=0.5: center
        theme(legend.position="none") +
        scale_x_log10( breaks=c(1, 3, 9, 27), labels=c(1, 3, 9, 27)) +
        scale_y_log10( limit=c(0.001, 1000), breaks=c(0.01, 0.1, 1, 10, 100), labels=c(0.01, 0.1, 1, 10, 100)) +
        labs(title=sessions.label[k], x="Real size of k", y="Estimated k / Real k")
    outer.title <- paste("Fig", caption, k, sep=".")
    assign(paste("ggp", outer.title, sep="."), value=p)
    ggsave(paste(outer.title, ".pdf", sep=""), device=cairo_pdf, plot=p, dpi=300, width=11/2.54, height=12/2.54)
    ggsave(paste(outer.title, ".eps", sep=""), device=cairo_pdf, plot=p, dpi=300, width=11/2.54, height=12/2.54)
    ggsave(paste(outer.title, ".png", sep=""), type="cairo", plot=p, dpi=300, width=11/2.54, height=12/2.54)
    # save as the enhanced metafile
    emf(file=paste(outer.title, ".emf", sep=""), width=11/2.54, height=12/2.54)
    print(p)
    dev.off()
}
warnings()


# combine pdf files
library(pdftools)
pdf_combine(paste("Fig", caption, sessions, "pdf", sep="."), output=paste("Fig", caption, "combined", "pdf", sep="."))


# Making compound figure via cowplot
library(cowplot)
ps2 <- cowplot::plot_grid(  get(paste("ggp.Fig.est_K", sessions[1], sep=".")),
                            get(paste("ggp.Fig.est_K", sessions[2], sep=".")),
                            align="v", rel_heights=c(1, 1), scale=c(1, 1), ncol=1  )
outer.title <- "Fig.est_K"
ggsave(paste(outer.title, ".pdf", sep=""), device=cairo_pdf, plot=ps2, dpi=300, width=11.4/2.54, height=24/2.54)
ggsave(paste(outer.title, ".eps", sep=""), device=cairo_pdf, plot=p, dpi=300, width=11.4/2.54, height=24/2.54)
ggsave(paste(outer.title, ".png", sep=""), type="cairo", plot=p, dpi=300, width=11.4/2.54, height=24/2.54)

emf(file=paste(outer.title, ".emf", sep=""), width=11.4/2.54, height=24/2.54)
print(ps2)
dev.off()


# Figure files for submission
file.copy(from="Fig.est_K.eps", to="FigureS8.eps", overwrite=TRUE, copy.date=TRUE)
file.copy(from="Fig.est_K.pdf", to="FigureS8.pdf", overwrite=TRUE, copy.date=TRUE)
file.copy(from="Fig.est_K.emf", to="FigureS8.emf", overwrite=TRUE, copy.date=TRUE)


################################################################################################################################
```
